# Covalent DNA-Encoded Library Workflow Drives Discovery of SARS-CoV-2 Non-structural Proteins Inhibitors

**DOI:** 10.1101/2024.05.08.593105

**Authors:** Xudong Wang, Liwei Xiong, Ying Zhu, Sixiu Liu, Wenfeng Zhao, Seydimemet Mengnisa, Linjie Li, Xian Lin, Jiaxiang Liu, Xuan Wang, Zhiqiang Duan, Weiwei Lu, Yanrui Suo, Xinyuan Wu, Mengqing Cui, Jinfeng Yue, Rui Jin, Yechun Xu, Lianghe Mei, Hangchen Hu, Xiaojie Lu

## Abstract

The global coronavirus disease 2019 (COVID-19) pandemic persists, with the ongoing mutation of the virus. Consequently, the development of inhibitors with diverse binding modes and mechanisms of action, along with the elucidation of novel binding sites is of paramount importance. The 3-chymotrypsin-like protease (3CLpro) and papain-like protease (PLpro) are two validated cysteine proteases that cleave the viral polyprotein and are essential for viral replication. In this study, we utilized covalent DNA-Encoded libraries (CoDELs) workflow to identify two series of triazine-based covalent inhibitors targeting 3CLpro and PLpro. Molecular docking facilitated the identification of optimization pathways, further refined through medicinal chemistry efforts, leading to the development of the non-peptide 3CLpro inhibitor LU9, which exhibited an IC50 value of 0.34 μM, and crystal structure of LU10 revealed a unique binding mode within the active site. Additionally, the X-ray cocrystal structure of SARS-CoV-2 PLpro with XD5 uncovered a previously unexplored binding site, adjacent to the catalytic pocket, providing an opportunity for further development of PLpro inhibitors. Overall, these novel compounds serve as valuable chemical probes for target validation and represent promising drug candidates for the continued development of SARS-CoV-2 antivirals.

## Introduction

The novel coronavirus was first identified in late 2019, subsequently rapidly spreading to numerous countries and regions, posing a severe threat to public health due to its high fatality rate and complex clinical manifestations.^1^ While vaccines and therapeutic antibodies have demonstrated efficacy in preventing and mitigating COVID-19, their effectiveness is impeded by the emergence of certain variants of concern (VOC), such as the Omicron variant.^2–7^

SARS-CoV-2 is an enveloped, positive-sense, single-stranded RNA virus (**Figure 1A**).^8–10^ The genome sequence exhibits approximately 86% and 50% similarity to those of SARS-CoV and MERS-CoV, two other members of the betacoronavirus family that have been responsible for previous major outbreaks.^11–13^ After the entry of the SARS-CoV-2 into the host cell, the nuclear envelope and the positive-strand RNA of the virus are released into the cytoplasm. The open reading frames ORF1a and ORF1b of the positive-strand RNA are translated to generate non-structural proteins (Nsps), which are further cleaved by papain-like protease (PL^pro^) and 3-chymotrypsin-like protease (3CL^pro^). This process ultimately produces 16 types of non-structural proteins, including helicase and RNA-dependent RNA polymerase (RdRp), which participate in viral transcription and replication processes. (**Figure 1B**).^14–16^ 3CL^pro^ exhibits a tripartit architecture comprising three distinct structural domains (**Figure 1C**). Within the well-defined cleft that interconnects structural domain I and structural domain II, a catalytic dyad, featuring Cys145 and His41 residues, is prominently situated. This catalytic dyad serves as the key determinant responsible for the precise cleavage of pp1a and pp1ab polyproteins, consequently engendering the generation of 12 indispensable functional proteins (Nsp 4–16).^17–19^ PL^pro^, a 35-kDa domain in the 215-kDa multidomain protein Nsp3, is involved in cleaving viral polyproteins pp1a and pp1ab at three sites, generating Nsp1, Nsp2, and Nsp3. It consists of thumb, fingers, and palm subdomains typical of ubiquitin-specific proteases, along with an N-terminal ubiquitin-like domain for substrate recognition (**Figure 1D**).^20–22^ The catalytic triad comprising Cys111, His272, and Asp286 is located at the thumb-palm interface, responsible for its enzymatic activity.

**Figure 1.**
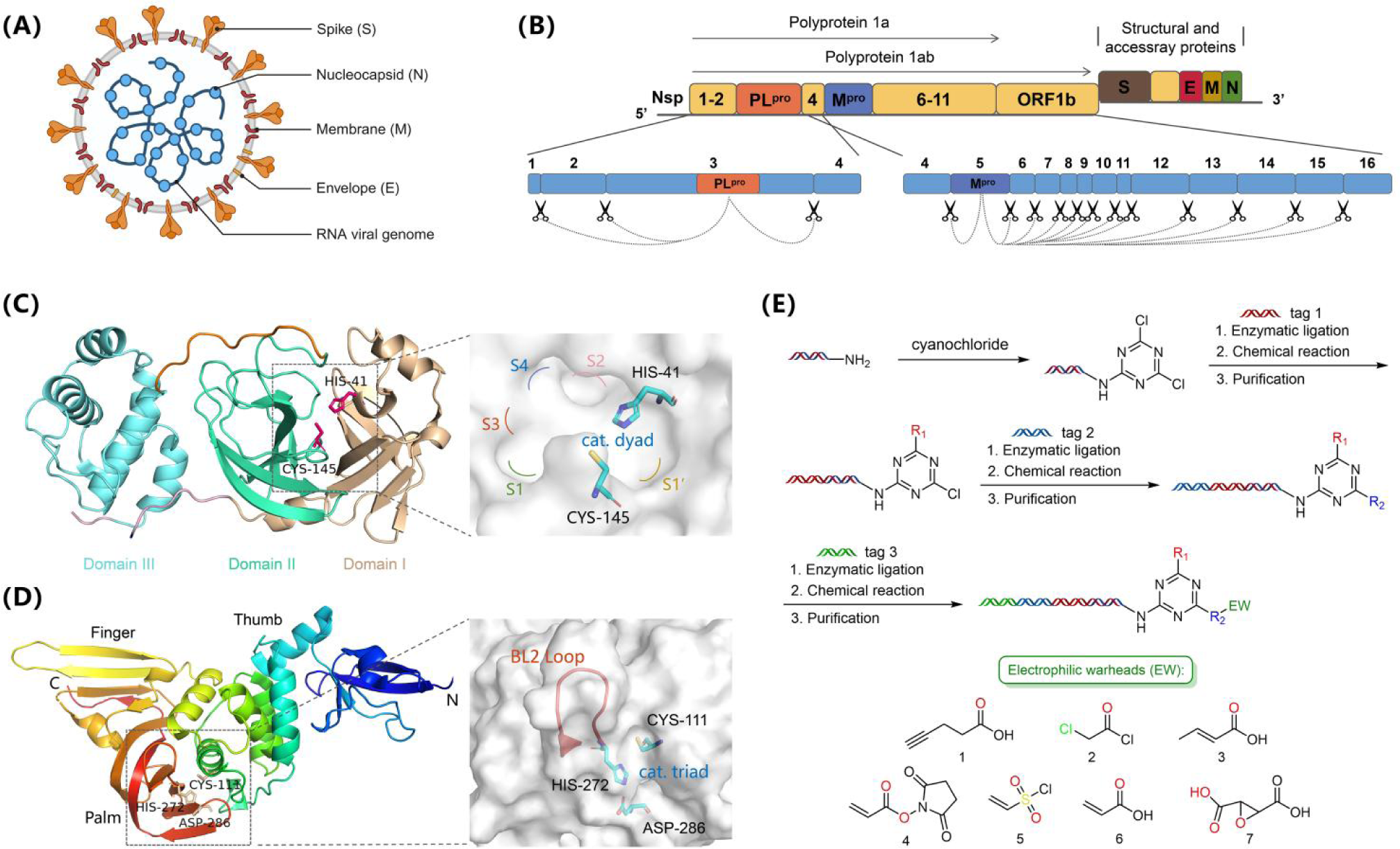
(A) Morphological structure and composition of SARS-CoV-2 coronavirus particle. (B) SARS-CoV-2 genes and their expression proteins. (C) Structural diagrams of 3CL^pro^ and its catalytic pocket. (D) Structural diagrams of PL^pro^ and its catalytic site. (E) Construction route of triazine-based CoDEL.

Multiple peptide-like covalent inhibitors targeting 3CL^pro^ have been extensively investigated, demonstrating significant 3CL^pro^ inhibitory potency and antiviral activity.^23–28^ Nevertheless, these peptidomimetic inhibitors exhibit inherent drawbacks concerning low membrane permeability and metabolic stability. An illustrative example is the approved drug paxlovid (PF-07321332) from Pfizer, which necessitates co-administration with ritonavir as a pharmacokinetic booster to address these limitations.^29^ Presently, the investigation of PL^pro^ binding pockets primarily revolves around those associated with its hydrolytic substrates. Numerous research studies have been dedicated to identifying inhibitors that target this active pocket, with particular emphasis on **GRL0617** and its analogs.^30–34^ Notably, among these investigations, the incorporation of linkers and electrophilic moieties into **GRL0617** led to the development of the first covalent inhibitor, which boasts a well-defined crystal structure.^35^ However, inhibitors designed against SARS-CoV-2 PL^pro^ demonstrate relatively uniform structural features, primarily concentrating on the catalytic pocket. Furthermore, the scarcity of reported crystallographic structures impedes a comprehensive understanding of the protein’s architecture.

In this paper, we present the inaugural application of a triazine-based covalent DNA-encoded library for the identification of structurally unique covalent inhibitors targeting SARS-CoV-2 3CLpro and PLpro (Figure 1E). X-ray crystallography elucidates the binding modes of hit molecules obtained through CoDEL screening. Notably, crystallographic analyses of non-peptidic covalent molecules reveal binding hotspots and potential avenues for further optimization in the context of the 3CLpro target. In addition, for the PLpro target, crystallographic insights unveil a previously unidentified binding site adjacent to the Cys270 residue, offering novel prospects and directions for the discovery of PLpro inhibitors. Most importantly, our research offers a clear research framework for the discovery of other covalent inhibitors targeting non-structural proteins.

## Result and discussion

### Covalent screen of triazine-based DNA encoded libraries for PL^pro^/3CL^pro^ inhibitors

Novel covalent inhibitors were identified through the screening of covalent DNA encoded libraries (CoDELs) based on the triazine scaffold (**Figure 1D**). Within this framework, trichlorotriazine serves as the trifunctional group scaffold, widely employed in library construction and DEL screening to derive a series of lead compounds.^36–39^ In the pursuit of targets 3CL^pro^ and PL^pro^, we initially employed a diverse set of non-covalent DELs for screening. However, the desired outcomes were not attained (**Table S6**). Consequently, through the innovative introduction of electrophilic warheads into reversible compound libraries, we established two covalent compound libraries, namely DEL0C1 and DEL0C2 (**Figure 1D, Figure S1**). Leveraging these covalent compound libraries, collectively referred to as CoDEL, we achieved successful identification of hit compounds targeting 3CL^pro^ and PL^pro^. To assess the screening outcomes, it is imperative to establish cutoff values for both libraries to mitigate background noise interference.^40^ Focusing on the PL^pro^ target, we set the copy value threshold for both libraries to exceed 5, with an enrichment criterion surpassing 120 (**Figure S3**). The findings reveal that DEL0C1 and DEL0C2 demonstrate selectivity solely towards the 5-aminoisoquinoline fragment in cycle 1. Concerning the fragments employed in the libraries construction process during cycle 2, both libraries demonstrate differential levels of enrichment towards various types of diamine structures (**Table S1, S2, S7**). This observation implies that, during the actual covalent binding process, the cycle 2 segment functions solely as a linker in ligand association. Upon analyzing the enrichment results, it is evident that the L9 and B8 fragment (**Table S7**) exhibits the highest degree of enrichment in the screening of both libraries. Specifically, the Select E values for L9 and B8 in DEL0C1 and DEL0C2 are 245 and 208, respectively. Furthermore, regarding the electrophilic fragment in cycle 3, both DEL0C1 and DEL0C2 screening results demonstrate specificity towards chloroacetamide (**Figure 2A**).

**Figure 2.**
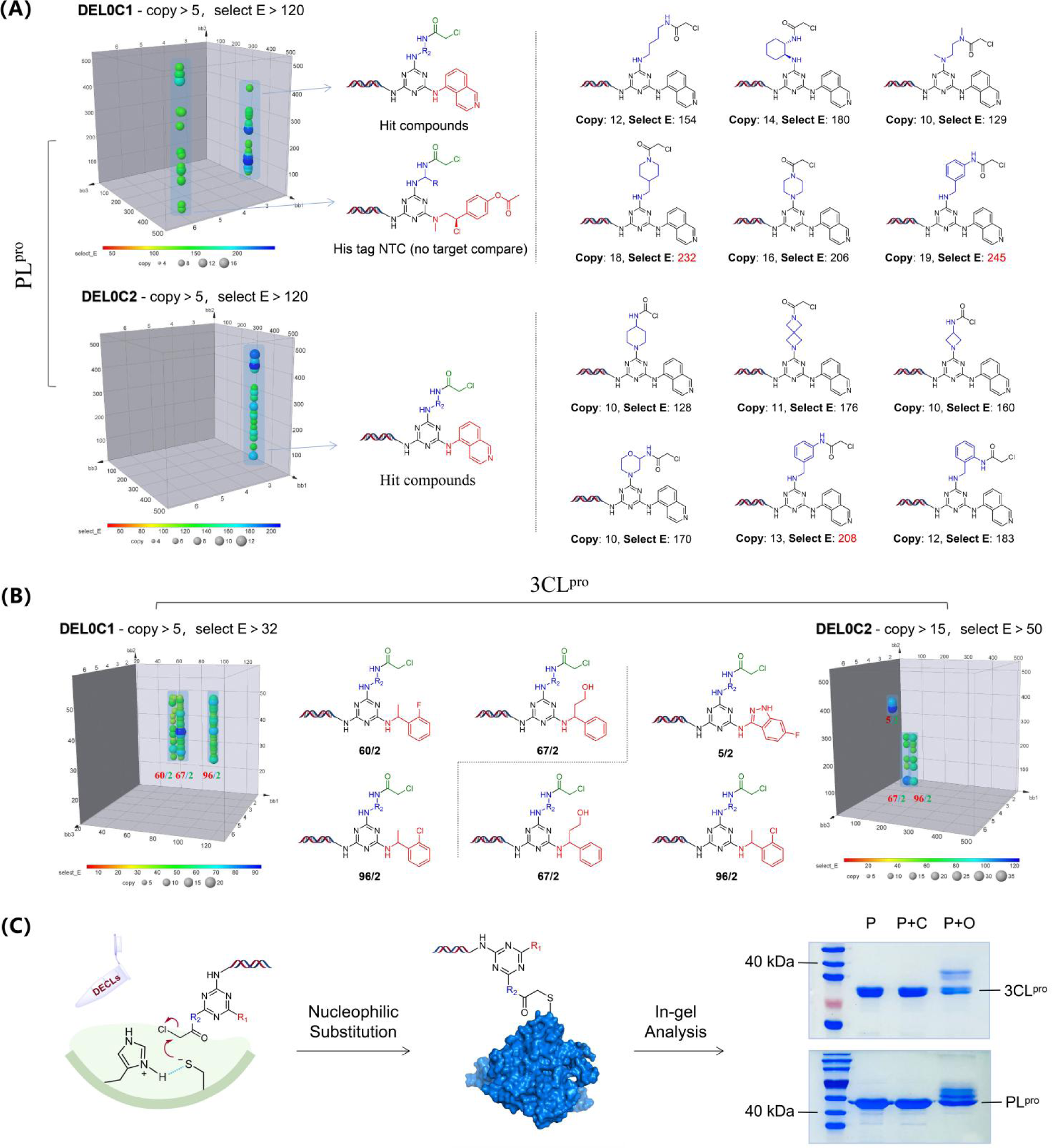
Covalent screening profile of DEL0C1 and DEL0C2 against SARS-CoV-2 PL^pro^ (A) and 3CL^pro^ (B). The selection data have shown the enrichment, copy value and linear relationship. The copy value in the box represents the number of observed library members. (C) Mechanism of covalent binding and the workflow of covalent verification. SDS-PAGE analysis stained with Coomassie Brilliant Blue showing protein bands. P represents protein bands, P+C represents protein bands after incubation with on-DNA control compound, and P+O represents protein bands after incubation with on-DNA covalent compounds.

Regarding the 3CL target, while its screening results similarly exhibit linear enrichment in the three-dimensional plot (**Figure 2B**), these linearly related points represent identical reaction building block. This occurrence stems from the fact that during library construction, each reaction building block in cycle 2 is encoded by multiple distinct tags (**Table S1, Table S2**). From the encoded structural information, it is evident that these identical building blocks all feature piperazine structures. This observation suggests the critical role of the piperazine ring in maintaining conformation during the covalent binding process. Experimental results validating this hypothesis were obtained through ring-opening of the linker and scaffold hoping (**Figure S10**). The screening results from Cycle 1 reveal enrichment towards a class of structurally similar building blocks containing benzylamine moieties (**Figure 2B**). Furthermore, these structures exhibit versatility at the benzyl position, allowing for diverse substitutions or derivatizations, such as methyl introduction, hydroxylation, and cyclization. These outcomes offer clear directions for subsequent derivatization efforts. For the electrophilic segment in cycle 3, screening results from DEL0C1 and DEL0C2 align with those of the PL^pro^ target screening, indicating specificity towards chloroacetamide.

Compared to conventional DEL affinity screening,^41–43^ covalent screening involves repetitive denaturation and elution steps during the screening process to ensure the subsequent identification of covalently bound molecules. However, in cases of high ligand-protein binding affinity, this process may not confer unequivocal assurance. Therefore, covalent binding validation is imperative to ascertain the occurrence of covalent interactions among screened molecules and to precisely identify the covalent binding sites. Our previous research findings indicate that PL^pro^ Cys270 residue represents a potential covalent binding residue,^44,45^ marking the first report of a small molecule inhibitor covalently targeting residue Cys270. Hence, our primary objective concerning covalent molecules derived from CoDEL screening is to delineate their covalent binding sites. To achieve this, we conducted a comparative screening using the PL C111S mutant protein to screen DEL0C1 and DEL0C2. The screening outcomes revealed no discernible enrichment of molecules in either library upon the introduction of the catalytic site cysteine mutation (**Figure S4**). This suggests that the site of covalent binding between molecules in CoDEL and the wild-type PL^pro^ is the Cys111 residue. Subsequently, we employed gel electrophoresis for on-DNA covalent binding validation of these two coronaviral polyproteins. Initially, covalent molecules carrying DNA chains and on-DNA control compound were synthesized (**Scheme S1, S2, S3**), and the procedure outlined in **Figure 2C** was utilized for in-gel analysis of the PL^pro^ and 3CL^pro^ targets. SDS-PAGE analysis demonstrated that hit molecules directly obtained from screening exhibited covalent binding activity.

### Hits Resynthesis and Evaluation of 3CL^pro^ Inhibition

Based on the analysis of DEL screening results, it is evident that positions R1 and R2 emerge as potential sites for modification or derivatization. Consequently, our investigation initially focused on positions R1 and R2, leading to the synthesis of compound **LU1**-**LU7** (**Figure 3A**). Notably, compounds **LU4**, **LU5**, and **LU7** were directly identified through screening, with **LU7** displaying significant enrichment in both libraries (**Figure 3A, Figure S5**). We conducted separate tests to evaluate the inhibitory effects of these seven compounds on the activity of SARS-CoV-2 3CL^pro^ at concentrations of 1 μM and 10 μM. Our results revealed that both **LU5** and **LU7** exhibited significant inhibition at 1 μM. Considering the superior enrichment of **LU7** indicated by DEL affinity screening, implying its stronger binding affinity to the protein, we proceeded to further investigate **LU7** as our primary compound of interest.

**Figure 3.**
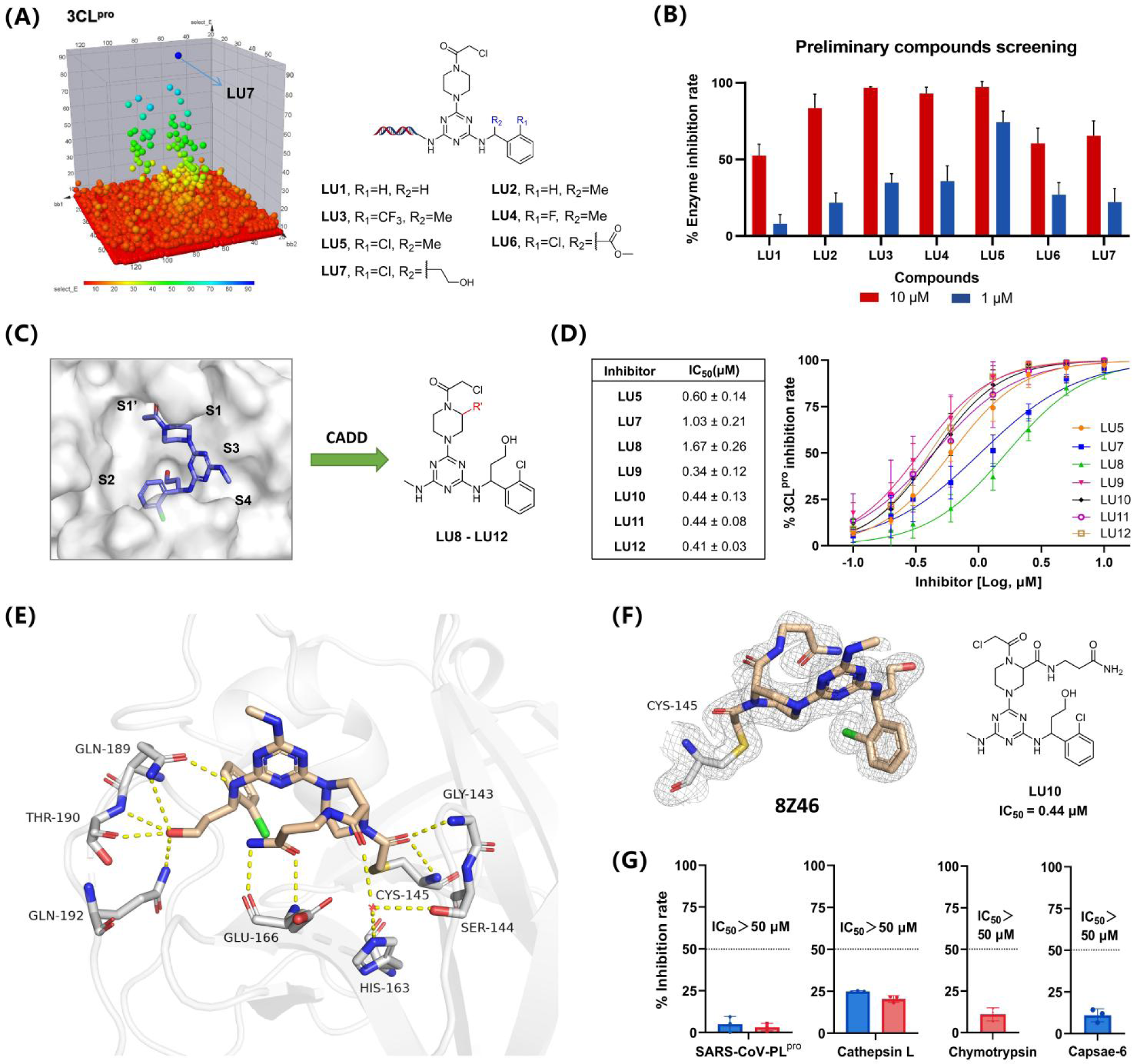
(A) The structure modification guided by DEL resulted in compounds **LU1**-**LU7**. (B) Inhibition rates of **LU1**-**LU7** on 3CL^pro^ enzyme at concentrations of 1μM and 10μM, respectively. (C) Compounds **LU8**-**LU12** obtained via computer-assisted drug design (SBDD) through the modification of compound **LU7**. (D) Dose-response curves for the inhibition of SARS-CoV2 3CL^pro^. The table indicates the corresponding IC_50_ values. (E) X-ray crystal structure of SARS-CoV-2 3CL^pro^ in complex with **LU10** (parchment). (F) Fo − Fc electron density map, show in gray, is contoured at 1σ. (G) Target selectivity of SARS-CoV-2 3CL^pro^ inhibitors against host proteases. Red bars indicates 20 μM, while blue bars indicates 50 μM.

As proteins rich in cysteine, the majority of cysteine residues are situated internally within the protein structure, with only Cys145 and Cys44 potentially exposed as reactive cysteines at the catalytic pocket (**Figure S6**). However, the reported covalent inhibitors of 3CL^pro^ predominantly target the catalytic Cys145.^46–50^ Consequently, we conducted covalent molecular docking of compound **LU7** with this binding site (**Figure 3C**). The covalent docking results reveal the existence of two potential binding modes, wherein a conformational flip of the small molecule occurs contingent upon the positioning of the methylamine moiety (**Figure S7A, S7B**). These dual outcomes afford a comprehensive elucidation concerning the binding of both small molecules and proteins. Nevertheless, DEL screening unequivocally establishes that the DNA tethering site must reside within the solvent-exposed region to enable conjugation. Consequently, the covalent docking outcomes, wherein the methylamine moiety extends toward the solvent-accessible domain, carry greater scientific merit (**Figure S7A**). The docking results reveal that the amide carbonyl group of the covalent linker can establish hydrogen bonding interactions with the amide nitrogen atom of Gly143, whereas the hydroxyl group introduced by the benzyl moiety of BB1 (Building Block 1) can engage in robust hydrogen bonding interactions with the amide carbonyl of His164 (**Figure S7C**). This observation underscores a potentially pivotal factor governing the binding affinity between **LU7** and the 3CL^pro^. Considering further molecular optimization, we observe that the S1 site exhibits considerable unoccupied space. Upon overlaying complexes of some previously reported 3CL^pro^ inhibitors, we find that fragments binding to the S1 site predominantly adopt a γ-butyrolactam structure, establishing multiple hydrogen-bonding interactions with His163 and Glu166 (**Figure S8**). In light of this, we introduced analogous structures into the molecular framework to synthesize compounds **LU8**-**LU12**. Subsequent testing of the IC_50_ values of these compounds revealed a significant enhancement in inhibitory activity upon incorporation of this structure (**Figure 3D**). This underscores the potential of derivatives on the piperazine ring as an alternative promising modification site. In conclusion, the selectivity of compound **LU12** against human proteins was assessed. The findings indicated values exceeding 50 μM across all evaluated human proteins (**Figure 3G**).

### X-ray Crystal Structures of SARS-CoV-2 3CL^pro^ in complex with LU10

Based on the active compound **LU10**, we elucidated its cocrystal structure with the 3CL^pro^ (**Figure 3E**). The cocrystal structure revealed that **LU10** covalently binds to the catalytic site at Cys145. The triazole ring acted as a scaffold in the structure, linking the active moiety, covalent linker, and DNA, respectively. The crystal structure showed that the active fragment occupy the S2 and S4 sites, and the hydroxyl group derived from the R2 moiety formed multiple hydrogen bonds with Gln189, Thr190, and Gln192, constituting a hydrogen bond network. The nitrogen atom connecting the active moiety also participated in hydrogen bonding with Gln189. The side chain derived from the piperazine ring bound at a different position from the reported γ-butyrolactam binding site (**Figure 4B**). Under the tension exerted by the piperazine ring, the derived angle deviated, extending towards the middle portion between the S1 and S4 sites and forming hydrogen bonds with nearby Glu166. Further molecular docking simulations demonstrated that when the side chain was methylated compound **LU8**, its methyl-derived direction aligned well with that of **LU10**. Extending the side chain to compounds **LU9** and **LU12** resulted in a deviation in the derived direction towards the S1’ pocket, indicating potential influences of side chain length and the conformation of carbon atoms bound to the side chain on the binding mode (**Figure 4C**). Relevant studies based on these findings had not been published. In the crystal structure of this complex, non-conserved water molecule formed hydrogen bond networks with Ser144, His163, and the carbonyl groups of piperazine side chains, further stabilizing the conformation of the piperazine ring. The amide carbonyl group at the covalent binding site interacted via hydrogen bonding with Gly143 and Cys146.

**Figure 4.**
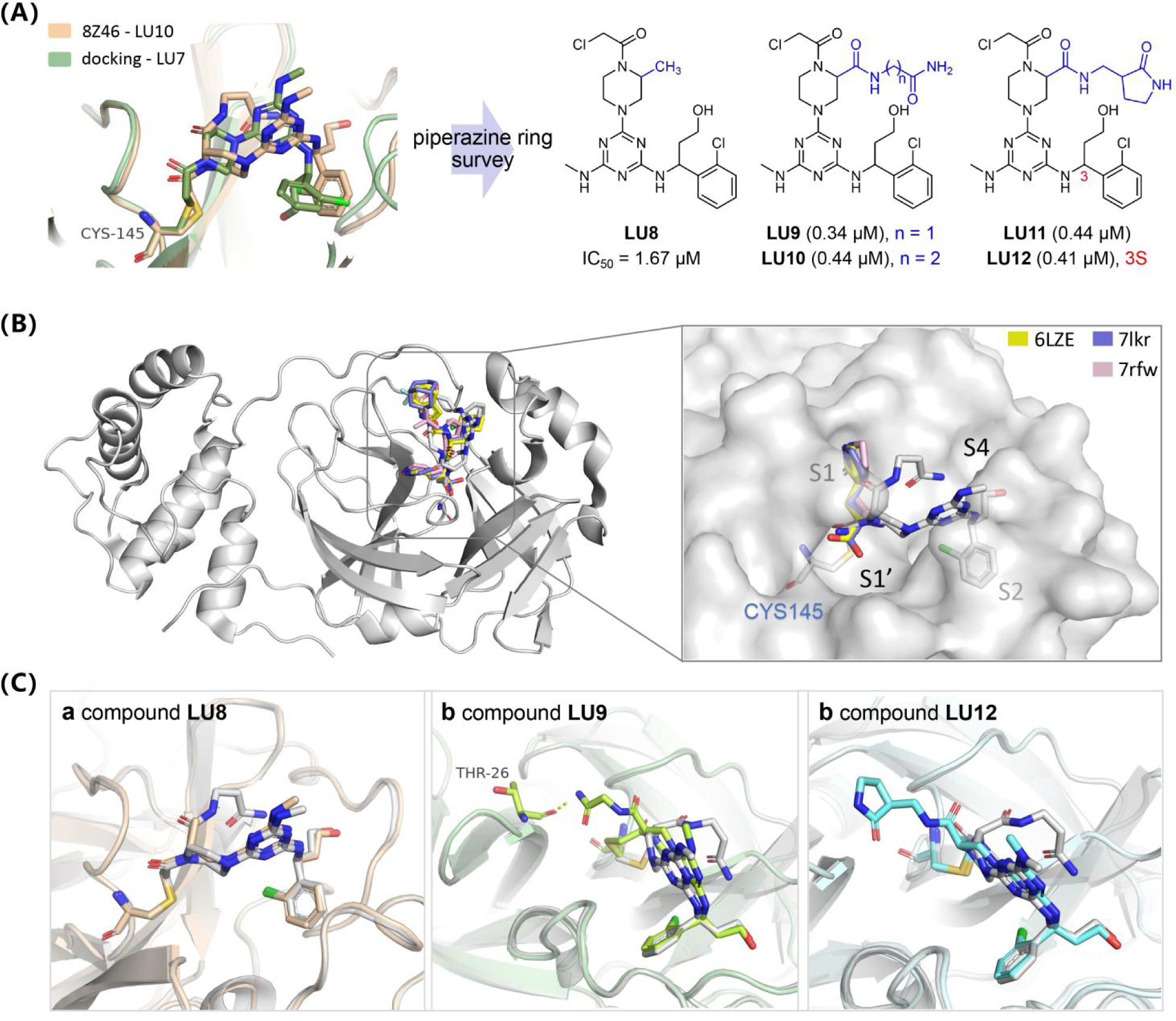
(A) Superposition of crystal structure of **LU10** (parchment) and docking structure of **LU7** (moss), and the structure of **LU8**-**LU12**. (B) Superposition of crystal structure of **LU10** (gray) and reported γ-butyrolactam structures. For clearer visualization, displaying only the structural fragment of γ-butyrolactam. (C) Superposition of crystal structure of **LU10** (gray) and docking structure of **LU8** (coconut), **LU9** (lime), **LU12** (blue)

### Hits Resynthesis and Evaluation of PL^pro^ Inhibition

The ultimate library architectures of DEL0C1 and DEL0C2 exhibit similarities (**Figure 5A**), albeit with minor discrepancies in the reagents utilized during each library synthesis round (**Table S1, Table S2**). Considering the non-specific enrichment observed for cycle 2, namely the linker portion, in both libraries, a comparative analysis of the enrichment levels of structure **XD7**, obtained from screening in both libraries (**Figure 2A**), was conducted. The Select E values representing **XD7** in different libraries were normalized (**Figure 5C**), facilitating the integration of the screening outcomes from the two libraries. We established a standardized E value cutoff of 200, resulting in the identification of five compounds **XD5**, **XD6**, **XD7**, **XD9**, and **XD11** that met the structural criteria. Subsequently, we evaluated the IC_50_ values of these compounds. Our findings revealed a linear correlation between the standardized E values and the IC_50_ activity (**Figure 5B**). Consequently, we were able to quantify the outcomes of the affinity screening, thereby providing a more comprehensive understanding of the DEL screening results. Unlike the screening outcomes observed for 3CL^pro^, PL^pro^ screening against DEL0C1 and DEL0C2 consistently demonstrated selectivity towards the 5-aminoisoquinoline fragment within BB1. Given the limited structural diversity within the library (**Figure S10**), we pursued diverse substitutions of quinoline and isoquinoline structures in cycle 1. However, these compounds exhibited modest PL^pro^ inhibition activity (**Table S8**). Furthermore, de-aromatization of the isoquinoline structure to yield compound **XD13** resulted in a noticeable activity decrease compared to **XD7**. These findings collectively underscore the indispensable activity of the 5-aminoisoquinoline fragment within the compounds, significantly contributing to their overall activity. Nonetheless, exploring 5-aminoisoquinoline with different substituents remains an intriguing avenue for future research. In the context of cycle 2, our previous analysis indicates that the structural segments in this region primarily serve as linkers in protein interactions. Subsequently, we synthesized structures exhibiting high enrichment within the library (**Figure 5C, Table S7**). Through the comparison of normalized E values and activity analyses, it becomes apparent that compounds such as **XD5**, **XD7**, featuring nucleophilic substitution occurring at one end of the triazine moiety connected to a methylene group, manifest favorable activity profiles. This observation suggests that increased flexibility of the N atom proximal to the triazine moiety and its associated C atom may enhance the overall molecular binding to the protein. Furthermore, the removal of the electrophilic warhead portion from the structure results in diminished compound activity (**Table S8**)

**Figure 5.**
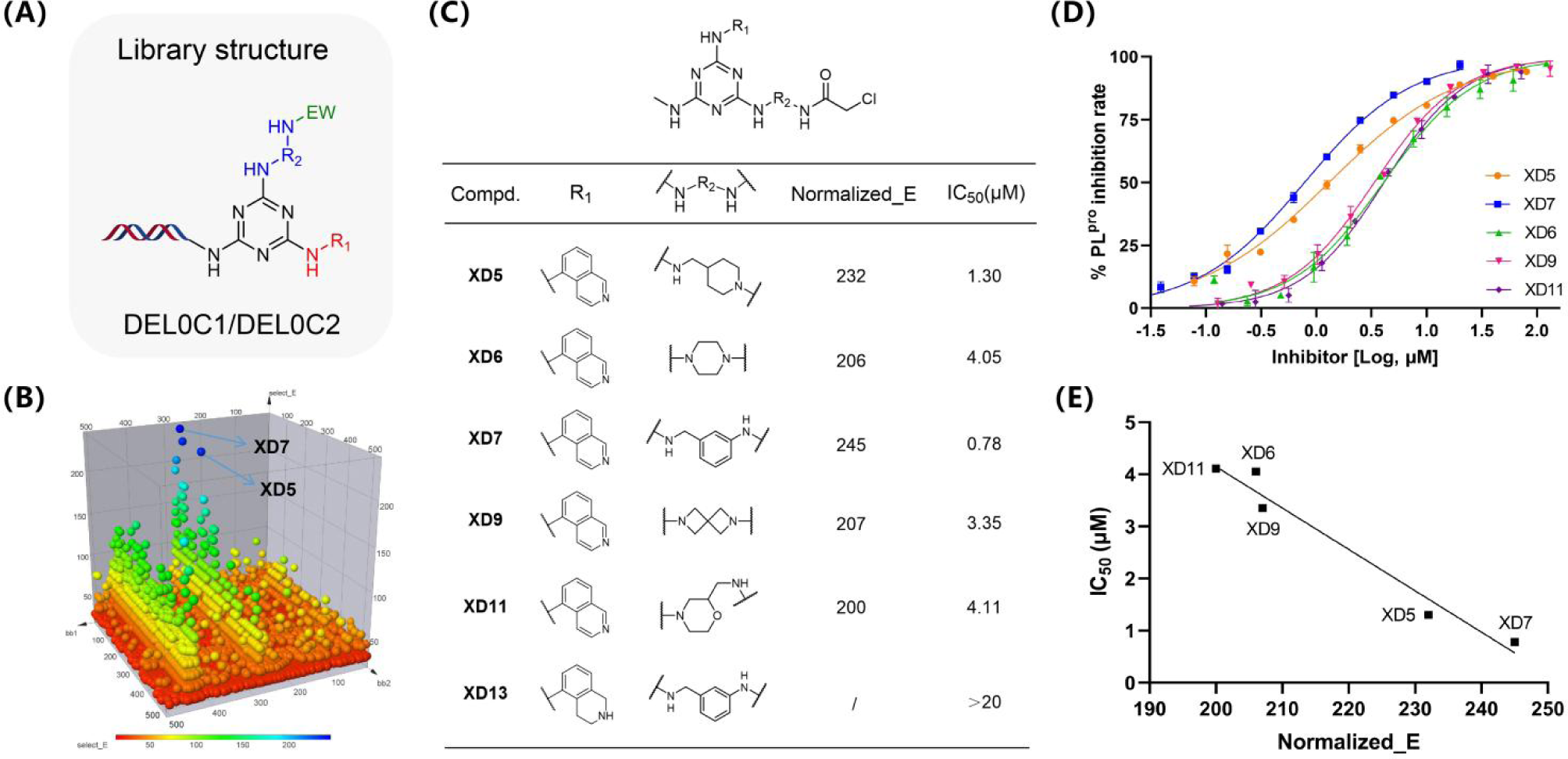
(A) General Structure of DEL0C1 and DEL0C2. (B) Three-dimensional screening structure of PL^pro^, where the vertical axis represents enrichment. (C) Compound structures along with standard E values and IC_50_ values. (D) Dose-response curves for the inhibition of SARS-CoV2 PL^pro^. (D) Graph depicting the relationship between normalized E values and IC_50_ values.

### X-ray Crystal Structures of SARS-CoV-2 PL^pro^ in complex with XD5

To gain a deeper understanding of the precise ligand binding mode, a crystal structure of SARS-CoV-2 PL^pro^ in complex with **XD5** was determined at a resolution of 2.3 Å (PDB:8Z4W). As anticipated, the contiguous electron density is clearly visible between C111 and the acetyl group, confirming the formation of a covalent bond between the protease and **XD5** (**Figure 6F**). The asymmetric unit was comprised of four SARS-CoV-2 PL^pro^ monomers (**Figure 6A**). Across diverse monomers, the compound distinctly formed covalent bonds with a superficial binding site proximal to the catalytic pocket of the SARS-CoV-2 PL^pro^. In all chains, the acetyl group of **XD5** establishes two hydrogen bonds (H-bonds) with Cys111 and Trp106. Specifically, in both chain B and chain D, the compound’s quinoline moiety was observed to form an additional hydrogen bond with Asn267 which is located on the loop region of the protein (**Figure 6B, 6D**). This interaction likely served as a pivotal determinant for the activity attributed to the quinoline moiety. Furthermore, the apparent proximity of Cys270 on the loop region to the **XD5** structure provided a structural rationale for subsequent compound design aimed at covalent binding to Cys270.

**Figure 6.**
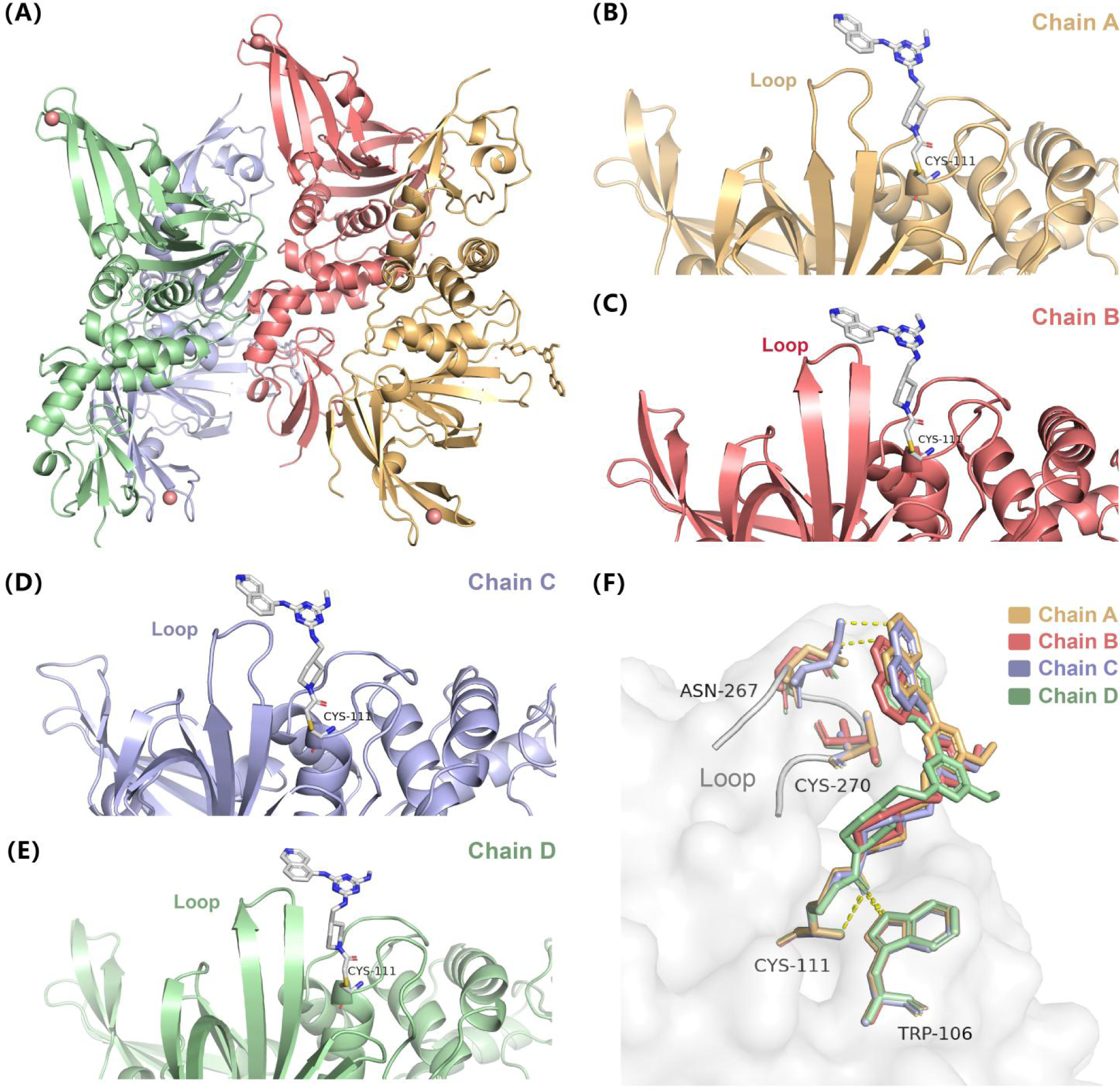
Determination of the co-crystal structure of the SARS-CoV-2 PL^pro^ - **XD5** (PDB:8Z4W). (A) Cartoon structures of the binding pockets of **XD5** and the depiction of four monomers in the structural unit. The crystal structure displays a shallow surface binding site near the catalytic pocket where the compound binds, and similar binding modes are observed in different monomers, such as (B), (C), (D) and (E). (F) The superimposed structures of the compounds binding within the four monomers, along with the nearby amino acid residues.

## Conclusion

In summary, our study introduces a pioneering methodology for discovering covalent inhibitors targeting the key proteases of SARS-CoV-2. Through innovative screening of covalent DNA-encoded libraries, we successfully identified lead compounds exhibiting selective binding to 3CL^pro^ and PL^pro^. By surmounting the limitations of traditional non-covalent screening methods, our research not only broadens the repertoire of potential antiviral agents but also elucidates the crucial role of covalent interactions in drug discovery. This approach holds considerable promise in combating COVID-19 and future viral outbreaks, offering a novel avenue for developing potent therapeutics. Moreover, our findings yield valuable insights into the structural basis of viral protease inhibition, laying the groundwork for rational drug design and optimization. By elucidating the precise binding sites and mechanisms of action of our lead compounds, we provide a foundation for the development of more efficacious antiviral inhibitors.

Furthermore, the methodology employed in our study has been successfully extended to the discovery of other nsp covalent inhibitors, encompassing the complex of nsp 7/8/12 and nsp 10/14, with related research pending publication. The successful application of this study offers pivotal guidance and insights for future antiviral drug development.

In conclusion, our study underscores the significance of innovative screening methodologies in drug discovery targeting the non-structural proteins of SARS-CoV-2 and emphasizes the potential of covalent inhibitors as a novel frontier in antiviral research.

## ASSOCIATED CONTENT

### Supporting Information

The Supporting Information is available free of charge on the ACS Publications website.

Synthesis and characterization of inhibitors, X-ray crystal structure details, Synthetic procedures and MS spectra of DELs on-DNA compounds, On-DNA covalent selection methods, Enrichment profiles, Figure S1-S10, and Table S6-S11. (PDF)

Table S1: Tags code and BBs Smiles data for DEL0C1 (XLSX);

Table S2: Tags code and BBs Smiles data for DEL0C2 (XLSX);

Table S3: Original screening data for DEL0C1 and DEL0C2 against SARS-CoV-2 PL^pro^ (ZIP);

Table S4: Original screening data for DEL0C1 and DEL0C2 against SARS-CoV-2 PL^pro^ C111S (ZIP);

Table S5: Original screening data for DEL0C1 and DEL0C2 against SARS-CoV-2 3CL^pro^ (ZIP); Compounds NMR Spectra, HPLC and HRMS. (ZIP)

### Notes

The authors declare the following competing financial interests (s): X.L., Y.X., and X.W. have submitted two patents application on the compounds. Crystal structures have been deposited in the PDB.

## ACKNOWLEDGMENT

X.L. was supported by the National Natural Science Foundation of China (NSFC) 22377139, 92253305. H.H. gratefully appreciates the National Natural Science Foundation of China (NSFC) 32301050 for support of this work. This work was also supported in part by Shanghai Action Plan for Science, Technology and Innovation (23HC1401200), Shanghai Institute of Pharmaceutical Sciences independently implements research projects (SIMM0220233001) and National Key Research and Development Program for Young Scientists 2022YFC2804400.

