## Supplementary material for "Covalent DNA-Encoded Library Workflow Drives Discovery of SARS-CoV-2 Non-structural Proteins Inhibitors": SI

#### General CoDEL Construction Information

##### Reagents & Instruments

Unless otherwise noted, chemicals were purchased from Sigma Aldrich, J&K Chemical, or Enamine and were used without further purification. DNA tags were purchased from General Biosystems and dissolved in H2O to make the stock solution at 1.8 mM. Tag solutions were stored at -4 ℃ for the short term. Chemical building blocks were obtained from commercial sources and dissolved in DMA to 200 mM; T4 DNA Ligase was obtained from GENEWIZ (600 U/ul). The 10X ligation buffer stock used in ligation reactions was composed as follows: 500 mM Tris pH 7.5, 100 mM MgCl2, and 10 mM ATP.

On-DNA reactions were performed in Eppendorf tubes (small scale) or centrifuge tubes (large scale). All the heating reactions were operated in a PCR amplifier (Bio-Rad®). The analysis for on-DNA reactions was performed by Agilent 6230 Time-of-Flight (TOF) mass spectrometer connected to an Agilent 1260 Infinite II HPLC system.

##### Analysis of DNA Compounds

Mass analysis was performed using Agilent 6230 TOF LC/MS with monitoring at 260 nm. Solvent A: 2.25% hexafluoroisopropanol (v/v) and 0.114% Triethylamine (v/v) in deionized water. Solvent B: 2.25% hexafluoroisopropanol (v/v) and 0.114% Triethylamine (v/v) in 90/10 methanol/water. Flow rate: 0.450 mL/min; Time: 6.00 min. Data visualization and integration were performed with Agilent MassHunter BioConfirm Software. The conversion for DNA-encoded chemistry was determined by full-scanning the TIC trace peak areas and calculating the abundance of the desired product.

##### Ethanol Precipitation for DNA

To a DNA reaction mixture was added 10% (v/v) 5 M NaCl solution and 2.5−3 times the volume of absolute ethanol. The colloidal solution was then allowed to sit at −80 °C for 0.5-1 hour. After centrifugation for 30 minutes at 4 ℃ in a microcentrifuge at 10 K rpm. The above supernatant was removed and the pellet (precipitate) was cooled in liquid nitrogen and then placed on a lyophilizer. After lyophilization, the dry pellet was recovered.

##### Quantification for DNA

The on-DNA Intermediate were purified by a 10 K Spinfilter tube after centrifugation for 30 min at 4 ℃ in a microcentrifuge at 10 K rpm. three times. Use the Thermo ScientificTM NanoDropTM One to quantify the concentration of products under the ds-DNA mode.

#### Materials

Headpiece (5’-/5phos/GAGTCA/iSp9/iUniAmM/iSp9/TGACTCCC-3’) was obtained from Biosearch Technologies, Novato, CA.

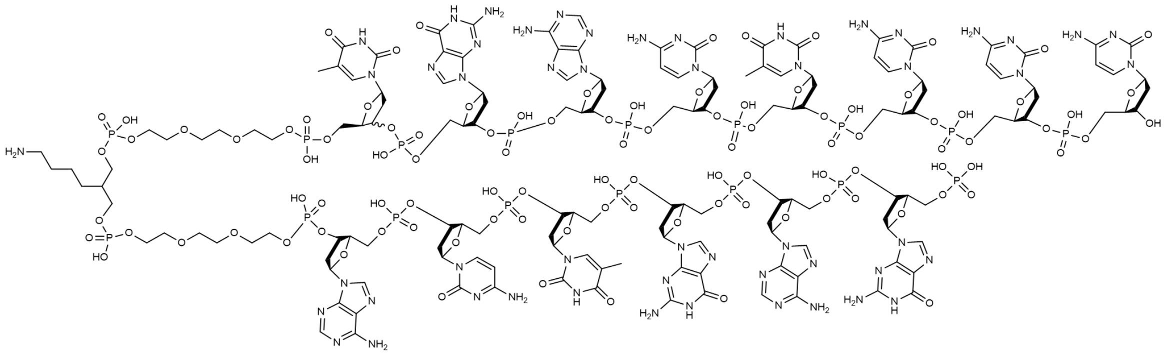

**Chemical Formula: $C_{154}H_{215}N_{52}O_{101}P_{17}$**
**Molecular Weight: 4937.23**

##### *Mass Spectrum of Headpiece:*

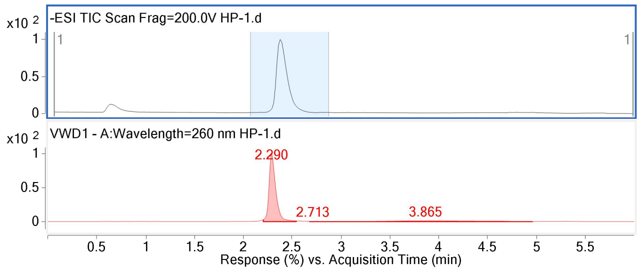


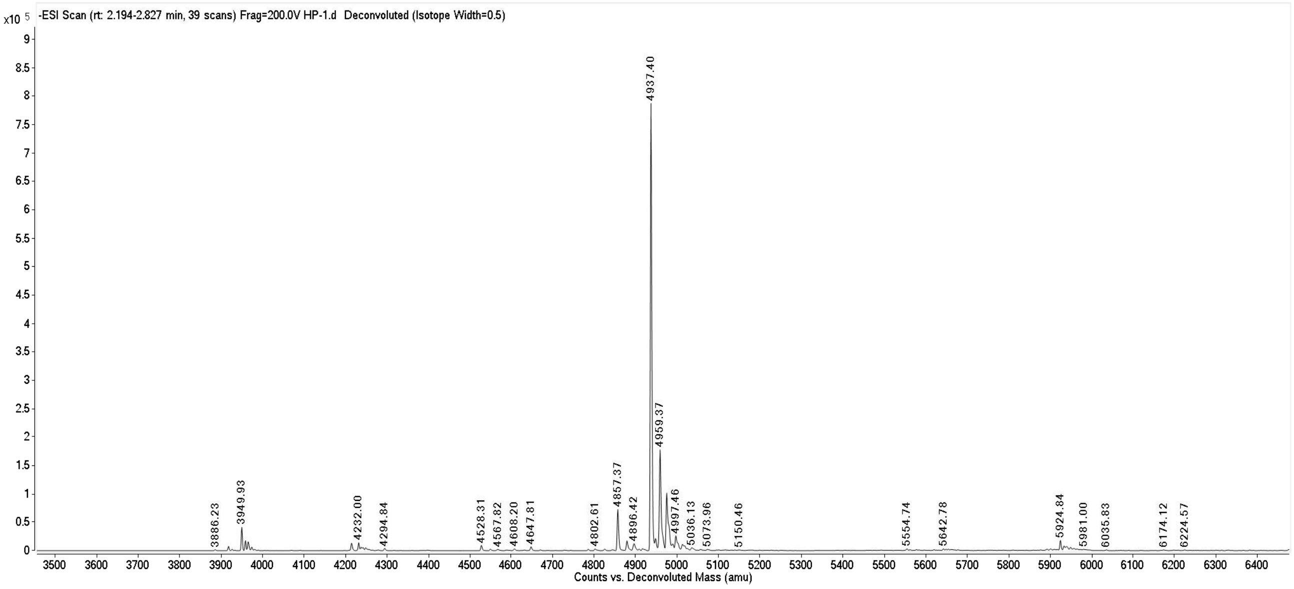


#### Sequence of primer

5’ AAATCGATGTG 3’

3’ GGTTTAGCTAC 5’

**The sequence of the extended headpiece**

TGACTCCCAAATCGATGTG 3’

ACTGAGGGTTTAGCTAC 5’

**Tags:** The DNA tags contained an 11bp coding region, flanked by two 2-base 3’ overhangs, all 5’-ends were phosphorylated.

5’ XXXXXXXXXAG XXXXXXXXXGT XXXXXXXXXGA

3’ ACXXXXXXXXX TCXXXXXXXXX CAXXXXXXXXX

DNA tags were purchased from General Biosystems and dissolved in H2O to make the stock solution at 1.8 mM. Tag solutions were stored at -4 °C for the short term. Chemical building blocks were obtained from commercial sources and dissolved in DMA to 200 mM; T4 DNA Ligase was obtained from GENEWIZ (600 U/ul). The 10X ligation buffer stock used in ligation reactions was composed as follows: 500 mM Tris pH 7.5, 100 mM MgCl2, and 10 mM ATP.

**Preparation of AOP-Headpiece**

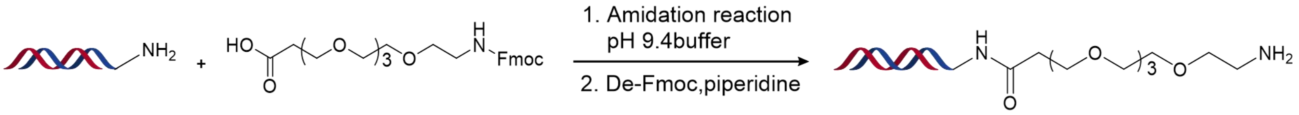


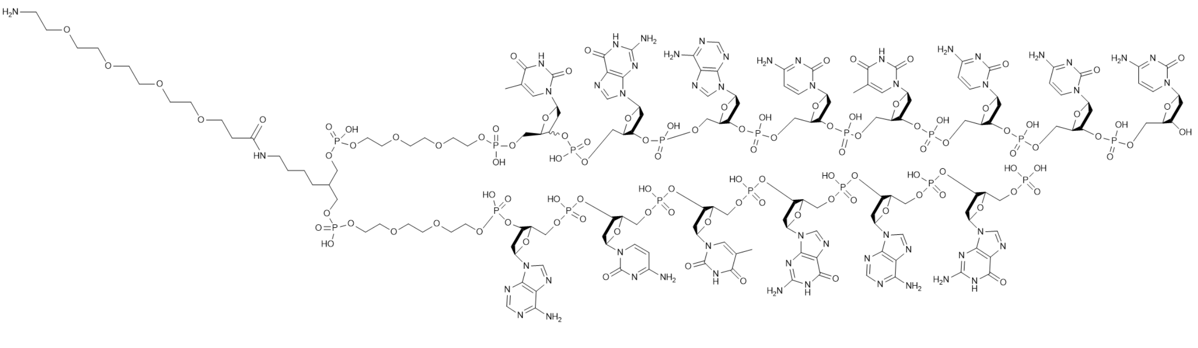

**Chemical Formula: $C_{165}H_{236}N_{53}O_{106}P_{17}$**
**Molecular Weight: 5184.5220**

***Materials:***

Headpiece: 1 mM in water

AOP: Fmoc-15-amino-4,7,10,13-tetraoxapentadecanoic acid 200 mM in DMA

DIPEA: 200 mM in DMA

HATU: 200 mM in DMA

pH 9.4 Buffer: 500 mM sodium borate in water (pH 9.4)

***Procedure:***

1) Mixing AOP with equal volume HATU and DIPEA to prepare **solution 1**.

2) To the headpiece solution (10 μmol, 10 mL), was added 10 mL pH 9.4 sodium borate buffer solution, 10 equiv. of **solution 1** (1.5 mL), and mix.

3) React at room temperature for 2 hours.

4) The product was obtained by ethanol precipitation as described above.

5) The lyophilized pellet was then deprotected by exposure to 1 mL of 10% piperidine in water.

6) The deprotected product was obtained by ethanol precipitation as described above.

***Mass Spectrum of AOP-Headpiece:***

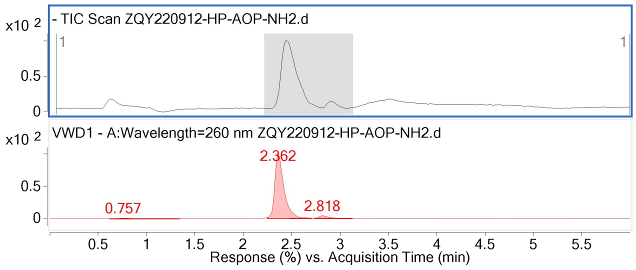


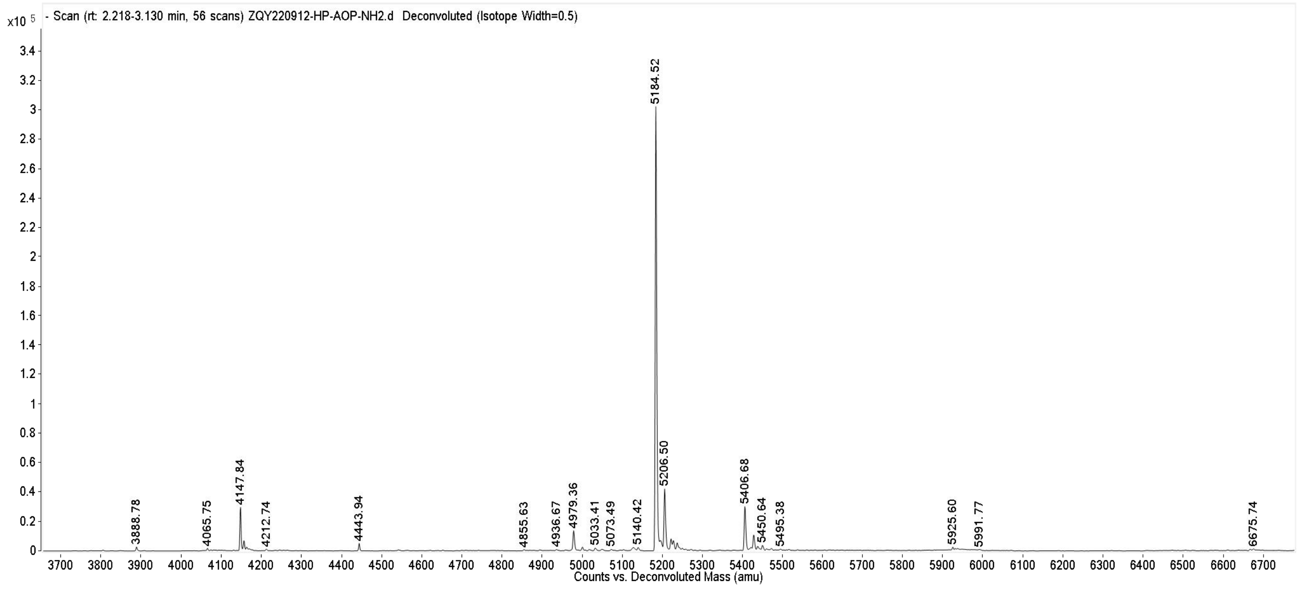

#### General protocol for T4 ligation

***Materials:***

DNA-linked compound: 1 mM in water

tag: 1.8 mM in water

T4 DNA ligase

T4 ligation buffer

***Procedure:***

1) To the DNA-linked compound (5 nmol, 5 uL), was added 9.4 uL dd $H_2O$, 3 uL tag (1.1 equiv, 5.5 nmol), 2 uL T4 buffer, and 0.6 uL T4 ligase. The mixture was vortexed and incubated at 16 °C for 16h.

2) Add 5 M NaCl solution (10 % by volume) and cold ethanol (2.5 times by volume, ethanol stored at -80 °C). The mixture was stored at -80°C for more than 30 minutes.

3) Centrifuge the sample for around 30 minutes at 4 °C in a microcentrifuge at 10 K rpm. The above supernatant was removed and the pellet (precipitate) was cooled in liquid nitrogen and then placed on a lyophilizer. After lyophilization, the dry pellet was recovered.

***Mass Spectrum of AOP-Headpiece-Primer:***

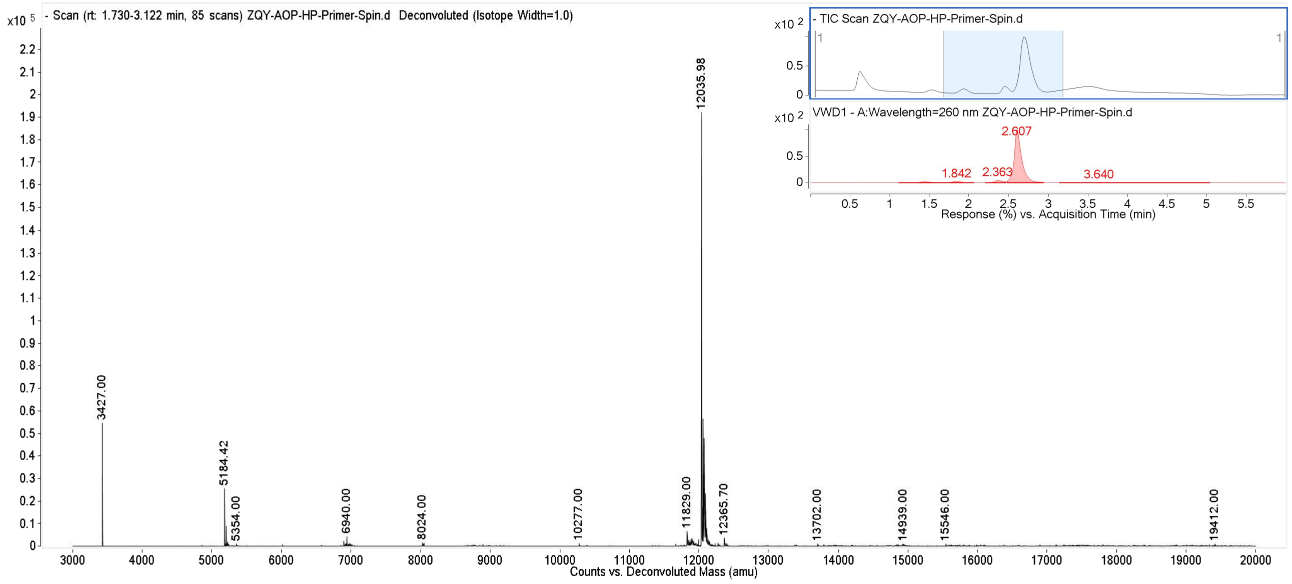

#### Covalent DEL Screening Methods

##### Reagent and Material

Magnetic beads: 20 μL of 88832 (purchased from ThermoFisher).

Protein: Target protein 5 μg

DEL library: 2.5 nmol

SDS-PAGE: 12% (formula in TaKaRa product catalog, page T-11)

WB (self-prepared): 50 mM PBS (pH 7.4), 0.1% Tween-20, 150 mM NaCl

SB (self-prepared): 50 mM PBS (pH 7.4), 0.1% Tween-20, 150 mM NaCl, 0.1 mg/mL sheared salmon sperm DNA

Staining solution: 2.5 mg/mL Coomassie brilliant blue R-250, ethanol: acetic acid: ddH2O = 9:2:9

Destaining solution: ethanol: acetic acid: ddH2O = 1:1:8

##### Immobilized protein detection

1) Remove the magnetic beads from the 4°C refrigerator, shake well (can vortex), and pipette 25 μL of magnetic beads into a clean 1.5 mL EP tube. Place the 1.5 mL EP tube on a magnetic stand for a few seconds (keep the magnetic stand on ice), and carefully remove the supernatant after it becomes clear (try to avoid touching the beads).

2) Add 200 μL of pre-prepared WB (kept on ice), mix well, place on a magnetic stand for a few seconds, remove the supernatant after it becomes clear, and repeat step 2 twice.

3) Add 125 μL of SB, mix well, transfer 25 μL to a clean 1.5 mL EP tube, labeled as Blank Beads.

4) Place the remaining 100 μL of magnetic beads on the magnetic stand for a few seconds, remove the supernatant after it becomes clear.

5) Dissolve 6 μg of protein in 120 μL of SB (protein amount can be adjusted according to actual conditions), transfer 20 μL to a clean 1.5 mL EP tube, labeled as Input.

6) Resuspend 4 in the magnetic beads using 100 μL of protein solution, mix well, place on a vertical mixer, and gently rotate at room temperature for 0.5 h (incubation time and temperature can be adjusted depending on the protein).

7) Place the magnetic beads on the magnetic stand, briefly centrifuge, wait for a few seconds, transfer the supernatant to a clean 1.5 mL EP tube, labeled as Flow-through.

8) Add 200 μL of SB to the magnetic beads, mix well using a pipette, place on the magnetic stand,

wait for a few seconds, transfer the supernatant to a clean 1.5 mL EP tube, labeled as Wash.

9) Resuspend the magnetic beads in 100 μL of SB, transfer 50 μL to a new 1.5 mL EP tube, labeled as Beads, and heat the remaining beads in a 95°C metal bath for 10 min.

10) Briefly centrifuge, place on the magnetic stand, wait for a few seconds, transfer the supernatant to a clean 1.5 mL EP tube, labeled as Heated Elution.

11) Add 50 μL of SB to the magnetic beads, labeled as Heated Elution Beads.

**Covalent screening:**

1) His-tagged protein (5 μg) and DEL library (2.5 nmol) were mixed in a 60 μL SB system and incubated at room temperature for 1 h.

2) Remove the magnetic beads from the 4°C refrigerator, shake well (can vortex), pipette 50 μL of magnetic beads into a clean 1.5 mL EP tube. Place the 1.5 mL EP tube on a magnetic stand for a few seconds (keep the magnetic stand on ice), and carefully remove the supernatant after it becomes clear (try to avoid touching the beads).

3) Add 200 μL of pre-prepared SB (kept on ice), mix well, place on a magnetic stand for a few seconds, remove the supernatant after it becomes clear, and repeat twice.

4) Add 100 μL of SB, mix well, divide equally into two tubes, label as NTC and Sample. Place on a magnetic stand for a few seconds, remove the supernatant after it becomes clear.

5) Resuspend the magnetic beads in the mixture prepared in step 1, vortex for 10 s.

6) Briefly centrifuge, place the magnetic beads on the magnetic stand, wait for a few seconds, and transfer the supernatant to a clean 1.5 mL EP tube.

7) Add 1000 μL of SB to the magnetic beads, mix well using a pipette, place on a magnetic stand, wait for a few seconds, and then remove the supernatant.

8) Add 100 μL of SB to resuspend the magnetic beads, vortex for 10 s, and incubate in a 95°C metal bath at 1000 rpm for 10 min.

9) Place on ice for 10 min.

10) Briefly centrifuge, place the magnetic beads on the magnetic stand, wait for a few seconds, and remove the supernatant.

11) Add 100 μL of SB to resuspend the magnetic beads, vortex for 10 s, and set aside.

### On-DNA Compounds Synthesis of SARS-CoV-2 3CL$^{pro}$

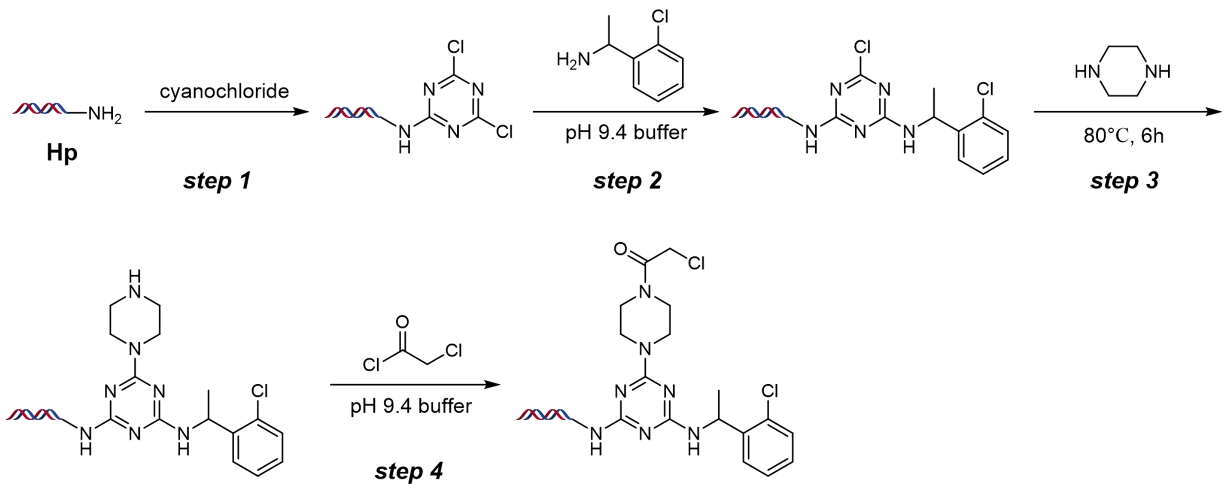


**Scheme S1.** On-DNA compound synthesis of SARS-CoV-2 3CL$^{pro}$

#### *Mass Spectrum of step 1*

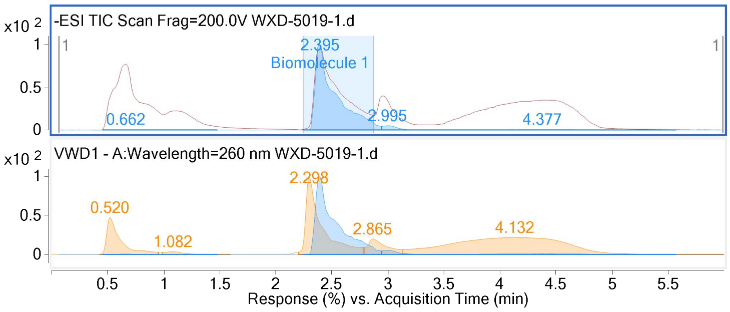


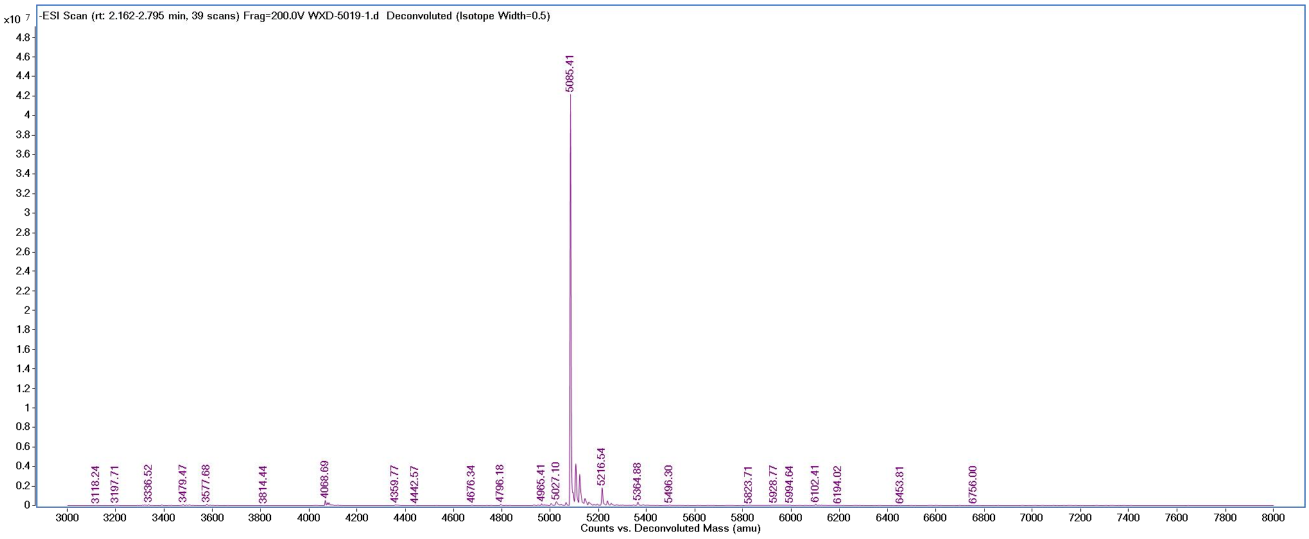


#### *Mass Spectrum of step 2:*

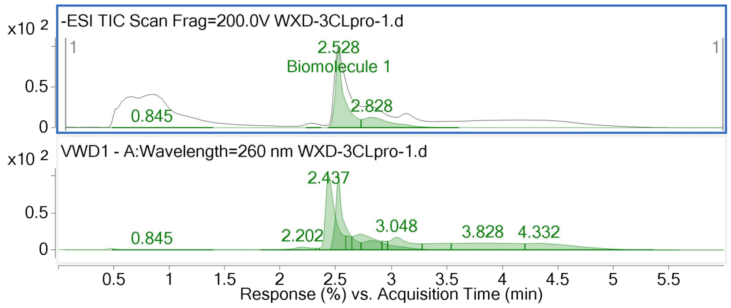


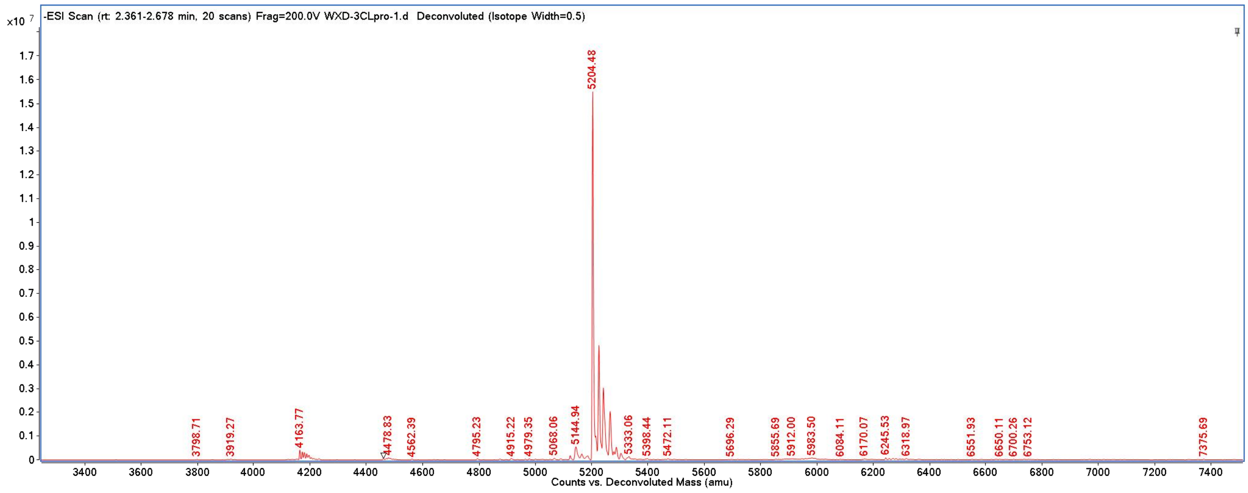


#### *Mass Spectrum of step 3:*

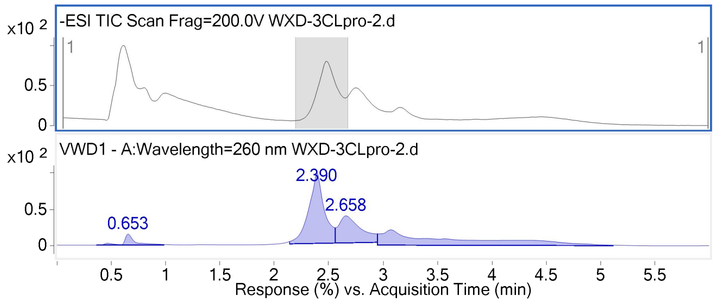


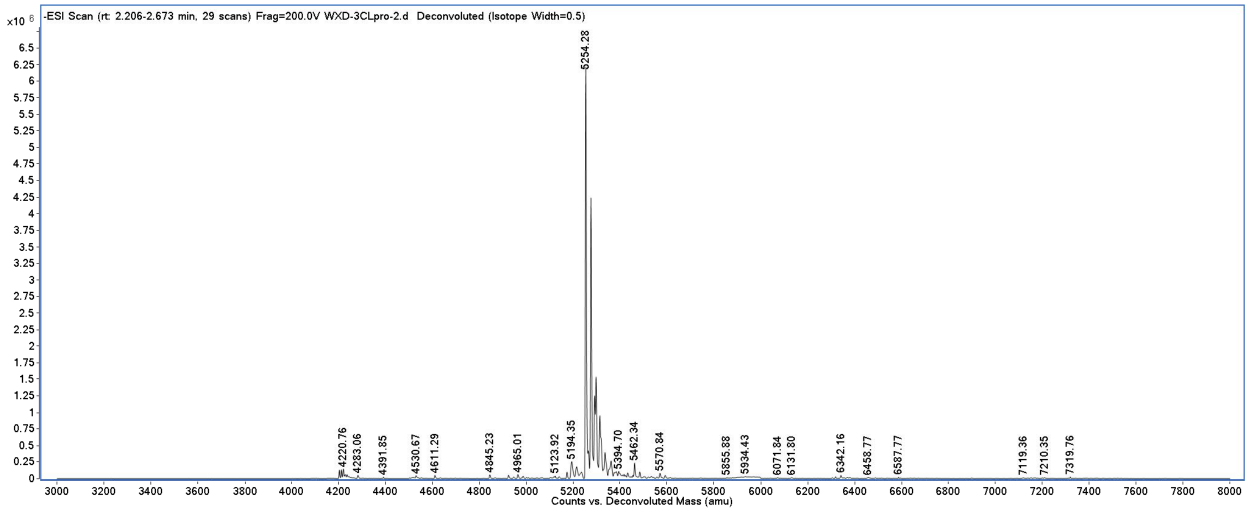

*Mass Spectrum of step 4:*

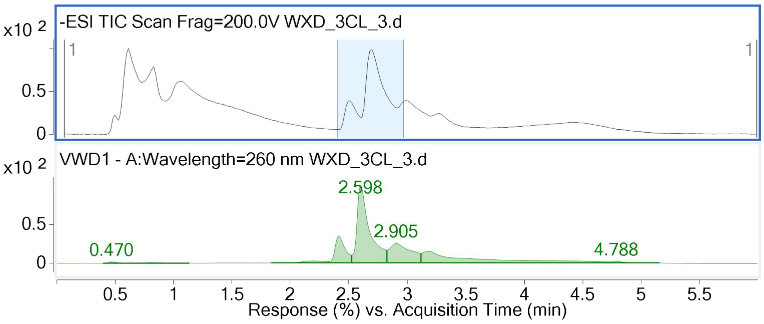


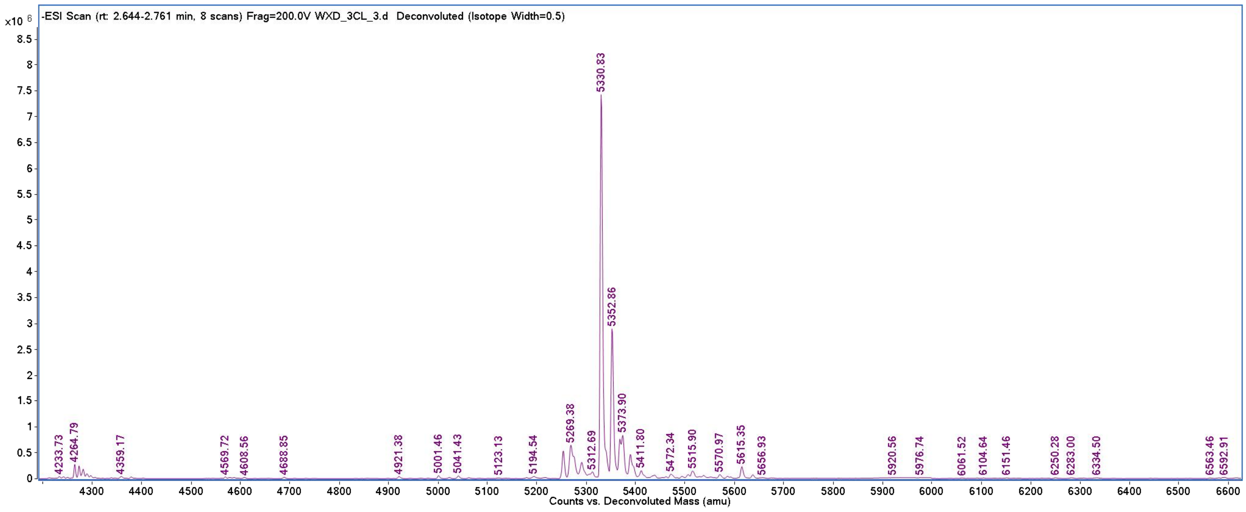


### On-DNA Compounds Synthesis of SARS-CoV-2 PL$^{pro}$

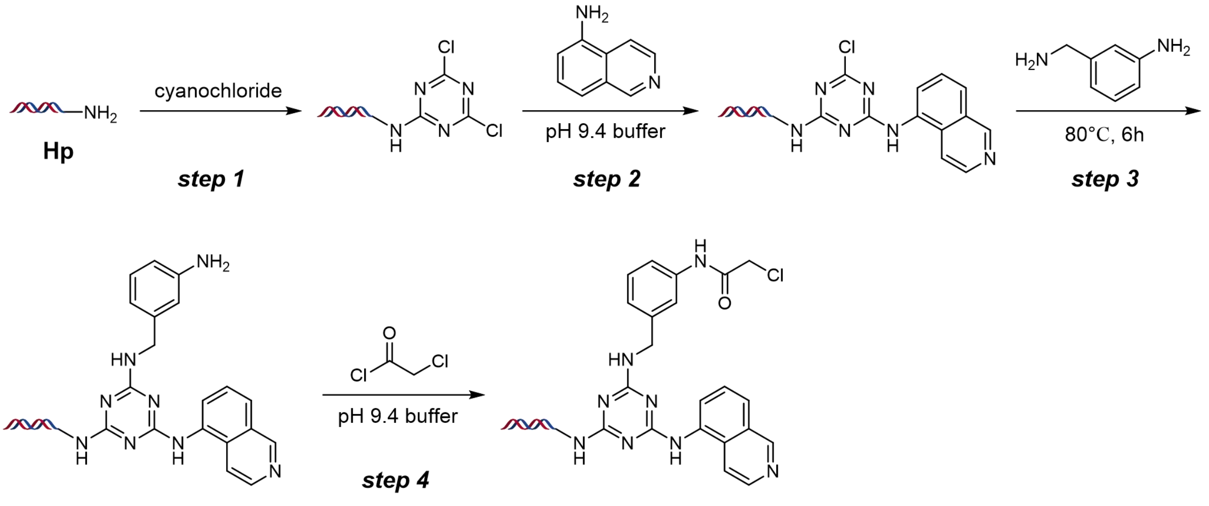


**Scheme S2.** On-DNA compound synthesis of SARS-CoV-2 3CL$^{pro}$

*Mass Spectrum of step 1*

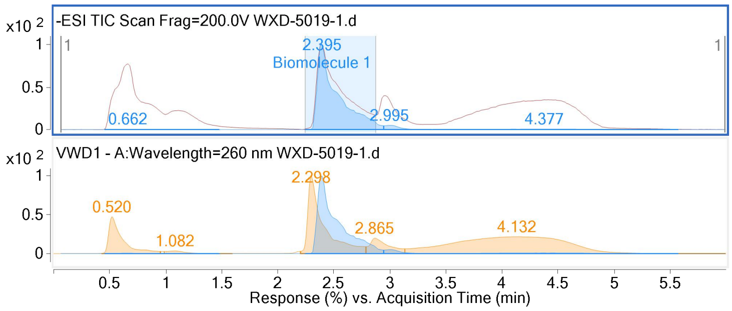


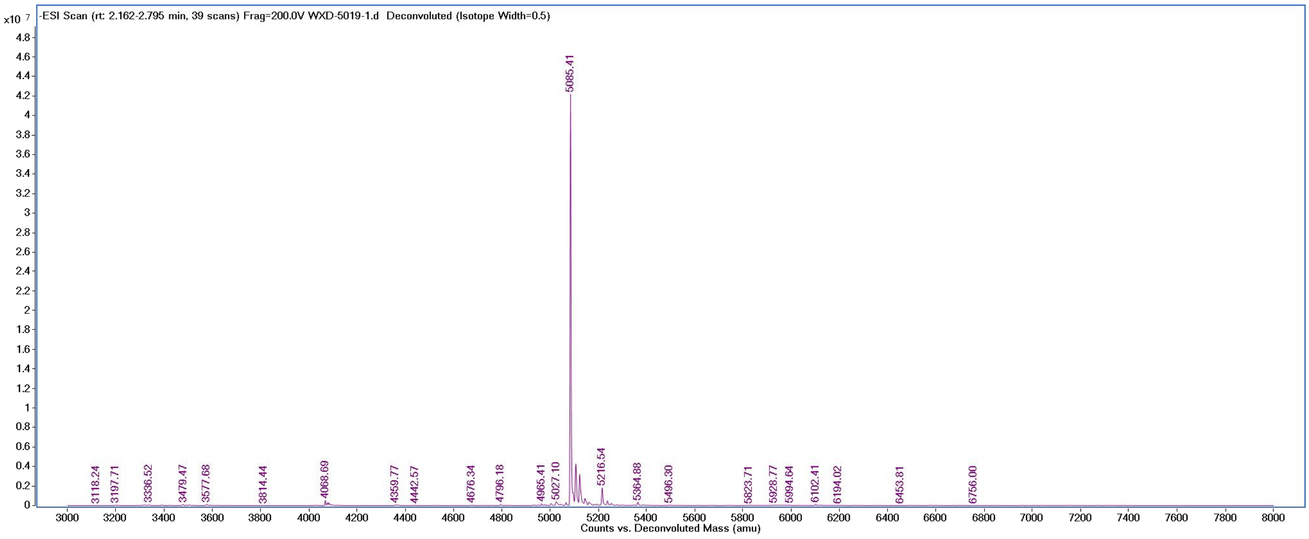


#### *Mass Spectrum of step 2*

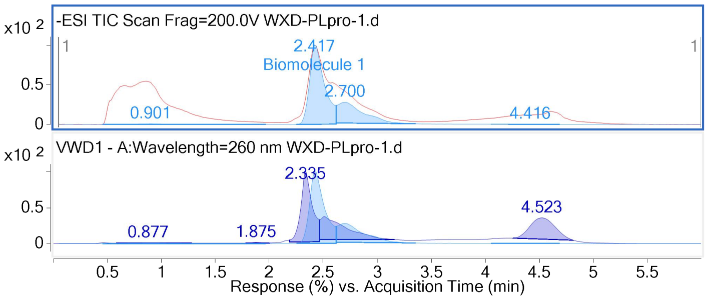


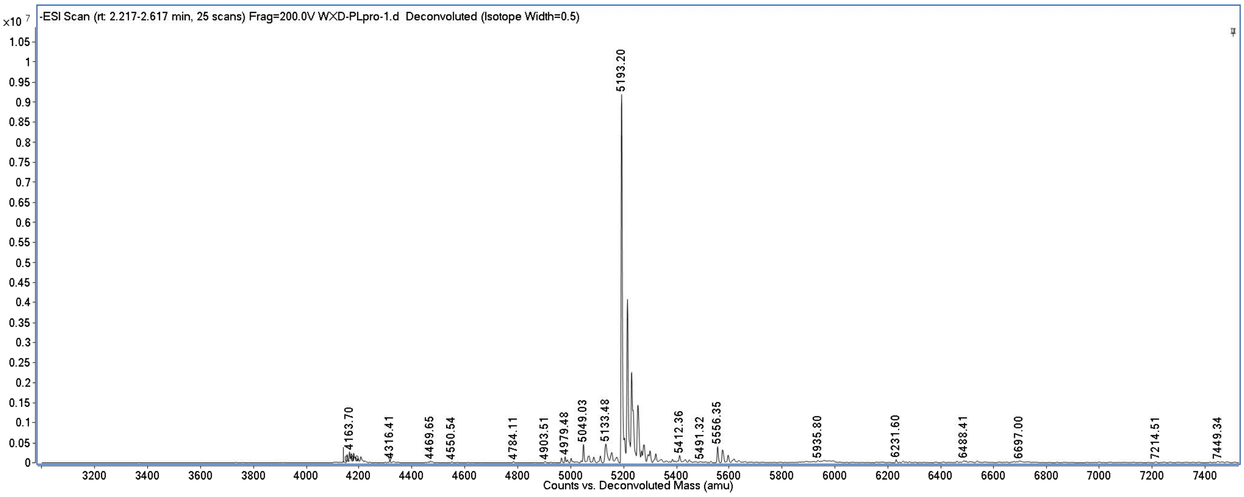

#### *Mass Spectrum of step 3*

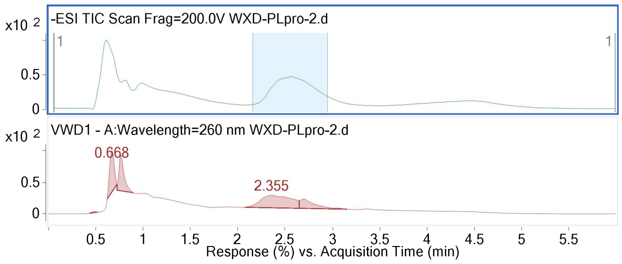


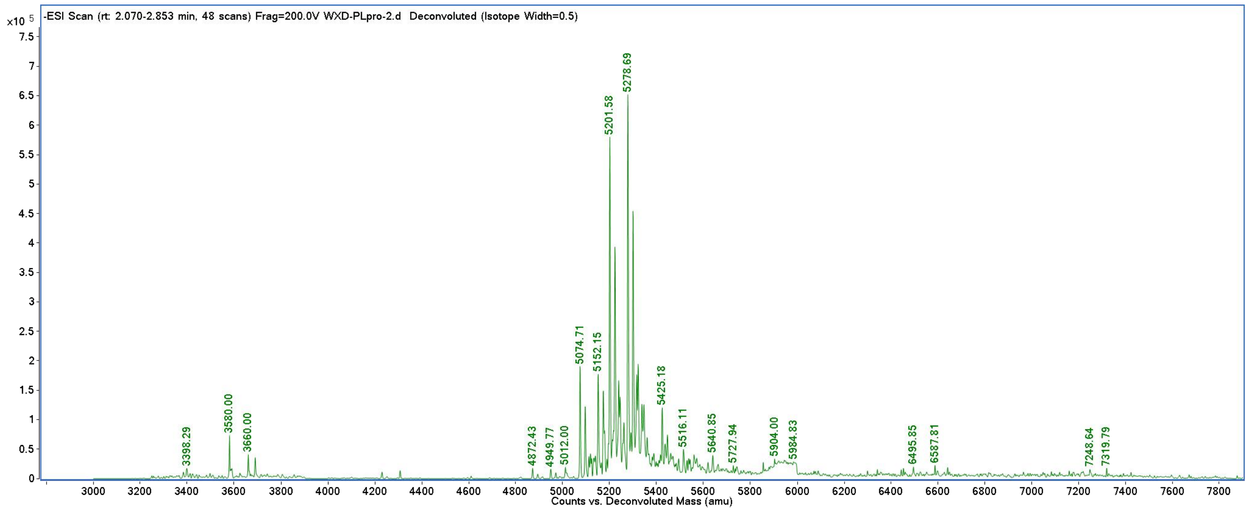


#### *Mass Spectrum of step 4*

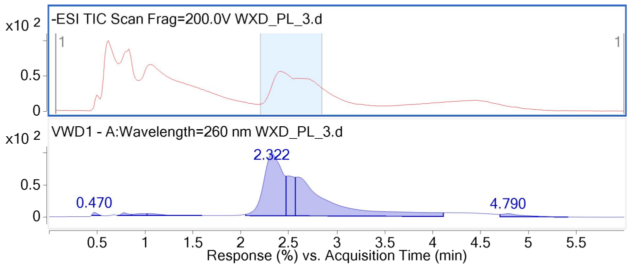


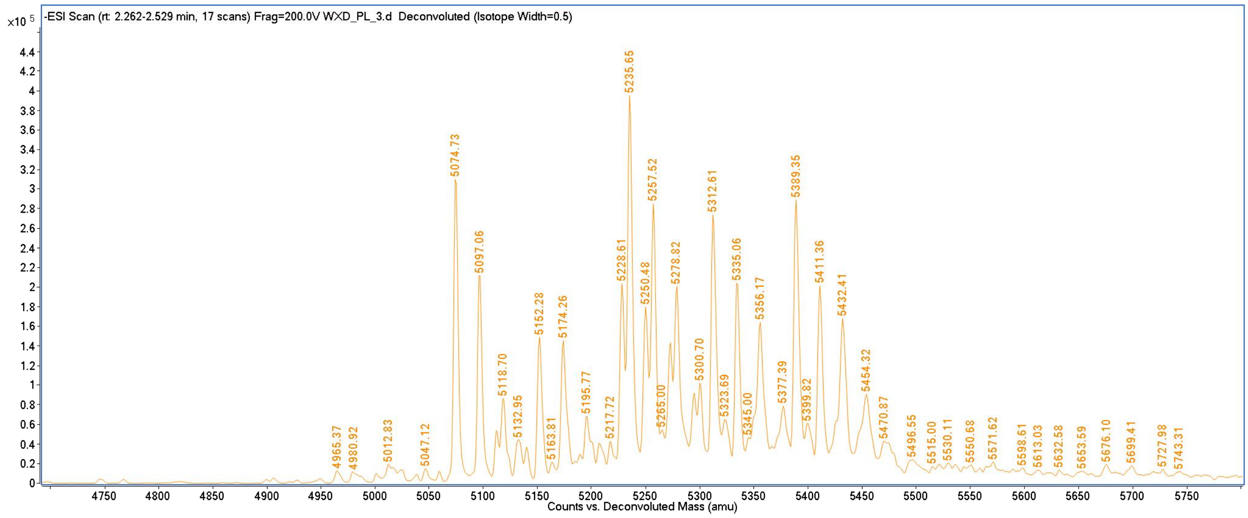

### On-DNA Control Compounds synthesis

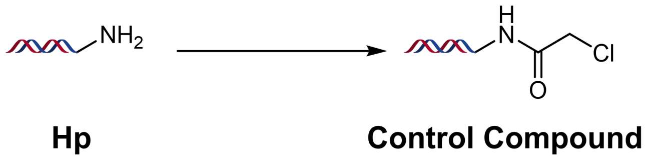


**Scheme S3.** On-DNA Control compound synthesis

#### *Mass Spectrum of On-DNA Control Compound*

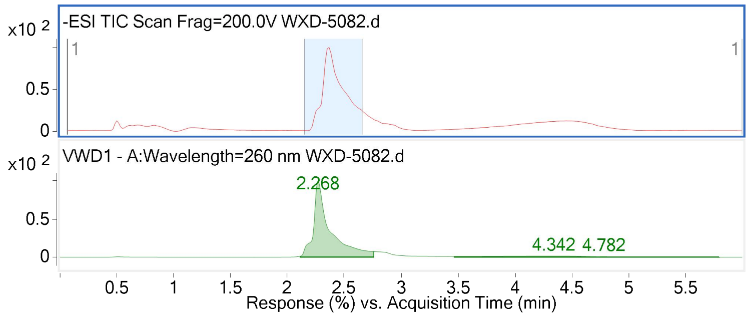


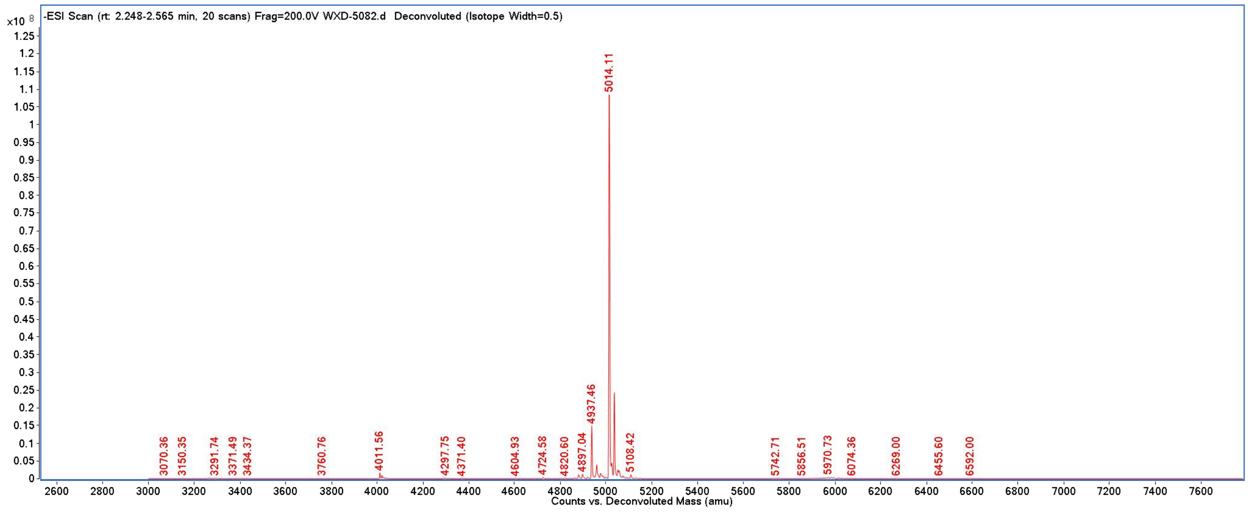

#### Off-DNA Compounds Biological Activity Assay

##### Expression and purification of SARS-CoV-2 PL$^{pro}$

The expression plasmids were transformed into E. coli. Rosetta (DE3) competent cells. The cells were grown in LB medium to an OD600 of 0.8 and induced by IPTG at a final concentration of 0.5 mM and shook at 18 °C overnight. The His-SUMO2-SARS-CoV-2 PL$^{pro}$ was first purified by the Ni-NTA column (GE Healthcare) and cleaved by SUMO Specific Peptidase 2. The resulting protein samples were further purified by SP-Sepharose (GE Healthcare) and Superdex75 (GE Healthcare). The eluted proteins were stored in a solution containing 25 mM HEPES (pH 7.5) and 2 mM DTT for the subsequent experiments.

##### Enzymatic assays of SARS-CoV-2 PL$^{pro}$

The enzymatic inhibition assays were performed using 96-well plates at room temperature and the fluorogeneic substrates used in the assay were RLRGG-AMC which was synthesized by GenScript. The reaction solution contained the following components: 50 mM HEPES pH 7.0, 0.1 mg/mL BSA, 50 nM SARS-CoV-2 PL$^{pro}$, indicated concentrations of compound or equal volume of solvent (DMSO). After 30 min incubation, reactions were initiated with the addition of RLRGG-AMC to reach a final concentration of 20 μM. After that, the fluorescent signal was immediately measured every 1 min for 5 min with a BioTek H1 plate reader (excitation: 360 nm, emission: 460 nm). The initial velocity of reaction was obtained by fitting the linear portion of the curve and the inhibition rate is calculated by comparing the initial velocity of the compound group to the DMSO group. Half maximal inhibitory concentration (IC50) was determined by nonlinear regression analysis of the does-response curves using GraphPad Prism.

##### Expression and purification of SARS-CoV-2 3CL$^{pro}$

The cDNA of SARS-CoV-2 3CL$^{pro}$ (GenBank: MN908947.3) or SARS-CoV 3CL$^{pro}$ (GenBank: AAP13442.1) was cloned into the pGEX6p-1 vector. To obtain the SARS-CoV-2 3CL$^{pro}$ or SARS-CoV 3CL$^{pro}$ with authentic N and C terminals, four amino acids (AVLQ) were inserted between the GST tag and the full-length SARS-CoV-2 3CL$^{pro}$ or SARS-CoV 3CL$^{pro}$, while eight amino acid (GPHHHHHH) were added to the C-terminal of SARS-CoV-2 3CL$^{pro}$ or SARS-CoV

3CL$^{pro}$. The plasmid was then transformed into BL21 (DE3) cells for protein expression. The N terminal GST tag and four amino acids (AVLQ) was self-cleavable. The expressed protein with authentic N terminal was purified by a Ni-NTA column (GE Healthcare) and transformed into the cleavage buffer (150 mM NaCl, 25 mM Tris, pH 7.5) containing human rhinovirus 3C protease for removing the additional residues. The resulting protein sample was further passed through a size-exclusion chromatography (HiLoad$^{TM}$16/600 Superdex$^{TM}$ 200pg, GE Healthcare). The eluted protein samples were stored in a solution (10 mM Tris, pH 7.5) for the enzymatic inhibition assay, native state mass spectrometry studies, protein crystallization, etc.

**Enzymatic assays of SARS-CoV-2 3CL$^{pro}$**

A fluorescence resonance energy transfer (FRET) protease assay was applied to measure the inhibitory activity of compounds against the SARS-CoV-2 3CL$^{pro}$ or SARS-CoV 3CL$^{pro}$. The fluorogenic substrate (MCA-AVLQSGFR-Lys(Dnp)-Lys-NH2) was synthesized by GenScript (Nanjing, China). The FRET-based protease assay was performed as follows. The recombinant SARS-CoV-2 3CL$^{pro}$ (30 nM at a final concentration) or SARS-CoV 3CL$^{pro}$ (100 nM at a final concentration) was mixed with serial dilutions of each compound in 80 µL assay buffer (50 mM Tris, pH 7.3, 1 mM EDTA) and incubated for 10 min. The reaction was initiated by adding 40 µL fluorogenic substrate with a final concentration of 20 µM. After that, the fluorescence signal at 320 nm (excitation)/405 nm (emission) was immediately measured every 30 s for 10 min with a Bio-Tek Synergy4 plate reader. The initial velocity of reactions added with compounds compared to the reaction added with DMSO were calculated and used to generate $IC_{50}$ curves.

**Crystallization and Data Collection of SARS-CoV-2 PL$^{pro}$ complexed with XD5**

SARS-CoV-2 PL$^{pro}$ (1 mg/mL) was incubated with 100 μM **XD5** in 25 mM HEPES (pH 7.5) at 10 °C for 2 h. The mixture was supplemented with 10 mM DTT and further purified through uperdex75 (GE Healthcare). The eluted proteins, preserved in a solution consisting of 25 mM HEPES (pH 7.5) and 2 mM DTT, were concentrated to a final concentration of 12 mg/mL for crystallization. The crystal of SARS-CoV-2 PL$^{pro}$ in complex with compound 2 was grown by mixing equal volumes of the protein/compound and a reservoir (20% (w/v) PEG 8000, 100 mM HEPES/ Sodium hydroxide pH 7.5) on a 96-well sitting plate at 20 °C. Before data collection, crystals were flash-frozen in liquid nitrogen directly in the presence of the reservoir solution. X-ray diffraction data were collected at beamline BL19U1 at the Shanghai Synchrotron Radiation Facility and processed with the program autoPROC. The structure was solved by the program PHASER and refined with the program PHENIX. The refined structure was deposited into the Protein Data Bank with accession code 8Z4W. The complete statistics as well as the quality of the solved structures are given in Table S11.

### Supplementary Figures

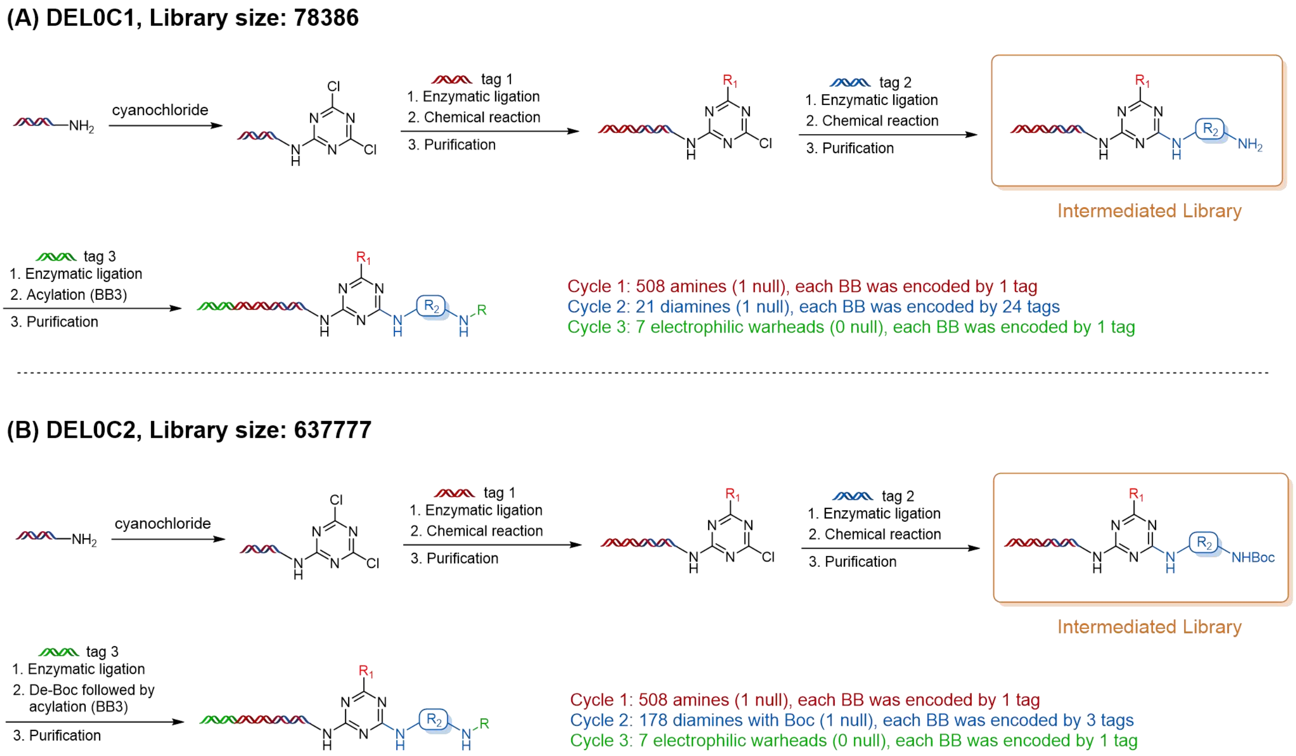


**Figure S1. Synthetic Route of DEL0C1 and DEL0C2.** (A) Cycle 1 was composed of 508 amines and each building block was encoded by 1 tag. Cycle 2 was composed of 21 diamines and each building block was encoded by 24 tags. Cycle 3 was composed of 7 electrophilic warheads and each building block was encoded by 1 tag. (B) Cycle 1 was composed of 508 amines and each building block was encoded by 1 tag. Cycle 2 was composed of 178 diamines with Boc and each building block was encoded by 3 tags. Cycle 3 was composed of 7 electrophilic warheads and each building block was encoded by 1 tag.

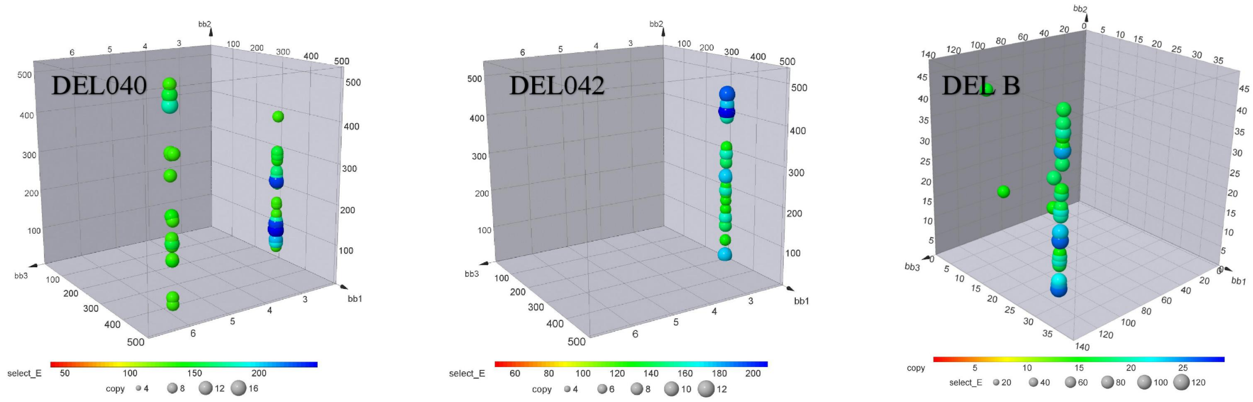


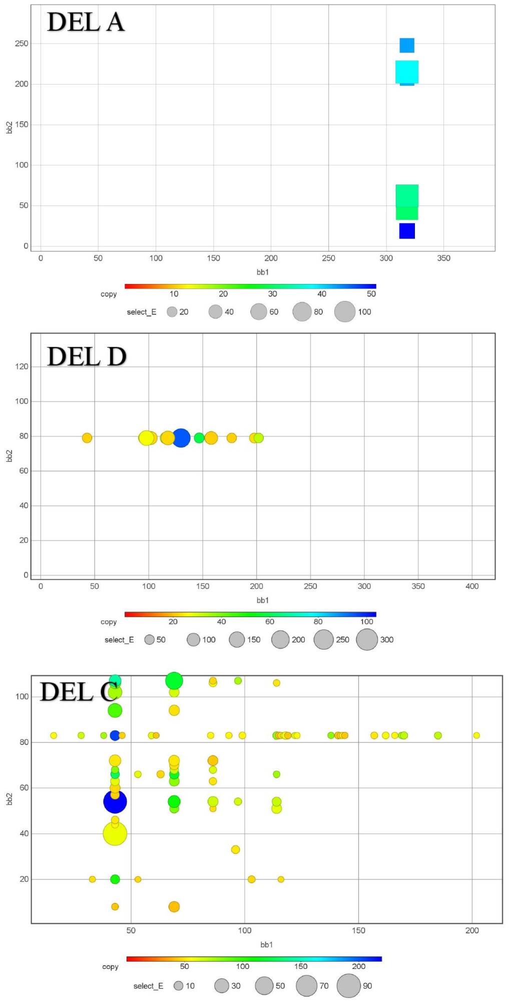


| CoDEL | Target | Structure | Copy | E | notes |
|---|---|---|---|---|---|
| DEL040 | PL$^{pro}$ | 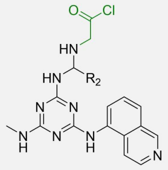 | ~12 | ~200 | This work |
| DEL042 | PL$^{pro}$ | 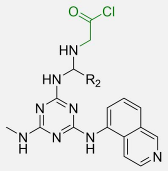 | ~14 | ~200 | This work |
| DEL A | PL$^{pro}$ | 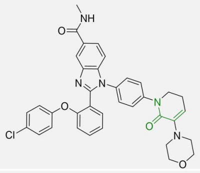 | 38 | 118 | Unstudied |
| DEL B | PL$^{pro}$ | 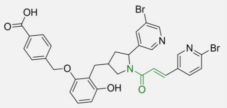 | 26 | 113 | Unstudied |
| DEL C | PL$^{pro}$ | 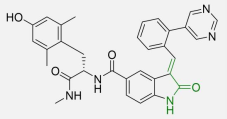 | 220 | 84 | Unstudied |
| DEL D | PL$^{pro}$ | 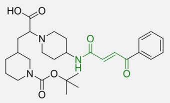 | 95 | 224 | Unstudied |

**Figure S2. Covalent screening result with obvious linear enrichment against SARS-CoV-2 PL$^{pro}$.** DEL0C1 and DEL0C2 are both covalent libraries based on the triazine scaffold, and they exhibit high enrichment (Select_$E_{max}$ > 200). These libraries, DEL A, DEL B, DEL C, and DEL D, have all identified potential hit molecules with activity yet to be validated, exhibiting a more pronounced enrichment relationship compared to non-covalent screening results.

(A)

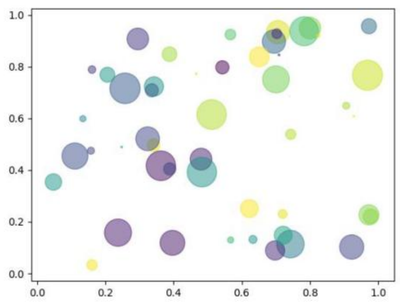


**For 2-cycle DELs:**

$$Select\_E = \frac{\text{Copy / Copy\_total}}{\text{Dilution / Dilution\_total}}$$

***Copy:*** copy number of the molecule, refers to the quantity of a specific compound in the library after screening.

***Copy_total:*** total reads, refers to the quantity of all compound in the library after screening.

***Dilution:*** copy number of the molecule, refers to the quantity of a specific compound in the library before screening.

***Dilution_total:*** total reads, refers to the quantity of all compound in the library before screening.

(B)

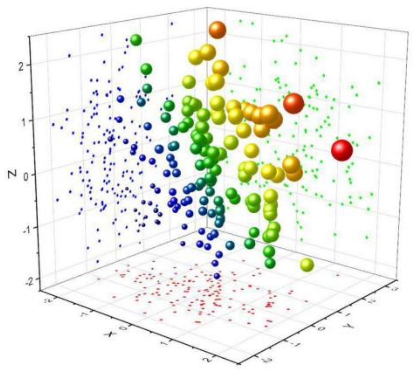


**For 3-cycle DELs:**

For three-round libraries, the same formula can be applied, but with adjustments. Due to the larger initial compound pool in three-round libraries ( Dilution_total >>> Dilution ),

therefore, $\frac{\text{Dilution}}{\text{Dilution\_total}} \approx \frac{1}{\text{Dilution\_total}}$

So, $Select\_E = \frac{Copy / Copy\_total}{1 / Dilution\_total} = \frac{Copy \times Dilution\_total}{Copy\_total}$

***Copy:*** copy number of the molecule, refers to the quantity of a specific compound in the library after screening.

***Copy_total:*** total reads, refers to the quantity of all compound in the library after screening.

***Dilution_total:*** total reads, refers to the quantity of all compound in the library before screening.

**Figure S3. The definition of Select_E and Copy in 2-cycle DELs or 3-cycle DELs.** The screening results of the library construction are quantitatively analyzed using Select_E value and Copy value, with their meanings being correlated between the 2-cycle DELs (A) and 3-cycle DELs (B).

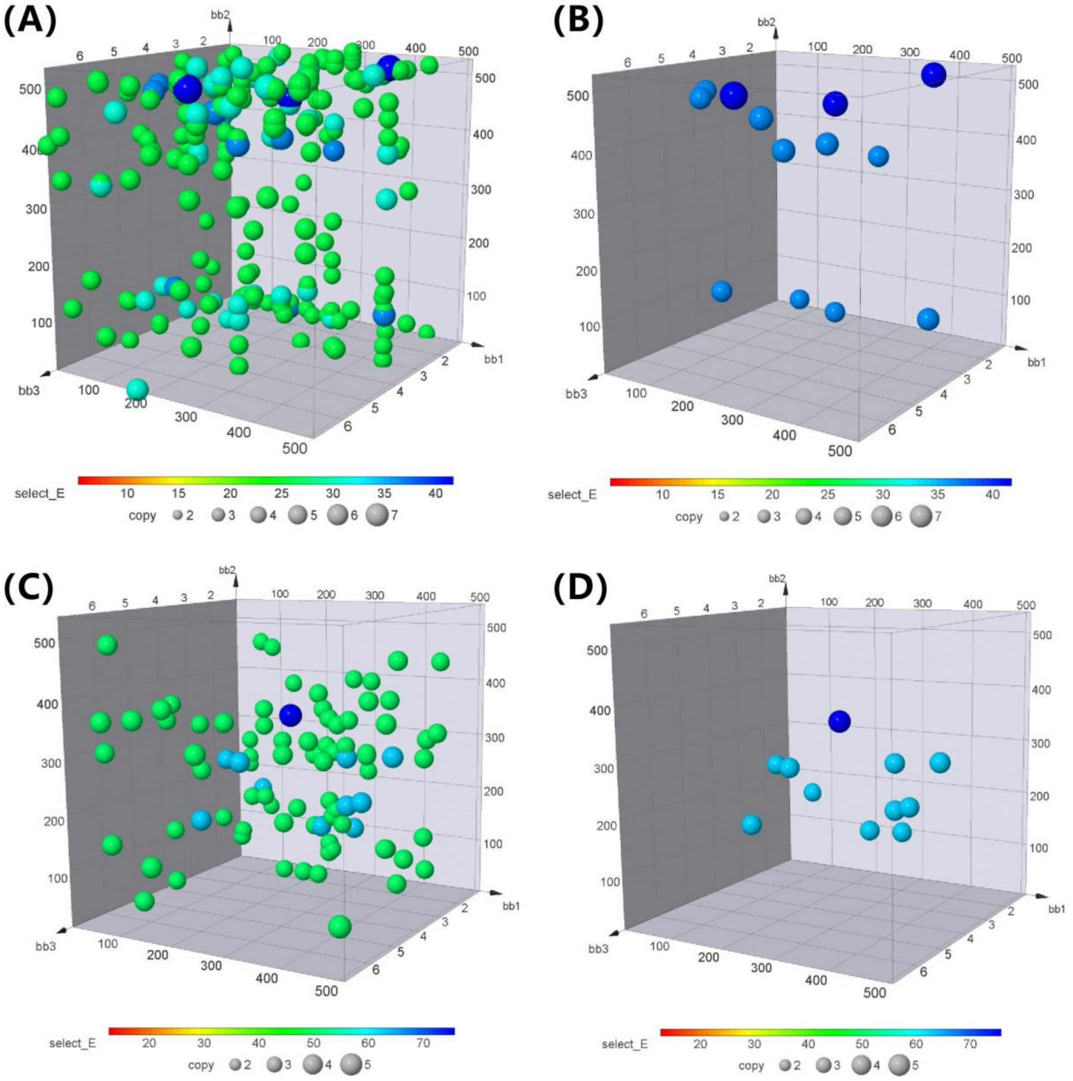


**Figure S4. PL$^{pro}$ C111S Screening data using DEL0C1 and DEL0C2.** Output data analysis profile in DEL0C1 after selection against SARS-CoV-2 PL$^{pro}$ C111S: (A) Copy cutoff at 5, Select_E cutoff at 20. (B) Copy cutoff at 5, Select_E cutoff at 35. Output data analysis profile in DEL0C2 after selection against SARS-CoV-2 PL$^{pro}$ C111S: (C) Copy cutoff at 3, Select_E cutoff at 40. (D) Copy cutoff at 3, Select_E cutoff at 60.

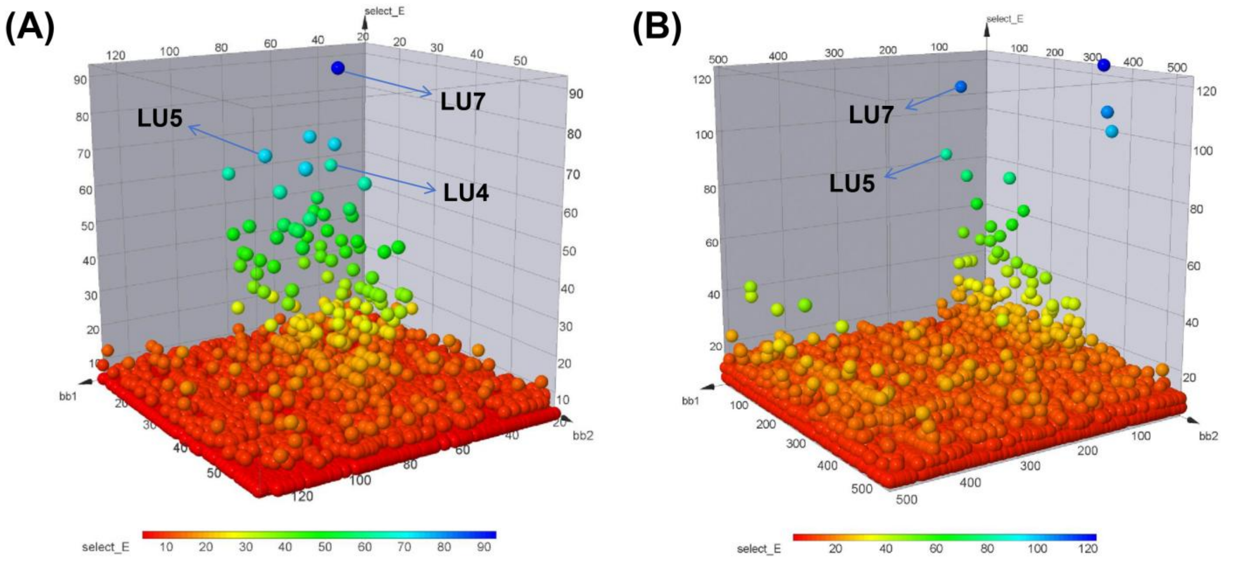


**Figure S5. Three dimensional dot plot showed the most enriched structure.** The two libraries DEL0C1 (A) and DEL0C2 (B) showed the same screening results for cycle 3, therefore the x and y axes of the three-dimensional graph were set as the compound numbers of BB1 and BB2. The Z-axis represents the degree of enrichment at each point.

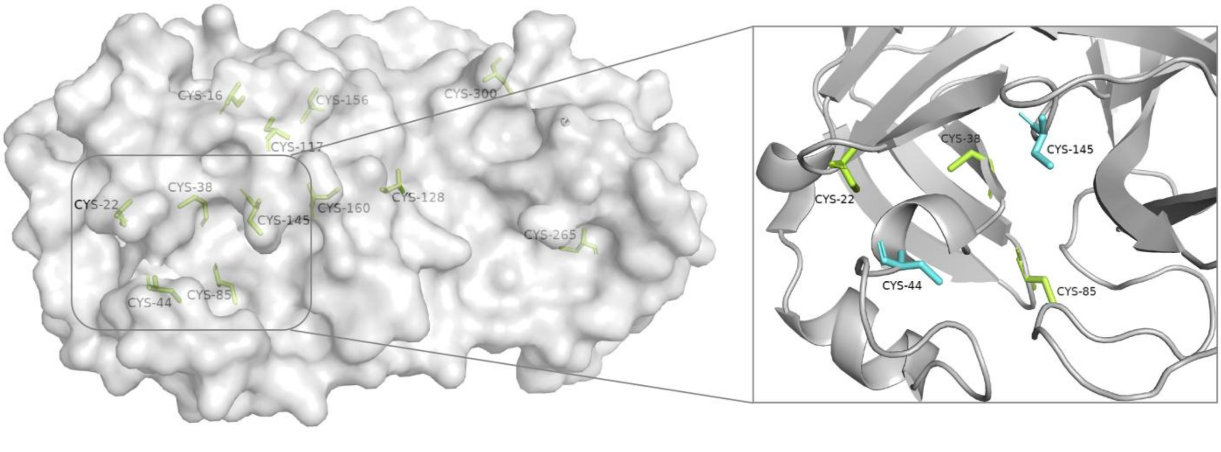


**Figure S6. X-ray crystal structure of SARS-CoV-2 3CL$^{pro}$ (PDB: 8IFT).** The 3CL$^{pro}$contains a total of 12 cysteine residues, show in green, with 5 of them located in the catalytic pocket, Cys22, Cys38 Cys44 Cys85 Cys145. The potentially covalent amino acid residues in the catalytic pocket are Cys145 and Cys44, show in blue.

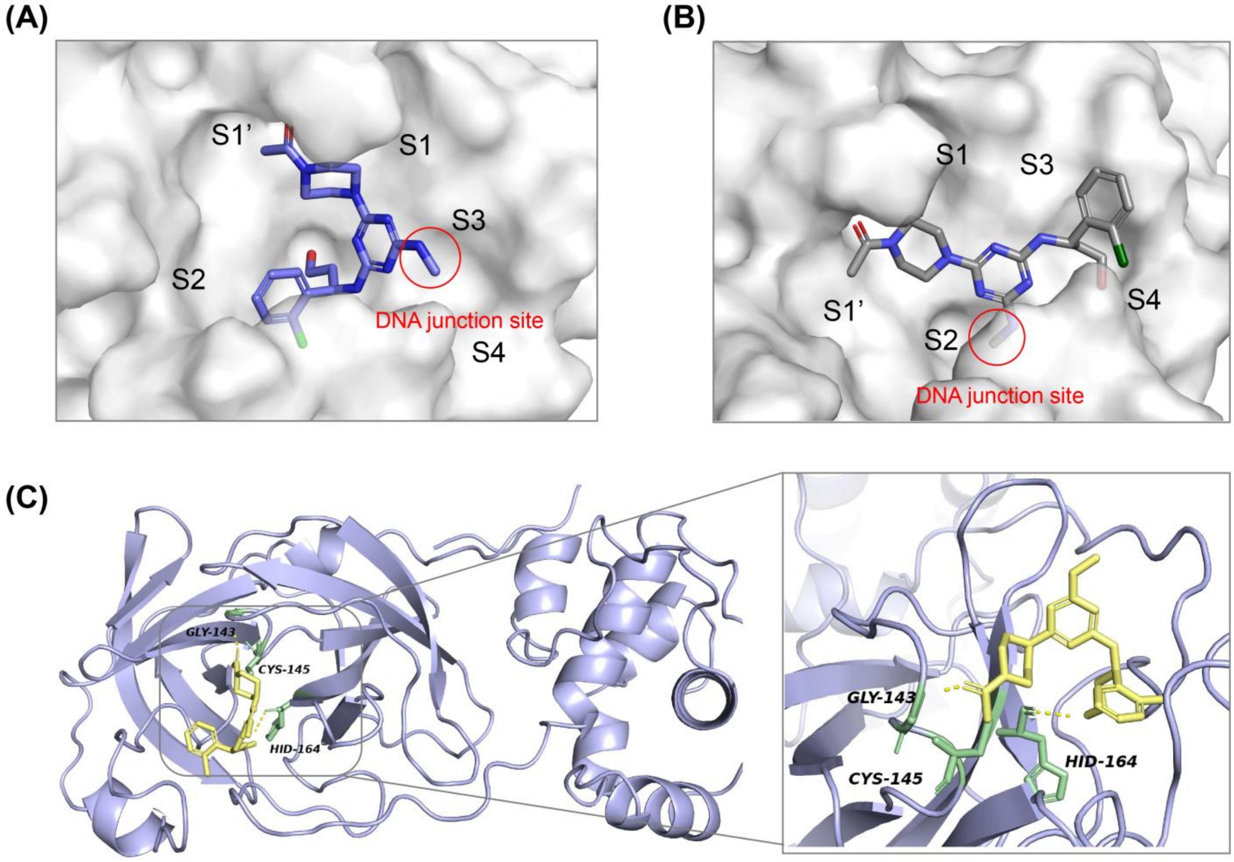


**Figure S7. Docking poses of the compound LU7 in the active site of SARS-CoV-2 3CL^pro^ (PDB: 7RFW).** The docking results reveal two docking conformations. The compounds is depicted in stick representation, distinguished by purple or gray coloration. (A) The docking pose showed that the part connected to DNA extended towards the solvent-accessible region, which was consistent with the DEL screening results. (B) The docking pose showed that the portion connected to DNA extended towards the interior of the protein, which was inconsistent with the DEL screening results. (C) Protein was represented as cartoon, binding site residues were depicted as sticks and labeled, putative hydrogen bonds were highlighted by a yellow dashed line.

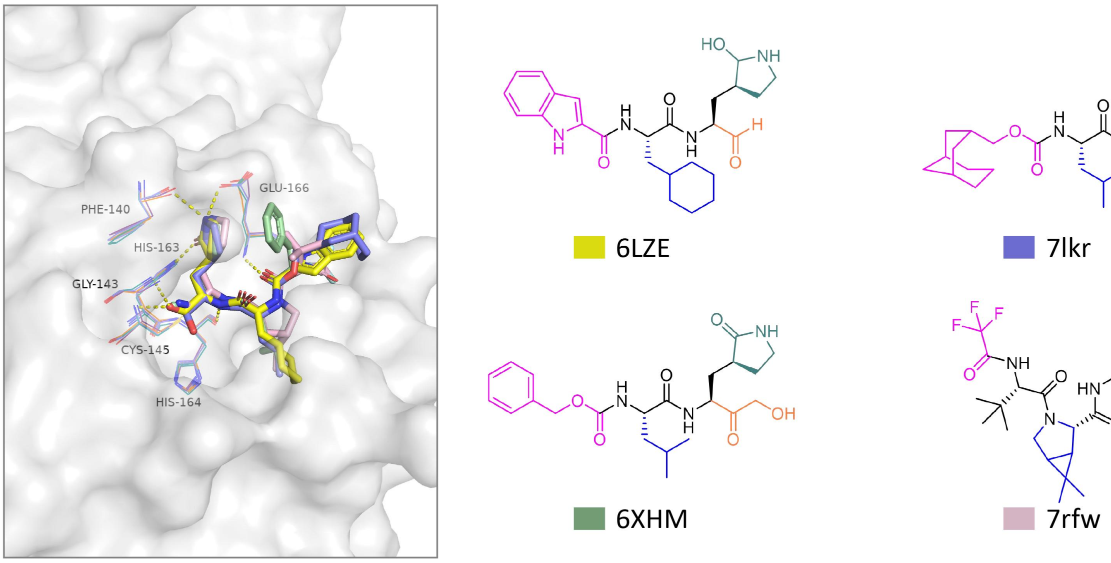


**Figure S8. Superposition of four SARS-CoV-2 3CL$^{pro}$ inhibitor crystal structures.** The chemical structures of inhibitors, their pdb codes are listed at right, with colored boxes corresponding to the coloring used in the structures at left. All of these compounds bind to the S1 site as pyrrolidone structural fragments

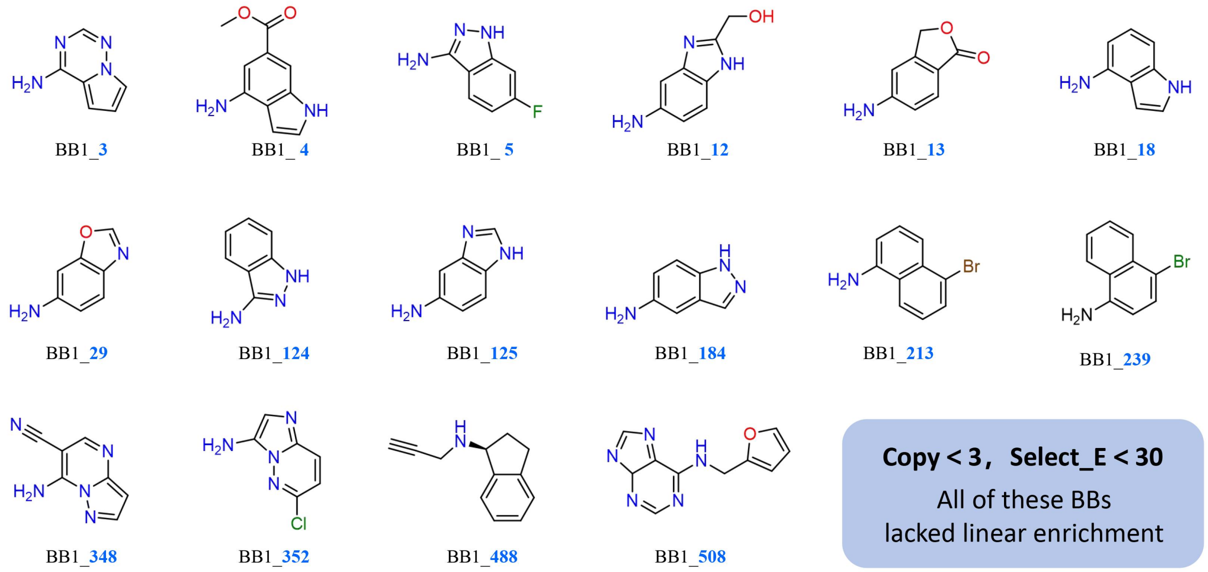


**Figure S9. Aromatic fused-ring structures similar to 5-aminoisoquinoline in DEL0C1/DEL0C2.** None of these structures are enriched in cycle 1, Copy ＜ 5, Select_E ＜ 30. All of these building blocks lacked linear enrichment

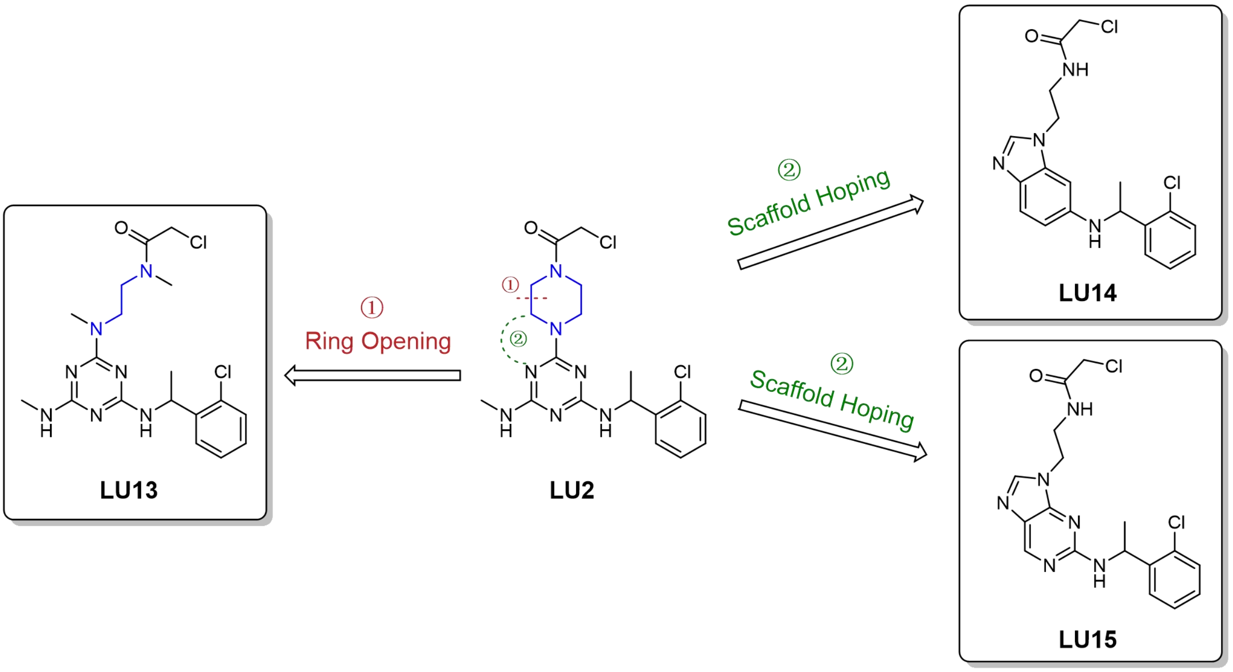


**Figure S10.** Traditional medicinal chemistry modifications of **LU2**, including ring-opening to yield compound **LU13** and skeletal transition to produce compounds **LU14** and **LU15**. All these compounds show no inhibitory activity at 10 μM.

#### Supplementary Tables

Table S1-5: Corresponding datasets provided in the attached supplementary files.

Table S1: Tags codes and BBs smiles data for **DEL0C1**.

Table S2: Tags codes and BBs smiles data for **DEL0C2**.

Table S3: Original screening data for **DEL0C1** and **DEL0C2** against SARS-CoV-2 $PL^{pro}$.

Table S4: Original screening data for **DEL0C1** and **DEL0C2** against SARS-CoV-2 $PL^{pro}$ C111S.

Table S5: Original screening data for **DEL0C1** and **DEL0C2** against SARS-CoV-2 $3CL^{pro}$.

### Table S6

| Protein name | 2 cycle Libraries | Count | 3 cycle Libraries | Count | notes |
|---|---|---|---|---|---|
| **Nsp3/PL$^{pro}$** | DEL Pool | 8 | DEL Pool (non-covalent) | 13 | **Undesried result** |
| **Nsp3/PL$^{pro}$** | DEL Pool | 2 | DEL Pool contain **DEL0C1**, **DEL0C2** | 8 | **This work** |
| **Nsp3/PL$^{pro}$** | DEL Pool | 4 | DEL Pool contain **DEL0C1**, **DEL0C2** | 16 | **This work** |
| **Nsp3/PL$^{pro}$** | DEL Pool | 23 | DEL Pool | 61 | **Unstudied/Undesried result** |
| **Nsp3/PL$^{pro}$** | / | / | DEL A, DEL B | 2 | **Published** |
| **Nsp5/3CL$^{pro}$** | / | / | DEL Pool (non-covalent) | 9 | **Undesried result** |

### Table S7

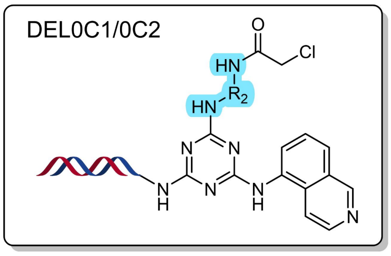


on-DNA hit compounds structure

| DEL040_BB2 | Copy | Select E | DEL040_BB2 | Copy | Select E |
|---|---|---|---|---|---|
| L1 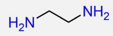 | 8-10 | 103~129 | L6 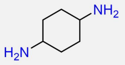 | 8-10 | 103~129 |
| L2 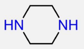 | 8-16 | 103~206 | L7 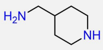 | 8-18 | 103-232 |
| L3 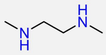 | 10 | 129 | L8 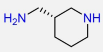 | 8-14 | 103-180 |
| L4 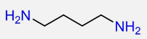 | 8-12 | 103-154 | L9 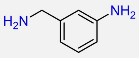 | 9-18 | 116-245 |
| L5 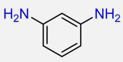 | 9 | 116 | L10 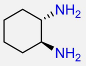 | 8-14 | 103-180 |

| DEL042_BB2 | Copy | Select E | DEL042_BB2 | Copy | Select E | DEL042_BB2 | Copy | Select E |
|---|---|---|---|---|---|---|---|---|
| B1 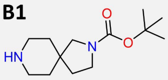 | 9 | 144 | B6 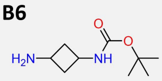 | 8 | 128 | B11 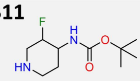 | 10 | 160 |
| B2 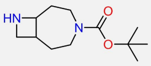 | 8 | 128 | B7 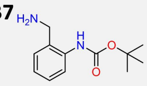 | 12 | 183 | B12 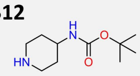 | 10 | 128 |
| B3 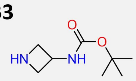 | 11 | 176 | B8 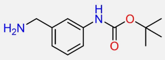 | 13 | 208 | B13 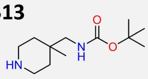 | 8 | 128 |
| B4 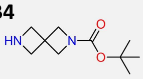 | 10 | 160 | B9 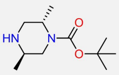 | 8 | 128 | B14 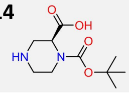 | 8 | 128 |
| B5 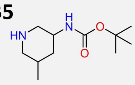 | 9 | 144 | B10 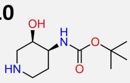 | 8 | 128 | B15 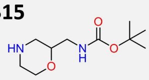 | 10 | 170 |

### Table S8. SARS-CoV-2 PLpro Inhibition Rate of Compounds.

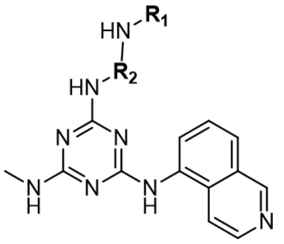

| Compd. | $R_1$ | 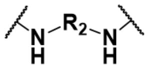 | Normalized_E | Inhibition rate(%)[a] | |
|---|---|---|---|---|---|
| | | | | 4 μM | 20 μM |
| XD1 | 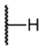 | 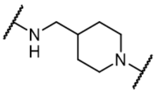 | / | $IC_{50}$ = 35.9 μM | |
| XD2 | 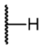 | 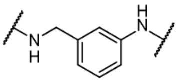 | / | $IC_{50}$ = 13.7 μM | |
| XD3 | 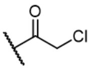 | 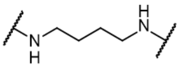 | 154 | 57.2 | 89.2 |
| XD4 | 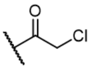 | 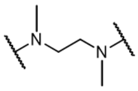 | 129 | 8.6 | 48.3 |
| XD5 | 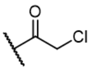 | 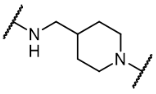 | 232 | 88.6 | 97.1 |
| XD6 | 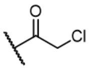 | 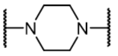 | 206 | 51.2 | 88.3 |
| XD7 | 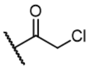 | 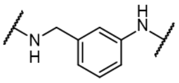 | 245 | 94.9 | 97.9 |
| XD8 | 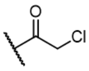 | 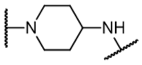 | 150 | 6.2 | 54.7 |
| XD9 | 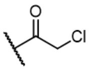 | 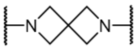 | 207 | 67.8 | 93.6 |
| XD10 | 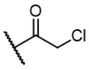 | 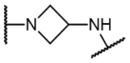 | 188 | 50.7 | 92.8 |
| XD11 | 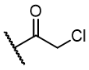 | 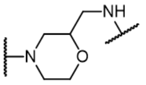 | 212 | 64.9 | 99.1 |

[a]Average inhibition rate from two independent measurements

#### Table S9. SARS-CoV-2 PLpro Inhibition Rate of Compounds.

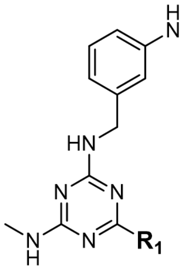


| Cmpd. | $R_1$ | Inhibition rate(%)[a] | |
|---|---|---|---|
| | | 20 μM | 100 μM |
| **XD12** | 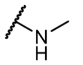 | 21.3 | 31.5 |
| **XD13** | 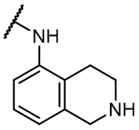 | 38.3 | 63.9 |
| **XD14** | 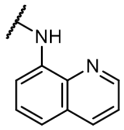 | 14.4 | 46.7 |
| **XD15** | 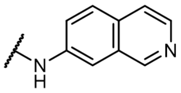 | 13.3 | 36.8 |
| **XD16** | 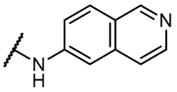 | 16.8 | 33.2 |

[a]Average inhibition rate from two independent measurements

**Table S10. Data and refinement statistics of SARS-CoV 2 3CL$^{pro}$ (8Z46)**

| Property | Value | Source |
|---|---|---|
| Space group | C 1 2 1 | Depositor |
| Cell constants<br>a, b, c, $\alpha$, $\beta$, $\gamma$ | 97.89Å 81.99Å 51.77Å<br>90.00° 114.92° 90.00° | Depositor |
| Resolution(Å) | 42.38 – 1.57<br>60.23 – 1.57 | Depositor<br>EDS |
| % Data completeness<br>(in resolution range) | 99.8 (42.38-1.57)<br>99.9 (60.23-1.57) | Depositor<br>EDS |
| $R_{merge}$ | 0.05 | Depositor |
| $R_{sym}$ | (Not available) | Depositor |
| $<I/\sigma(I)>$ | 1.96 (at1.57Å) | Xtriage |
| Refinement program | PHENIX (1.20.1_4487) | Depositor |
| R,$R_{free}$ | 0.196, 0.221<br>0.196, 0.222 | Depositor<br>DCC |
| $R_{free}$ test set | 2547 reflections (4.93%) | wwPDB-VP |
| Wilson B-factor(Å$^2$) | 22.2 | Xtriage |
| Anisotropy | 0.436 | Xtriage |
| Bulk solvent $k_{sol}$ (e/Å$^3$),<br>$B_{sol}$(Å$^2$) | 0.36, 41.6 | EDS |
| L-test for twinning | $<\|L\|>$ =0.51,$<L^2>$=0.34 | Xtriage |
| Estimated twinning fraction | No twinning to report. | Xtriage |
| $F_o$,$F_c$ correlation | 0.96 | EDS |
| Total number of atoms | 2475 | wwPDB-VP |
| Average B,all atoms (Å$^2$) | 31.0 | wwPDB-VP |

**Table S11. Data and refinement statistics of SARS-CoV 2 PL$^{pro}$ (8Z4W)**

| Property | Value | Source |
|---|---|---|
| Space group | P 1 | Depositor |
| Cell constants<br>a, b, c, $\alpha$, $\beta$, $\gamma$ | 58.23Å 75.61Å 88.35Å<br>106.05° 105.94° 105.14° | Depositor |
| Resolution(Å) | 33.52 – 2.33<br>78.75 – 2.33 | Depositor<br>EDS |
| % Data completeness<br>(in resolution range) | 97.1 (33.52-2.33)<br>97.1 (78.75-1.57) | Depositor<br>EDS |
| $R_{merge}$ | 0.05 | Depositor |
| $R_{sym}$ | (Not available) | Depositor |
| $\langle I/\sigma(I)\rangle$ | 2.03 (at2.32Å) | Xtriage |
| Refinement program | PHENIX (1.17.1_3660:? ? ? ) | Depositor |
| R,$R_{free}$ | 0.192, 0.235<br>0.193, 0.235 | Depositor<br>DCC |
| $R_{free}$ test set | 2500 reflections (4.64%) | wwPDB-VP |
| Wilson B-factor(Å$^2$) | 24.1 | Xtriage |
| Anisotropy | 0.267 | Xtriage |
| Bulk solvent $k_{sol}$ (e/Å$^3$),<br>$B_{sol}$(Å$^2$) | 0.37, 39.8 | EDS |
| L-test for twinning | $\langle\|L\|\rangle$ =0.48,$\langle L^2\rangle$=0.32 | Xtriage |
| Estimated twinning fraction | No twinning to report. | Xtriage |
| $F_o$,$F_c$ correlation | 0.93 | EDS |
| Total number of atoms | 9848 | wwPDB-VP |
| Average B,all atoms (Å$^2$) | 25.0 | wwPDB-VP |

#### Chemical Experimental Information

Starting materials were either commercially available or can be prepared according to reported literature procedures. Solvents and reagents were purchased from commercial suppliers and used as received. Reactions were monitored by thin-layer chromatography (TLC) or liquid chromatography−mass spectrometry (LC-MS). The TLC was performed on glass or aluminum-backed 60 silica plates coated with UV254 fluorescent indicator. Spots were visualized using UV light (254 or 365 nm) or an alkaline KMnO4 solution, followed by gentle heating. The LC-MS analysis was performed on a Waters Acquity UPLC instrument equipped with a CSH C18 column (50 mm × 2.1 mm, 1.7 μm packing diameter) and a Waters micro mass ZQ MS using alternate-scan positive and negative electrospray. Analytes were detected as a summed UV wavelength of 210−350 nm. Two liquid phase methods were used: Formic-40 °C, 1 mL/min flow rate. Gradient elution with the mobile phases as (A) H2O containing 0.1% volume/volume (v/v) formic acid and (B) acetonitrile containing 0.1% (v/v) formic acid. High pH-40 °C, 1 mL/min flow rate. Gradient elution with the mobile phases as (A) 10 mM aqueous ammonium bicarbonate solution, adjusted to pH 10 with 0.88 M aqueous ammonia and (B) acetonitrile. Flash column chromatography was performed using Biotage SP4 or Isolera One apparatus with SNAP silica cartridges. No unexpected or unusually high safety hazards were encountered. NMR spectra were recorded at ambient temperature (unless otherwise stated) using standard pulse methods on any of the following spectrometers and signal frequencies: Bruker AV-400 (1 H = 400 MHz, 13C = 101 MHz,), Bruker AV-500 (1 H = 500 MHz, 13C = 126 MHz), Bruker AV-600 (1 H = 600 MHz, 13C = 150 MHz) or Bruker AV4 700 MHz spectrometer (1 H = 700 MHz, 13C = 176 MHz). Chemical shifts are referenced to tetramethylsilane (TMS) or the residual solvent peak and are reported in parts per million. Coupling constants are quoted to the nearest 0.1 Hz, and multiplicities are given by the following abbreviations and combinations thereof: s (singlet), δ (doublet), t (triplet), q (quartet), quin (quintet), sxt (sextet), m (multiplet), br. (broad). The purity of synthesized compounds was determined by an LC-MS analysis. All compounds for biological testing were more than 95% pure.

**General procedure for synthesis of intermediates S1, S2, S3, S4, S5**

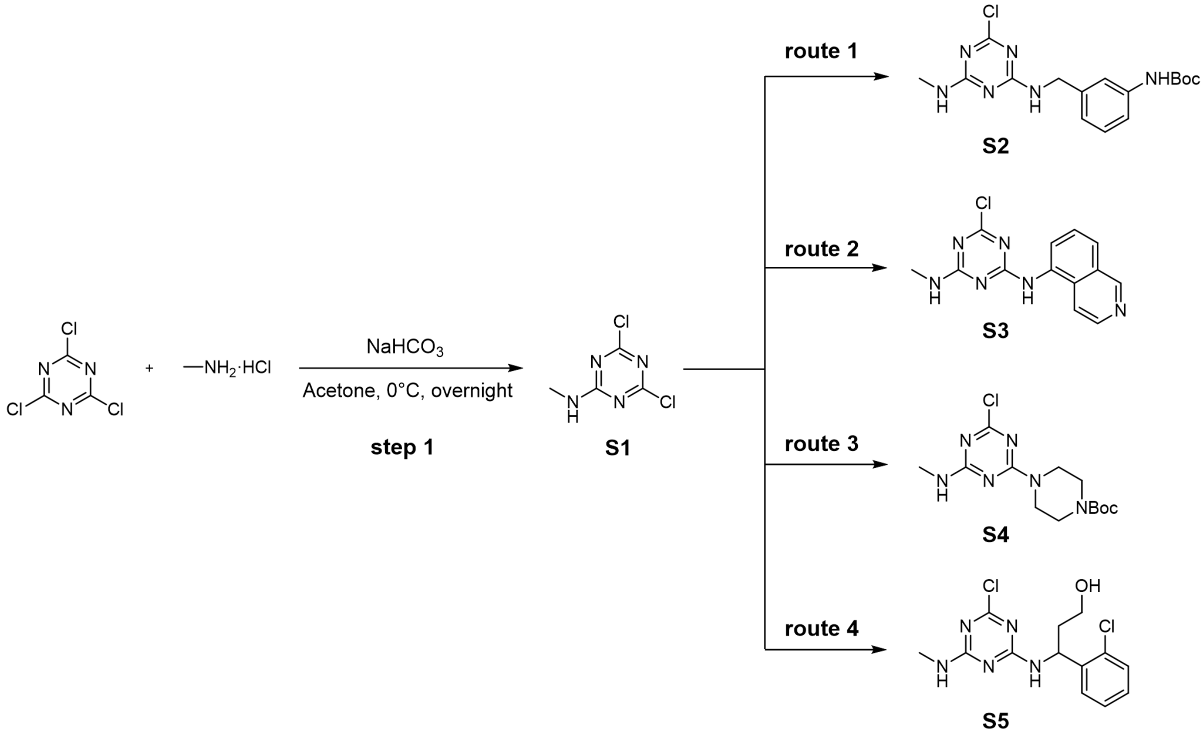


**Step 1:** To a solution of 2,4,6-trichloro-1,3,5-triazine (5.0 g, 27.1 mmol) and methylamine hydrochloride (1.83 g, 27.1 mmol) in Acetone (40 mL) was added $NaHCO_3$ (6.83 g, 81.3mmol) at 0 °C. The reaction mixture was stirred at 0°C over 12h. LC-MS indicated that no starting materials remained. The purification by flash column chromatography ( Hexenes/EtOAc, 10:1) afforded the **S1** (4.5 g, yield 93%). **$^1$H NMR** (400 MHz, DMSO) δ 9.02 – 8.95 (m, 1H), 2.82 (d, J = 4.7 Hz, 3H). **$^{13}$C NMR** (101 MHz, DMSO) δ 169.92, 168.72, 166.02, 165.98, 28.07, 28.04. **LRMS** (ESI) calcd for $[M+H]^+$ 179.0, found 179.0.

**Synthetic route 1:** To a solution of **S1** (1.5 g, 8.4 mmol) and tert-butyl (3-(aminomethyl)phenyl)-carbamate (1.87 g, 8.4 mmol) in ACN (20 mL) was added DIPEA (3.24 g, 25.1 mmol) at room temperature. The reaction mixture was stirred at 25°C over 12h. LC-MS indicated that no starting materials remained. The purification by flash column chromatography ( Hexenes/EtOAc, 3:1) afforded the **S2** (2.8 g, yield 91%). **$^1$H NMR** (400 MHz, DMSO) δ 7.51 – 7.35 (m, 1H), 7.29 (t, *J* = 10.1 Hz, 1H), 7.18 (t, *J* = 7.9 Hz, 1H), 6.95 – 6.84 (m, 1H), 4.40 (dd, *J* = 8.1, 4.7 Hz, 2H), 2.74 (dd, *J* = 9.3, 4.0 Hz, 3H), 1.46 (s, 9H), 1.23 (s, 1H). **$^{13}$C NMR** (101 MHz, DMSO) δ 166.18, 153.25, 140.39, 139.98, 128.96, 121.81, 121.09, 117.78, 117.12, 79.43, 44.25, 40.53, 40.32, 40.11, 39.90, 39.69, 39.48, 39.27, 28.57, 27.70. **LRMS** (ESI) calcd for $[M-56+H]^+$ 309.1, found 309.2.

**Synthetic route 2:** To a solution of **S1** (1.5 g, 8.4 mmol) and isoquinolin-5-amine (1.81 g, 12.6 mmol) in ACN (20 mL) was added DIPEA (3.24 g, 25.1 mmol) at room temperature. The reaction mixture was stirred at 25°C over 16h. LC-MS indicated that no starting materials remained. After adding cold water, the crude product was precipitated and filtered. The crude product was washed with acetonitrile (5 mL×3) without further purification to afford **S3**. **LRMS** (ESI) calcd for $[M+H]^+$ 287.1, found 287.1.

**Synthetic route 3:** To a solution of **S1** (1.5 g, 8.4 mmol) and tert-butyl piperazine-1-carboxylate (1.56 g, 8.4 mmol) in ACN (20 mL) was added DIPEA (3.24 g, 25.1 mmol) at room temperature. The reaction mixture was stirred at 25°C over 12h. LC-MS indicated that no starting materials remained. The purification by flash column chromatography ( Hexenes/EtOAc, 5:1) afforded the **S4** (2.6 g, yield 94%). **$^1$H NMR** (400 MHz, CDCl3) δ 6.83 (s, 1H), 3.88 – 3.70 (m, 4H), 3.46 (ddt, J = 11.9, 8.1, 4.9 Hz, 4H), 2.97 (d, J = 4.9 Hz, 3H), 1.48 (d, J = 2.1 Hz, 9H). **$^{13}$C NMR** (101 MHz, $CDCl_3$) δ 168.54, 165.98, 164.52, 154.66, 80.23, 77.40, 77.08, 76.76, 43.29, 28.39, 27.71. **LRMS** (ESI) calcd for $[M-56+H]^+$ 273.1, found 273.2.

**Synthetic route 4:** To a solution of **S1** (1.5 g, 8.4 mmol) and 3-amino-3-(2-chlorophenyl)propan -1-ol (1.56 g, 8.4 mmol) in ACN (10 mL) was added DIEA (3.24 g, 25.1 mmol) at room temperature. The reaction mixture was stirred at 25°C over 12h. LC-MS indicated that no starting materials remained. The purification by flash column chromatography ( Hexenes/EtOAc, 1:1) afforded the **S5** (2.2 g, yield 80%). **$^1$H NMR** (400 MHz, DMSO) δ 8.61 – 8.29 (m, 1H), 7.68 (q, $J$ = 4.7 Hz, 1H), 7.52–7.44 (m, 1H), 7.40 (dt, $J$ = 8.0, 1.7 Hz, 1H), 7.35–7.21 (m, 2H), 5.57 – 5.40 (m, 1H), 4.63 (dt, $J$ = 9.6, 4.8 Hz, 1H), 3.52 (dq, $J$ = 10.5, 4.8 Hz, 2H), 2.82 – 2.67 (m, 3H), 1.95 – 1.78 (m, 2H). **$^{13}$C NMR** (101 MHz, DMSO) δ 168.78, 168.44, 168.02, 166.19, 166.04, 166.01, 165.87, 165.40, 165.08, 142.43, 142.20, 141.89, 132.18, 132.01, 131.98, 129.76, 129.68, 129.42, 128.74, 128.71, 128.63, 127.96, 127.92, 127.89, 127.80, 58.22, 58.13, 49.15, 49.03, 48.94, 40.56, 40.35, 40.15, 39.94, 39.73, 39.52, 39.31, 38.66, 38.42, 38.36, 27.80, 27.71, 27.40. **LRMS** (ESI) calcd for $[M+H]^+$ 328.1, found 328.1.

**General procedure for synthesis of copound XD1, XD2, XD5, XD7, XD8, XD10, XD11.**

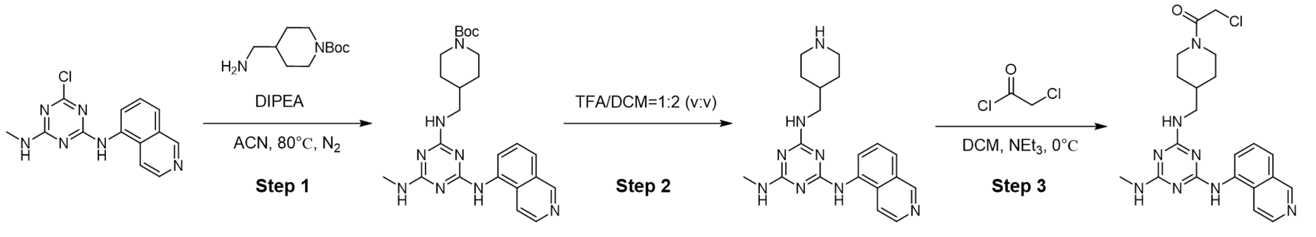


**Step 1:** To a solution of **S3** (150 mg, 0.52 mmol) and tert-butyl 4-(aminomethyl)-piperidine-1-carboxylate (214 mg, 0.78 mmol) in ACN (6 mL) was added DIPEA (203 mg, 1.57 mmol) at 80 °C. The reaction mixture was stirred at $N_2$ atmosphere for 4h. LC-MS indicated that no starting materials remained. After adding cold water, the crude product was precipitated and filtered off. The purification by flash column chromatography (DCM/MeOH, 20:1) afforded the **S3-1** (221 mg, yield 91%) as a white solid. **$^1$H NMR** (400 MHz, $CD_3OD$_SPE) δ 7.64 (s, 1H), 7.06 (t, $J$ = 7.7 Hz, 1H), 6.92 (s, 1H), 6.78 (t, $J$ = 7.8 Hz, 2H), 6.22 (d, $J$ = 8.8 Hz, 1H), 4.13 – 4.00 (m, 2H), 3.37 (s, 3H), 2.95 (s, 2H), 2.74 (s, 2H), 1.75 (s, 2H), 1.46 (s, 9H), 1.30 (d, $J$ = 3.8 Hz, 1H), 1.16 (d, $J$ = 15.1 Hz, 2H). **$^{13}$C NMR** (101 MHz, $CD_3OD$_SPE) δ 155.14, 141.83, 128.60, 126.57, 122.65, 117.62, 115.61, 79.52, 48.48, 48.27, 48.05, 47.84, 47.63, 47.41, 47.20, 46.99, 45.66, 36.54, 29.67, 27.34, 26.40. **LRMS** (ESI) calcd for $[M+H]^+$ 465.3, found 465.2.

**Step 2:** Compound **S3-1** (221 mg, 0.48 mmol) was subjected to N-Boc deprotection procedure with TFA (2mL) and DCM (4 mL). The reaction mixture was stirred for 2h. The purification by Prep-HPLC afforded the **XD1** (166 mg, yield 96%) as a white solid. **HRMS** (ESI) calcd for $[M+H]^+$ 365.2197, found 365.2198.

**Step 3:** To a solution of **XD1** (166 mg, 0.46 mmol) and $NEt_3$ (138 mg, 1.37 mmol) in DCM (4 mL) was added 2-chloroacetyl chloride (203 mg, 1.57 mmol, in 2mL DCM ) slowly at 0°C. The reaction mixture was stirred for 4h. The purification by Prep-HPLC afforded the **XD5** (93 mg, yield 46%) as a white solid. **$^1$H NMR** (500 MHz, DMSO) δ 10.71 – 10.45 (m, 1H), 9.51 – 9.20 (m, 1H), 8.74 (tt, $J$ = 16.6, 7.2 Hz, 1H), 8.44 – 8.08 (m, 2H), 7.96 – 7.78 (m, 2H), 7.41 (d, $J$ = 7.7 Hz, 1H), 4.38 – 4.27 (m, 3H), 3.50 – 3.22 (m, 2H), 3.02 (t, $J$ = 8.2 Hz, 2H), 2.91 (dd, $J$ = 15.5, 4.7 Hz, 3H), 2.63 (d, $J$ = 12.1 Hz, 1H), 1.92 – 1.73 (m, 4H), 1.30 – 1.04 (m, 3H). **$^{13}$C NMR** (126 MHz, DMSO) δ 166.80,

166.29, 165.96, 165.70, 164.81, 162.25, 162.01, 158.91, 158.66, 146.44, 146.25, 145.99, 145.95, 133.13, 133.01, 128.08, 127.97, 127.42, 127.34, 126.84, 126.63, 120.99, 119.56, 119.33, 119.22, 118.47, 116.11, 46.06, 45.99, 45.80, 42.49, 42.05, 36.27, 36.08, 35.71, 30.40, 29.71, 29.59, 28.08, 27.96. **HRMS** (ESI) calcd for $[M+H]^+$ 441.1914, found 441.1913.

N2-(3-aminobenzyl)-N4-(isoquinolin-5-yl)-N6-methyl-1,3,5-triazine-2,4,6-triamine (**XD2**)

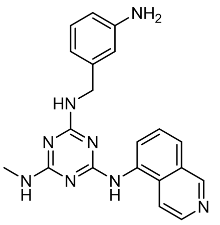

The preparation was described as above procedure to provide compound **XD2** by Prep-HPLC as solid (153 mg, yield 82%). **$^1$H NMR** (500 MHz, DMSO) δ 10.71 – 10.43 (m, 1H), 9.52 – 9.18 (m, 1H), 8.96 – 8.61 (m, 2H), 8.33 – 8.14 (m, 1H), 8.02 – 7.79 (m, 2H), 7.44 – 7.40 (m, 1H), 7.37 – 7.30 (m, 1H), 7.25 – 7.09 (m, 2H), 7.05 (dt, *J* = 13.5, 4.0 Hz, 1H), 4.83 – 4.55 (m, 2H), 3.14 – 2.69 (m, 3H). **$^{13}$C NMR** (126 MHz, DMSO) δ 166.72, 166.29, 166.13, 165.76, 162.18, 158.97, 158.70, 146.01, 141.02, 133.05, 129.97, 128.09, 127.34, 126.66, 121.01, 119.64, 119.41, 119.26, 118.25, 28.12, 28.06, 27.98, 7.89. **HRMS** (ESI) calcd for $[M+H]^+$ 373.1884, found 373.1886.

2-chloro-N-(3-(((4-(isoquinolin-5-ylamino)-6-(methylamino)-1,3,5-triazin-2-yl)amino)methyl)phenyl)acetamide (**XD7**)

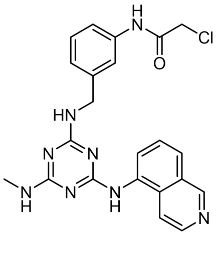

The preparation was described as above procedure to provide compound **XD7** by Prep-HPLC as solid (73 mg, yield 41%). **HRMS** (ESI) calcd for $[M+H]^+$ 449.1600, found 449.1602.

2-chloro-N-(1-(4-(isoquinolin-5-ylamino)-6-(methylamino)-1,3,5-triazin-2-yl)piperidin-4-yl)acetamide **(XD8)**

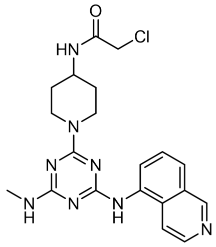


The preparation was described as above procedure to provide compound **XD7** by Prep-HPLC as solid (66 mg, yield 72%). **$^1$H NMR** (500 MHz, DMSO) δ 10.77 – 10.58 (m, 1H), 9.58 – 9.35 (m, 1H), 8.67 (dd, *J* = 16.0, 7.5 Hz, 1H), 8.35 (s, 1H), 8.26 – 8.15 (m, 1H), 8.07 – 7.94 (m, 1H), 7.83 (t, *J* = 7.9 Hz, 1H), 7.40 (d, *J* = 7.9 Hz, 1H), 4.94 – 4.47 (m, 3H), 4.07 (d, *J* = 3.7 Hz, 1H), 3.03 – 2.92 (m, 4H), 1.90 (s, 2H), 1.46 – 1.16 (m, 4H). **HRMS** (ESI) calcd for $[M+H]^+$ 427.1756, found 427.1754

2-chloro-N-(1-(4-(isoquinolin-5-ylamino)-6-(methylamino)-1,3,5-triazin-2-yl)azetidin-3-yl)acetamide (**XD10**)

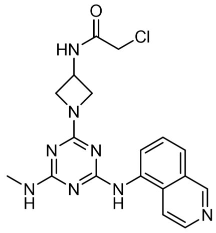


The preparation was described as above procedure to provide compound **XD10** by Prep-HPLC as solid (43mg, yield 38%). **$^1$H NMR** (500 MHz, DMSO) δ 10.73 – 10.47 (m, 1H), 9.47 – 9.24 (m, 1H), 9.17 (d, J = 6.6 Hz, 1H), 8.75 – 8.65 (m, 1H), 8.39 – 8.25 (m, 1H), 8.06 – 7.87 (m, 1H), 7.81 (td, J = 8.0, 1.8 Hz, 1H), 7.40 (d, J = 7.8 Hz, 1H), 4.65 (s, 2H), 4.55 (td, J = 8.7, 4.3 Hz, 2H), 4.49 – 4.34 (m, 1H), 4.14 (d, J = 1.6 Hz, 2H), 3.02 – 2.88 (m, 3H). **$^{13}$C NMR** (126 MHz, DMSO) δ 166.66, 165.28, 164.84, 161.83, 158.64, 146.61, 145.96, 132.96, 128.19, 127.04, 121.00, 119.56, 56.97, 42.95, 40.37, 40.21, 40.04, 39.87, 39.71, 39.54, 39.37, 27.99. **LRMS** (ESI) calcd for $[M+H]^+$ 399.1, found 399.2

2-chloro-N-((4-(4-(isoquinolin-5-ylamino)-6-(methylamino)-1,3,5-triazin-2-yl)morpholin-2-yl)methyl)acetamide (**XD11**)

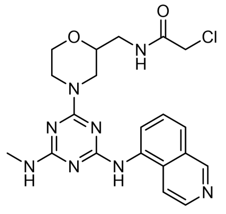

The preparation was described as above procedure to provide compound **XD11** by Prep-HPLC as solid (50mg, yield 48%). **$^1$H NMR** (500 MHz, DMSO) δ 10.78 – 10.55 (m, 1H), 9.58 – 9.31 (m, 1H), 8.76 – 8.63 (m, 1H), 8.55 – 8.49 (m, 1H), 8.35 – 8.19 (m, 1H), 8.07 – 7.77 (m, 2H), 7.40 (dd, *J* = 7.8, 3.8 Hz, 1H), 4.85 – 4.41 (m, 2H), 4.16 – 4.06 (m, 2H), 4.02 (d, *J* = 11.6 Hz, 1H), 3.63 – 3.25 (m, 4H), 3.23 – 3.02 (m, 2H), 3.00 – 2.86 (m, 3H). **$^{13}$C NMR** (126 MHz, DMSO) δ 168.99, 166.79, 166.06, 164.82, 164.37, 146.51, 133.00, 127.39, 122.41, 120.92, 119.26, 75.06, 74.10, 66.19, 51.98, 43.00, 41.67. **HRMS** (ESI) calcd for $[M+H]^+$ 443.1705, found 443.1703

**General procedure for one-pot synthesis of copound XD3, XD4, XD6, XD9.**

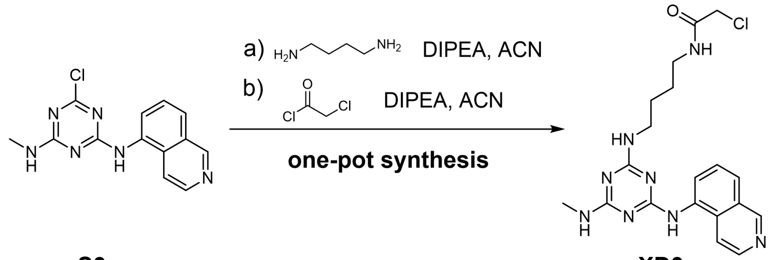


To a solution of **S3** (120 mg, 0.42 mmol) and butane-1,4-diamine (55 mg, 0.63 mmol) in ACN (4 mL) was added DIPEA (163 mg, 1.26 mmol) at 80 °C. The reaction mixture was stirred at 80 °C in $N_2$ atmosphere for 4h. LC-MS indicated that no starting materials remained. Cooling the reaction to room temperature followed by an ice bath and then added 2-chloroacetyl chloride (47 mg, 0.42 mmol). The purification by Prep-HPLC afforded the **XD3** (68 mg, yield 39%) as a white solid. **$^1$H NMR** (500 MHz, MeOD) δ 10.77 – 10.45 (m, 1H), 9.58 – 9.33 (m, 1H), 8.63 – 8.50 (m, 1H), 7.98 – 7.73 (m, 2H), 7.49 – 7.39 (m, 1H), 4.04 (d, *J* = 9.1 Hz, 2H), 3.65 – 3.34 (m, 3H), 3.19 – 2.88 (m, 4H), 1.75 – 1.62 (m, 4H). **$^{13}$C NMR** (126 MHz, MeOD) δ 167.98, 166.51, 166.17, 145.45, 132.54, 127.85, 127.57, 126.15, 126.04, 120.02, 119.64, 119.49, 119.41, 119.36, 48.14, 47.97, 47.92, 47.80, 47.63, 47.46, 47.29, 47.12, 41.84, 40.20, 40.05, 39.93, 39.07, 38.84, 26.70, 26.67, 26.29, 26.19, 26.08. **HRMS** (ESI) calcd for $[M+H]^+$ 415.1756, found 415.1755

2-chloro-N-(2-((4-(isoquinolin-5-ylamino)-6-(methylamino)-1,3,5-triazin-2-yl)(methyl)amino)ethyl) -N-methylacetamide (**XD4**)

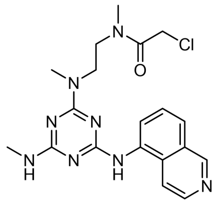

The preparation was described as above procedure to provide compound **XD4** by Prep-HPLC as solid (42 mg, yield 24%). **$^1$H NMR** (500 MHz, MeOD) δ 10.90 – 10.56 (m, 1H), 9.55 – 9.40 (m, 1H), 8.64 – 8.52 (m, 1H), 8.08 – 7.75 (m, 2H), 7.44 (q, *J* = 6.0 Hz, 1H), 4.32 – 4.15 (m, 2H), 4.02 – 3.87 (m, 2H), 3.79 – 3.65 (m, 2H), 3.34 (dd, *J* = 15.9, 4.0 Hz, 3H), 3.19 – 3.01 (m, 6H). **$^{13}$C NMR** (126 MHz, MeOD) δ 167.71, 146.13, 145.33, 132.56, 132.37, 128.05, 126.14, 125.93, 120.15, 120.09, 120.01, 119.57, 48.16, 47.99, 47.82, 47.65, 47.48, 47.30, 47.13, 45.92, 45.64, 45.32, 41.45, 40.86, 35.65, 35.35, 34.25, 26.82, 26.68. **HRMS** (ESI) calcd for $[M+H]^+$ 415.1756, found 415.1759

2-chloro-1-(4-(4-(isoquinolin-5-ylamino)-6-(methylamino)-1,3,5-triazin-2-yl)piperazin-1-yl)ethan-1-one **(XD6)**

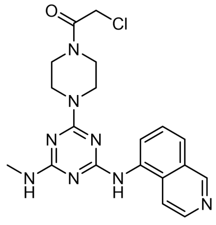

The preparation was described as above procedure to provide compound **XD6** by Prep-HPLC as solid (50 mg, yield 29%). **$^1$H NMR** (500 MHz, DMSO) δ 10.81 – 10.59 (m, 1H), 9.60 – 9.35 (m, 1H), 8.68 (dd, J = 14.6, 7.3 Hz, 1H), 8.28 (dt, J = 15.6, 7.6 Hz, 1H), 8.08 – 7.91 (m, 1H), 7.84 (td, J = 8.0, 1.8 Hz, 1H), 7.41 (d, J = 7.8 Hz, 1H), 4.49 (t, J = 6.4 Hz, 2H), 4.12 – 3.90 (m, 4H), 3.63 (s, 3H), 3.07 – 2.92 (m, 4H). **$^{13}$C NMR** (126 MHz, DMSO) δ 165.45, 158.57, 146.66, 145.90, 137.73, 133.06, 128.15, 127.42, 120.89, 119.29, 85.34, 61.78, 42.48, 40.43, 40.35, 40.26, 40.18, 40.09, 39.93, 39.76, 39.59, 39.43, 28.05. **HRMS** (ESI) calcd for $[M+H]^+$ 413.1600, found 413.1601

2-chloro-1-(6-(4-(isoquinolin-5-ylamino)-6-(methylamino)-1,3,5-triazin-2-yl)-2,6-diazaspiro[3.3]heptan-2-yl)ethan-1-one (**XD9**)

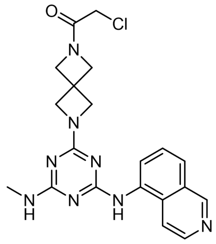

The preparation was described as above procedure to provide compound **XD6** by Prep-HPLC as solid (62 mg, yield 35%). $^{1}$**H NMR** (500 MHz, DMSO) δ 10.65 – 10.45 (m, 1H), 9.41 – 9.19 (m, 1H), 8.84 – 8.64 (m, 1H), 8.44 – 8.14 (m, 1H), 8.03 – 7.85 (m, 1H), 7.81 (td, *J* = 7.9, 1.6 Hz, 1H), 7.40 (d, *J* = 7.7 Hz, 1H), 4.75 – 4.33 (m, 4H), 4.37 – 4.15 (m, 4H), 4.14 (d, *J* = 2.6 Hz, 2H), 3.12 – 2.76 (m, 3H). $^{13}$**C NMR** (126 MHz, DMSO) δ 166.36, 166.03, 165.79, 165.01, 164.62, 162.11, 161.75, 146.47, 145.96, 133.11, 128.15, 128.02, 127.32, 126.84, 126.56, 121.04, 119.32, 60.37, 59.82, 58.64, 40.57, 40.55, 33.60, 27.96. **HRMS** (ESI) calcd for $[M+H]^+$ 425.1600, found 425.1600

**General procedure for synthesis of copound XD13.**

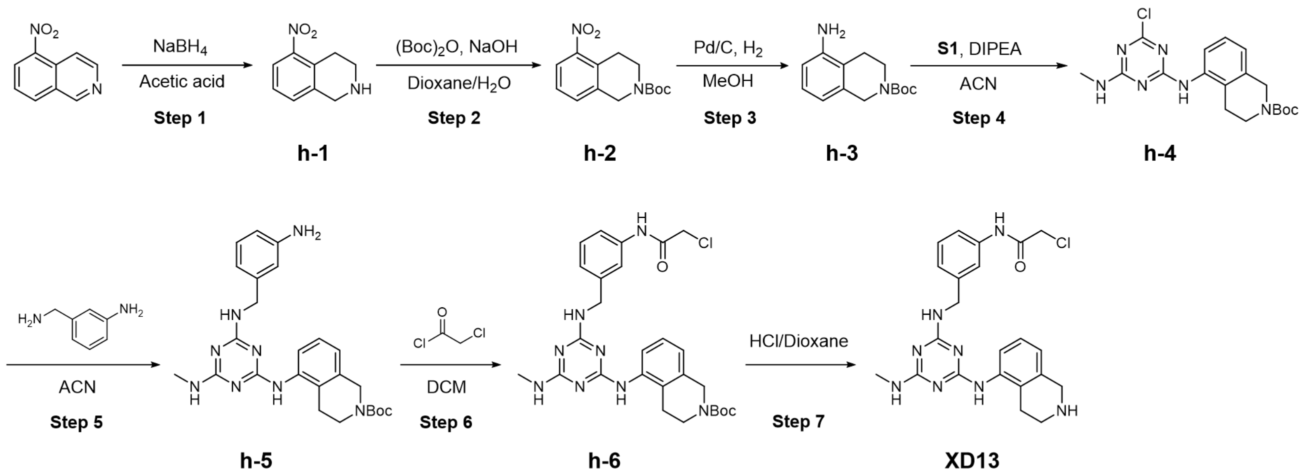


**Step 1:** To a solution of 5-nitroisoquinoline (500 mg, 2.85 mmol) in acetic acid (25 mL) was added small portions of $NaBH_4$ until TLC examination of the reaction mixture indicating the absence of the starting material. The solution was poured onto ice, basified with NaOH and then extracted three times with $CH_2Cl_2$. The organic layer was washed twice with water, dried over sodium sulfate and evaporated. The purification by flash column chromatography ( Hexenes/EtOAc, 1:1) afforded the **h-1** (425 mg, yield 84%). $^{1}$**H NMR** (400 MHz, $CDCl_3$) δ 7.79 (dd, *J* = 7.7, 1.7 Hz, 1H), 7.32 – 7.20 (m, 2H), 4.09 (s, 2H), 3.16 (t, *J* = 6.0 Hz, 2H), 3.08 (t, *J* = 6.0 Hz, 2H). **LRMS** (ESI) calcd for $[M+H]^+$ 179.1, found 179.1.

**Step 2:** To a mixture of **h-1** (425 mg, 2.39 mmol) in dioxane (5 mL) and water (5 mL) was added NaOH (115 mg, 2.84 mmol) followed by di-tert-butyl dicarbonate (1.04 g, 4.78 mmol). The reaction mixture was stirred at room temperature for 6 h. The solvent was removed by rotary evaporation, and 3 N aqueous hydrochloric acid (6 mL) was added dropwise to the residue. A precipitate was obtained, collected, washed with EtOAc, and dried to provide **h-2** (650 mg, yield 98%). **$^1$H NMR** (400 MHz, $CDCl_3$) δ 7.81 (d, $J$ = 7.7 Hz, 1H), 7.41 – 7.30 (m, 2H), 4.66 (s, 2H), 3.64 (t, $J$ = 5.9 Hz, 2H), 3.10 (t, $J$ = 5.9 Hz, 2H), 1.50 (d, $J$ = 1.4 Hz, 9H). **LRMS** (ESI) calcd for $[M\text{-}56+H]^+$ 223.1, found 223.1.

**Step 3:** The compound **h-2** (650mg, 2.3 mmol) and Pd/C (10%, 65 mg) were suspended on methanol (30 mL). The system was vacuumed, replaced by hydrogen and reacted at room temperature for 12 hours. The reaction mixture was then diluted with MeOH (30 mL), filtered through celite, and concentrated in vacuo. The purification by flash column chromatography (Hexanes/EtOAc, 1:1) afforded the **h-3** (432 mg, 74%). **$^1$H NMR** (400 MHz, $CDCl_3$) δ 7.81 (d, $J$ = 7.7 Hz, 1H), 7.41 – 7.30 (m, 2H), 4.66 (s, 2H), 3.64 (t, $J$ = 5.9 Hz, 2H), 3.10 (t, $J$ = 5.9 Hz, 2H), 1.50 (d, $J$ = 1.4 Hz, 9H). **LRMS** (ESI) calcd for $[M+H]^+$ 249.2, found 249.1.

**Step 4:** To a solution of **h-3** (180 mg, 0.73 mmol) and **S1** (130 mg, 0.73 mmol) in ACN (6 mL) was added DIPEA (283 mg, 2.19 mmol) at 90 °C. The reaction mixture was stirred at 90 °C in $N_2$ atmosphere for 2h. LC-MS indicated that no starting materials remained. Cooling the reaction to room temperature followed by an ice bath and the crude product was precipitated and filtered. The purification by flash column chromatography (Hexanes/EtOAc, 1:1) afforded the **h-4** (243mg, 86%). **$^1$H NMR** (400 MHz, DMSO) δ 9.53 – 9.26 (m, 1H), 7.99 – 7.73 (m, 1H), 7.34 – 6.98 (m, 3H), 4.51 (s, 2H), 3.51 (d, $J$ = 5.2 Hz, 2H), 2.83 – 2.56 (m, 5H), 1.43 (s, 9H). **$^{13}$C NMR** (101 MHz, DMSO) δ 166.43, 166.30, 164.96, 154.40, 136.34, 136.29, 131.53, 126.44, 126.19, 125.26, 124.50, 79.43, 60.23, 40.58, 40.37, 40.17, 39.96, 39.75, 39.54, 39.33, 28.56, 28.54, 27.72, 24.91, 21.21, 14.54. **LRMS** (ESI) calcd for $[M+H]^+$ 391.2, found 391.2

**Step 5:** To a solution of **h-4** (243 mg, 0.62 mmol) and 3-(aminomethyl)aniline (76 mg, 0.62 mmol) in ACN (6 mL) was added DIPEA (240 mg, 1.86 mmol) at 90 °C. The reaction mixture was stirred at

90 °C in $N_2$ atmosphere for 4h. LC-MS indicated that no starting materials remained. Cooling the reaction to room temperature followed by an ice bath and the crude product was precipitated and filtered. The crude product was washed with acetonitrile (3 mL×3) without further purification to afford **h-5** (280 mg, 94%). **$^1$H NMR** (400 MHz, $CD_3OD$_SPE) δ 7.50 – 7.25 (m, 1H), 7.08 (t, *J* = 7.8 Hz, 1H), 6.99 (t, *J* = 7.7 Hz, 1H), 6.87 (d, *J* = 7.6 Hz, 1H), 6.70 – 6.47 (m, 3H), 4.53 – 4.24 (m, 4H), 3.51 (t, *J* = 6.0 Hz, 2H), 2.79 (s, 3H), 2.62 (s, 2H), 1.46 (s, 9H). **$^{13}$C NMR** (101 MHz, $CD_3OD$_SPE) δ 165.96, 164.79, 155.15, 147.32, 140.63, 136.65, 134.01, 128.77, 125.86, 122.67, 116.72, 113.84, 79.96, 53.51, 48.33, 48.12, 47.98, 47.91, 47.70, 47.48, 47.27, 47.06, 43.83, 27.45, 27.45, 26.43, 24.34. **LRMS** (ESI) calcd for $[M+H]^+$ 477.3, found 477.2

**Step 6:** To a solution of **h-5** (280 mg, 0.59 mmol) and $NEt_3$ (179 mg, 1.76 mmol) in DMF (5 mL) was added 2-chloroacetyl chloride (66 mg, 0.59 mmol, in 1mL DMF ) slowly at 0°C. The reaction mixture was stirred for 4h. The purification by flash column chromatography (DCM/MeOH, 20:1) afforded the **h-6** (160 mg, yield 49%) as a white solid. **$^1$H NMR** (400 MHz, $CDCl_3$) δ 8.39 (s, 1H), 7.49 – 7.37 (m, 2H), 7.19 – 7.00 (m, 2H), 6.93 – 6.72 (m, 2H), 5.76 (s, 1H), 5.17 (s, 1H), 4.53 (s, 4H), 4.13 (s, 2H), 3.58 (s, 2H), 2.85 (d, *J* = 4.8 Hz, 3H), 2.65 (s, 2H), 2.45 (s, 1H), 1.47 (s, 9H), 1.30 – 1.19 (m, 1H). **$^{13}$C NMR** (101 MHz, $CDCl_3$) δ 164.11, 154.81, 140.59, 136.99, 129.26, 126.11, 124.18, 119.04, 79.92, 77.41, 77.09, 76.77, 44.31, 42.95, 28.50, 27.62, 24.61. **LRMS** (ESI) calcd for $[M+H]^+$ 553.2, found 553.3

**Step 7:** Compound **h-6** (160 mg, 0.29 mmol) was subjected to N-Boc deprotection procedure with HCl (4M in dioxane, 3mL) and dioxane (3 mL). The reaction mixture was stirred for 2h. The purification by Prep-HPLC afforded the **XD13** (119 mg, yield 91%) as a white solid. **$^1$H NMR** (400 MHz, DMSO) δ 10.38 (d, *J* = 14.0 Hz, 1H), 9.28 (s, 1H), 7.55 – 7.38 (m, 2H), 7.29 (t, *J* = 7.4 Hz, 2H), 7.21 – 6.88 (m, 2H), 4.56 – 4.37 (m, 2H), 4.35 – 4.21 (m, 4H), 3.36 (s, 3H), 2.97 – 2.86 (m, 2H), 2.82 (s, 2H). **$^{13}$C NMR** (101 MHz, DMSO) δ 165.14, 159.41, 148.10, 143.05, 139.02, 138.49, 129.26, 123.15, 118.33, 115.80, 62.07, 56.39, 44.04, 40.59, 40.38, 40.19, 40.17, 39.96, 39.75, 39.54, 39.33, 27.82, 21.54, 12.50, 8.22. **LRMS** (ESI) calcd for $[M+H]^+$ 453.2, found 453.1

**General procedure for synthesis of copound XD12, XD14-XD16.**

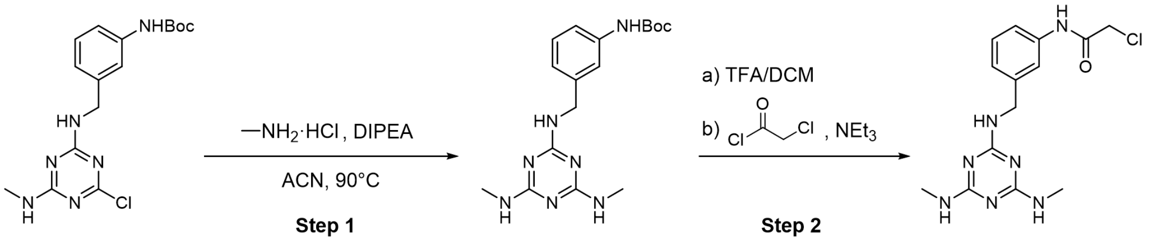


**Step 1:** To a solution of **S2** (150 mg, 0.41 mmol) and methylamine hydrochloride (56 mg, 0.82 mmol) in ACN (4 mL) was added DIPEA (159 mg, 1.23 mmol) at 90 °C. The reaction mixture was stirred at 90 °C in $N_2$ atmosphere for 4h. LC-MS indicated that no starting materials remained. Cooling the reaction to room temperature followed by an ice bath and the crude product was precipitated and filtered. The purification by flash column chromatography (Hexanes/EtOAc, 3:1) afforded the **S2-1** (135 mg, yield 91%). **$^1$H NMR** (400 MHz, DMSO) δ 9.28 (s, 1H), 7.52 – 7.37 (m, 1H), 7.28 – 7.11 (m, 2H), 6.89 (s, 1H), 6.60 – 6.22 (m, 2H), 4.58 – 4.24 (m, 2H), 3.01 (s, 1H), 2.71 (d, *J* = 13.5 Hz, 6H), 1.47 (d, *J* = 8.8 Hz, 9H). **$^{13}$C NMR** (101 MHz, DMSO) δ 166.48, 153.26, 141.99, 139.77, 135.56, 128.65, 121.14, 116.86, 79.30, 55.38, 40.60, 40.39, 40.19, 39.98, 39.77, 39.56, 39.35, 28.60, 27.58. **LRMS** (ESI) calcd for $[M+H]^+$ 360.2, found 360.2

**Step 2:** Compound **S2-1** (135 mg, 0.38 mmol) was subjected to N-Boc deprotection procedure with TFA (2mL) and DCM (4 mL). The reaction mixture was stirred for 2h. LC-MS indicated that no starting materials remained. Without further purification, the crude product was dissolved in 4 mL DCM and then added $NEt_3$ (114.0 mg, 1.14 mmol), 2-chloroacetyl chloride (42 mg, 0.38 mmol). The reaction mixture was stirred for 4h. The purification by Prep-HPLC afforded the **XD12** (64 mg, yield 50 %) as a white solid. **$^1$H NMR** (400 MHz, MeOD) δ 7.71 – 7.61 (m, 1H), 7.43 (d, *J* = 8.1 Hz, 1H), 7.29 (t, *J* = 7.4 Hz, 1H), 7.15 – 7.07 (m, 1H), 4.62 – 4.52 (m, 2H), 4.17 (s, 2H), 2.99 – 2.88 (m, 6H). **$^{13}$C NMR** (101 MHz, MeOD) δ 166.09, 139.53, 138.73, 138.15, 128.78, 123.66, 123.50, 119.22, 118.96, 48.29, 48.11, 48.08, 47.89, 47.86, 42.69, 26.38, 26.18. **LRMS** (ESI) calcd for $[M+H]^+$ 336.1, found 336.1

2-chloro-N-(3-(((4-(methylamino)-6-(quinolin-8-ylamino)-1,3,5-triazin-2-yl)amino)methyl)phenyl)acetamide (**XD14**)

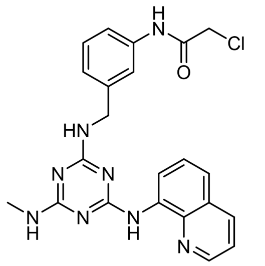

The preparation was described as above procedure to provide compound **XD14** by Prep-HPLC as solid (42 mg, yield 34%). **$^{1}$H NMR** (500 MHz, DMSO) δ 10.38 (s, 1H), 9.07 – 8.87 (m, 2H), 8.44 (dd, *J* = 15.2, 8.3 Hz, 1H), 7.72 – 7.60 (m, 3H), 7.62 – 7.44 (m, 2H), 7.33 (td, *J* = 7.9, 4.1 Hz, 1H), 7.16 – 7.07 (m, 1H), 4.61 – 4.54 (m, 2H), 4.24 (s, 2H), 4.02 (s, 2H), 2.97 – 2.87 (m, 3H), 1.23 (d, *J* = 5.5 Hz, 1H). **$^{13}$C NMR** (126 MHz, DMSO) δ 171.37, 165.15, 165.06, 159.17, 158.90, 158.64, 158.37, 149.43, 139.98, 139.14, 139.07, 138.61, 138.43, 137.29, 134.28, 134.05, 130.10, 129.38, 129.34, 129.14, 128.33, 128.29, 127.45, 127.33, 123.80, 123.15, 122.82, 122.77, 122.59, 122.25, 120.42, 119.05, 118.65, 118.45, 117.99, 117.71, 115.72, 62.31, 44.40, 44.04, 43.99, 35.58, 31.74, 29.55, 29.50, 29.28, 29.19, 29.15, 29.03, 28.13, 28.00, 27.80, 27.06, 25.57, 22.56, 14.41. **HRMS** (ESI) calcd for $[M+H]^+$ 449.1600, found 449.1600

2-chloro-N-(3-(((4-(isoquinolin-7-ylamino)-6-(methylamino)-1,3,5-triazin-2-yl)amino)methyl)pheny l)acetamide (**XD15**)

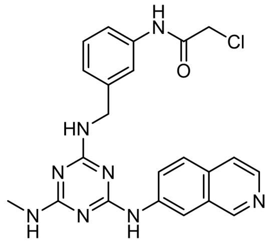

The preparation was described as above procedure to provide compound **XD15** by Prep-HPLC as solid (49 mg, yield 40%). **$^{1}$H NMR** (400 MHz, $CD_3OD$_SPE) δ 9.63 – 9.37 (m, 1H), 9.12 – 8.76 (m, 1H), 8.48 (d, *J* = 6.3 Hz, 1H), 8.40 – 8.13 (m, 4H), 7.83 (s, 1H), 7.36 (t, *J* = 7.1 Hz, 2H), 7.24 – 7.17 (m, 1H), 4.75 (s, 2H), 4.18 (d, *J* = 7.6 Hz, 2H), 3.12 – 3.01 (m, 3H), 1.38 – 1.26 (m, 1H). **$^{13}$C NMR** (126 MHz, MeOD) δ 166.15, 146.78, 140.20, 138.64, 138.26, 135.11, 131.62, 130.60, 128.94, 128.42, 127.80, 124.26, 123.14, 119.01, 117.76, 115.78, 113.51, 56.08, 55.91, 44.04, 42.65, 29.37, 26.73, 25.52, 22.33, 16.03, 15.88, 15.73. **HRMS** (ESI) calcd for $[M+H]^+$ 449.1600, found 449.1601

2-chloro-N-(3-(((4-(isoquinolin-6-ylamino)-6-(methylamino)-1,3,5-triazin-2-yl)amino)methyl)phenyl)acetamide (**XD16**)

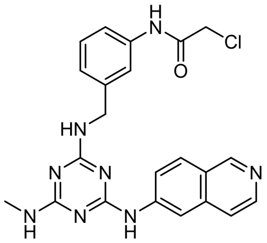

The preparation was described as above procedure to provide compound **XD16** by Prep-HPLC as solid (47 mg, yield 38%). **$^1$H NMR** (400 MHz, DMSO) δ 10.38 – 10.32 (m, 1H), 9.55 (d, *J* = 10.3 Hz, 1H), 8.52 – 8.42 (m, 1H), 8.35 – 8.27 (m, 1H), 8.23 – 8.01 (m, 2H), 7.94 (s, 1H), 7.49 (d, *J* = 8.5 Hz, 1H), 7.30 (td, *J* = 8.0, 3.8 Hz, 2H), 7.12 (d, *J* = 7.8 Hz, 1H), 4.63 – 4.54 (m, 2H), 4.23 (s, 2H), 2.93 – 2.80 (m, 3H), 1.22 (s, 1H). **$^{13}$C NMR** (126 MHz, DMSO) δ 165.14, 165.09, 163.99, 163.63, 159.39, 159.11, 158.83, 158.55, 148.15, 147.93, 147.15, 145.96, 145.51, 141.03, 140.48, 140.27, 139.99, 139.06, 132.51, 132.21, 131.51, 130.10, 129.27, 124.94, 123.63, 123.44, 123.05, 122.64, 120.06, 118.88, 118.29, 117.92, 117.72, 115.39, 113.05, 112.35, 62.31, 44.03, 43.64, 35.57, 29.50, 29.28, 29.16, 29.03, 27.96, 27.77, 27.59, 27.02, 25.57, 22.55, 14.41. **HRMS** (ESI) calcd for $[M+H]^+$ 449.1600, found 449.1599

**General procedure for synthesis of copound LU1-LU6**

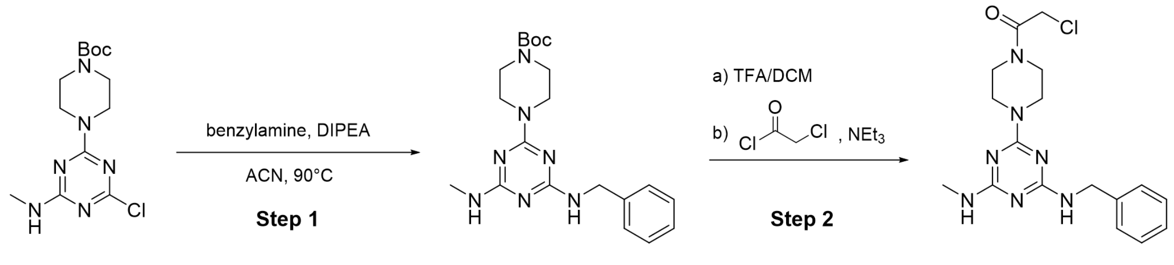


S4 S4-1 LU1

**Step 1:** To a solution of **S4** (120 mg, 0.37 mmol) and benzylamine (221 mg, 0.56 mmol) in ACN (4 mL) was added DIPEA (143 mg, 1.11 mmol) at 90 °C. The reaction mixture was stirred at 90 °C in $N_2$ atmosphere for 4h. LC-MS indicated that no starting materials remained. Cooling the reaction to room temperature followed by an ice bath and the crude product was precipitated and filtered. The purification by flash column chromatography (DCM/MeOH, 20:1) afforded the **S4-1** (129 mg, yield 88%). **$^1$H NMR** (400 MHz, $CDCl_3$) δ 7.27 – 7.14 (m, 5H), 4.48 (d, *J* = 5.6 Hz, 2H), 3.66 (s, 4H),

3.34 (t, *J* = 5.2 Hz, 4H), 3.00 (s, 1H), 2.82 (d, *J* = 4.8 Hz, 3H), 1.40 (s, 9H). **$^{13}$C NMR** (101 MHz, $CDCl_3$) δ 164.79, 154.82, 139.37, 128.48, 127.52, 127.12, 79.96, 44.69, 43.00, 29.70, 28.43, 27.56. **LRMS** (ESI) calcd for $[M+H]^+$ 400.2, found 400.2

**Step 2:** Compound **S4-1** (129 mg, 0.32 mmol) was subjected to N-Boc deprotection procedure with TFA (2mL) and DCM (4 mL). The reaction mixture was stirred for 2h. LC-MS indicated that no starting materials remained. Without further purification, the crude product was dissolved in 4 mL DCM and then added $NEt_3$ (98.9 mg, 0.97 mmol), 2-chloroacetyl chloride (36 mg, 0.32 mmol). The reaction mixture was stirred for 4h. The purification by Prep-HPLC afforded the **LU1** (78 mg, yield 64 %) as a white solid. **$^1$H NMR** (400 MHz, DMSO) δ 7.50 – 7.16 (m, 6H), 6.67 (s, 1H), 4.42 (d, *J* = 6.7 Hz, 4H), 3.79 – 3.60 (m, 4H), 3.45 (s, 4H), 2.72 (d, *J* = 4.6 Hz, 3H). **$^{13}$C NMR** (101 MHz, $CDCl_3$) δ 165.36, 164.98, 139.38, 128.51, 127.47, 127.16, 77.40, 77.08, 76.77, 46.19, 44.70, 43.08, 42.67, 42.10, 40.90, 27.60. **HRMS** (ESI) calcd for $[M+H]^+$ 376.1647, found 376.1649

2-chloro-1-(4-(4-(methylamino)-6-((1-phenylethyl)amino)-1,3,5-triazin-2-yl)piperazin-1-yl)ethan-1-one (**LU2**)

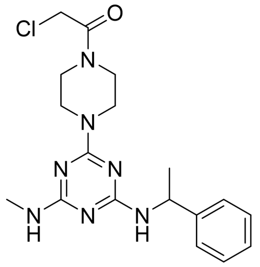

The preparation was described as above procedure to provide compound X by Prep-HPLC as solid ( mg, yield %). **$^1$H NMR** (400 MHz, $CDCl_3$) δ 7.30 (q, *J* = 8.0 Hz, 4H), 7.25 – 7.15 (m, 1H), 5.29 (s, 1H), 5.09 (s, 1H), 4.08 (s, 2H), 3.84 – 3.38 (m, 8H), 2.88 (d, *J* = 4.9 Hz, 3H), 1.49 (d, *J* = 6.9 Hz, 3H). **$^{13}$C NMR** (101 MHz, $CDCl_3$) δ 165.33, 164.77, 144.79, 128.40, 126.87, 125.98, 77.41, 77.29, 77.09, 76.77, 53.46, 46.13, 43.08, 42.68, 42.04, 40.89, 27.56, 22.79. **HRMS** (ESI) calcd for $[M+H]^+$ 390.1804, found 390.1802

2-chloro-1-(4-(4-(methylamino)-6-((1-(2-(trifluoromethyl)phenyl)ethyl)amino)-1,3,5-triazin-2-yl)pip erazin-1-yl)ethan-1-one (**LU3**)

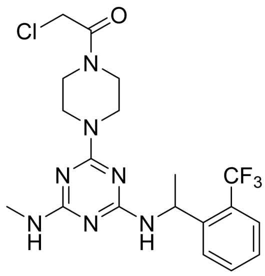

The preparation was described as above procedure to provide compound X by Prep-HPLC as solid ( mg, yield %). **$^1$H NMR** (400 MHz, $CDCl_3$) δ 7.60 (dd, *J* = 14.9, 7.9 Hz, 2H), 7.48 (t, *J* = 7.6 Hz, 1H), 7.34 – 7.26 (m, 1H), 5.42 (s, 1H), 4.08 (s, 2H), 3.93 – 3.28 (m, 8H), 3.02 (s, 1H), 2.97 – 2.71 (m, 3H), 1.47 (d, *J* = 6.8 Hz, 3H). **$^{13}$C NMR** (101 MHz, $CDCl_3$) δ 165.35, 164.67, 144.98, 132.36, 126.64, 126.38, 125.96, 125.82, 125.76, 125.70, 125.64, 123.23, 77.39, 77.28, 77.07, 76.76, 46.95, 46.04, 42.95, 42.58, 41.98, 40.87, 29.69, 27.51, 24.17. **HRMS** (ESI) calcd for $[M+H]^+$ 458.1677, found 458.1676

2-chloro-1-(4-(4-((1-(2-fluorophenyl)ethyl)amino)-6-(methylamino)-1,3,5-triazin-2-yl)piperazin-1-yl) ethan-1-one (**LU4**)

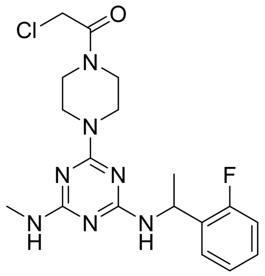

The preparation was described as above procedure to provide compound X by Prep-HPLC as solid ( mg, yield %). **$^1$H NMR** (400 MHz, $CDCl_3$) δ 7.30 (t, *J* = 7.2 Hz, 1H), 7.18 (q, *J* = 5.8 Hz, 1H), 7.09 – 6.94 (m, 2H), 5.34 (s, 1H), 4.08 (s, 2H), 3.80 – 3.35 (m, 8H), 2.87 (d, *J* = 4.8 Hz, 3H), 2.35 (s, 1H), 1.49 (d, *J* = 6.7 Hz, 3H). **$^{13}$C NMR** (101 MHz, $CDCl_3$) δ 165.36, 128.32, 128.24, 127.40, 124.16, 115.49, 115.29, 77.38, 77.06, 76.74, 46.14, 45.17, 43.03, 42.64, 42.05, 40.88, 29.70, 27.55, 22.04. **HRMS** (ESI) calcd for $[M+H]^+$ 408.1709, found 408.1709

2-chloro-1-(4-(4-((1-(2-chlorophenyl)ethyl)amino)-6-(methylamino)-1,3,5-triazin-2-yl)piperazin-1-y l)ethan-1-one (**LU5**)

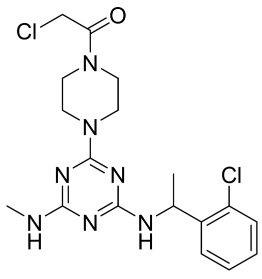

The preparation was described as above procedure to provide compound X by Prep-HPLC as solid ( mg, yield %). **$^{1}$H NMR** (400 MHz, $CDCl_3$) δ 7.38 – 7.28 (m, 2H), 7.23 – 7.08 (m, 2H), 5.44 (s, 1H), 4.07 (s, 2H), 3.88 – 3.26 (m, 8H), 2.88 (s, 3H), 2.55 (s, 1H), 1.45 (d, *J* = 6.9 Hz, 3H). **$^{13}$C NMR** (101 MHz, $CDCl_3$) δ 165.34, 142.80, 132.10, 129.39, 127.78, 127.14, 126.54, 77.38, 77.06, 76.74, 47.88, 46.10, 43.04, 42.67, 42.01, 40.88, 29.70, 27.55, 21.75. **HRMS** (ESI) calcd for $[M+H]^+$ 424.1414, found 424.1416

methyl 2-((4-(4-(2-chloroacetyl)piperazin-1-yl)-6-(methylamino)-1,3,5-triazin-2-yl)amino)-2-(2-chlorophenyl)acetate (**LU6**)

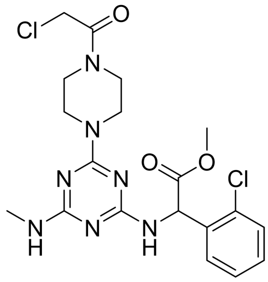

The preparation was described as above procedure to provide compound X by Prep-HPLC as solid ( mg, yield %). **$^{1}$H NMR** (400 MHz, $CDCl_3$) δ 7.44 – 7.33 (m, 2H), 7.23 (dd, *J* = 5.6, 3.6 Hz, 2H), 6.11 (s, 1H), 4.89 (s, 1H), 4.09 (s, 2H), 3.87 – 3.42 (m, 11H), 2.89 (d, *J* = 5.0 Hz, 3H), 2.12 (s, 1H). **$^{13}$C NMR** (101 MHz, $CDCl_3$) δ 165.36, 135.84, 133.85, 129.84, 129.38, 128.63, 127.26, 77.38, 77.06, 76.75, 54.83, 52.86, 46.15, 43.04, 42.66, 42.06, 40.89, 27.60. **HRMS** (ESI) calcd for $[M+H]^+$ 468.1312, found 468.1314

**General procedure for synthesis of copound LU7-LU12**

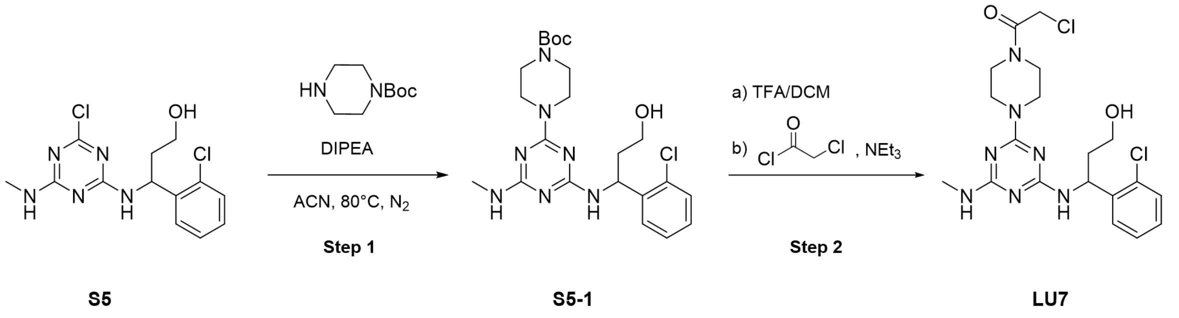


**Step 1:** To a solution of **S5** (150 mg, 0.46 mmol) and tert-butyl piperazine-1-carboxylate (128 mg, 0.69 mmol) in ACN (6 mL) was added DIPEA (178 mg, 1.38 mmol) at 80 °C. The reaction mixture was stirred at $N_2$ atmosphere for 4h. LC-MS indicated that no starting materials remained. After adding cold water, the crude product was precipitated and filtered off. The crude product was washed with acetonitrile (5 mL×3) without further purification to afford **S5-1** (196 mg, yield 90%) . **LRMS** (ESI) calcd for $[M+H]^+$ 478.2, found 478.2

**Step 2:** Compound **S5-1** (196 mg, 0.41 mmol) was subjected to N-Boc deprotection procedure with TFA (2mL) and DCM (4 mL). The reaction mixture was stirred for 2h. LC-MS indicated that no starting materials remained. Without further purification, the crude product was dissolved in 4 mL DCM and then added $NEt_3$ (126 mg, 1.23 mmol), 2-chloroacetyl chloride (46 mg, 0.41 mmol). The reaction mixture was stirred for 4h. The purification by Prep-HPLC afforded the **LU7** (77 mg, yield 41 %) as a white solid. **$^1$H NMR** (400 MHz, DMSO) δ 7.49 (d, $J$ = 7.8 Hz, 1H), 7.37 (d, $J$ = 7.7 Hz, 1H), 7.29 (t, $J$ = 7.5 Hz, 1H), 7.20 (t, $J$ = 7.7 Hz, 1H), 6.91 – 6.54 (m, 1H), 5.61 – 5.39 (m, 1H), 4.59 (s, 1H), 4.41 (d, $J$ = 4.1 Hz, 2H), 3.78 – 3.45 (m, 8H), 2.81 – 2.57 (m, 3H), 1.95 – 1.70 (m, 2H), 1.24 (s, 2H). **$^{13}$C NMR** (101 MHz, $CDCl_3$) δ 165.39, 129.55, 128.26, 127.58, 127.21, 100.00, 77.38, 77.27, 77.06, 76.75, 59.34, 45.98, 42.85, 41.97, 40.85, 29.70, 27.59, 22.69, 14.13. **HRMS** (ESI) calcd for $[M+H]^+$ 454.1520, found 454.1523

2-chloro-1-(4-(4-((1-(2-chlorophenyl)-3-hydroxypropyl)amino)-6-(methylamino)-1,3,5-triazin-2-yl)-2-methylpiperazin-1-yl)ethan-1-one (**LU8**)

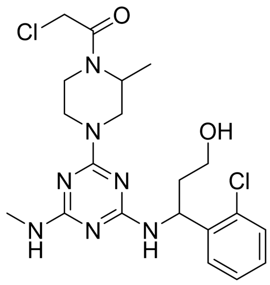

The preparation was described as above procedure to provide compound X by Prep-HPLC as solid ( mg, yield %). **$^1$H NMR** (400 MHz, DMSO) δ 7.49 (d, *J* = 7.9 Hz, 1H), 7.37 (d, *J* = 7.7 Hz, 1H), 7.27 (q, *J* = 6.6 Hz, 1H), 7.23 – 7.15 (m, 1H), 6.58 (s, 1H), 5.59 – 5.26 (m, 1H), 4.74 – 3.96 (m, 6H), 3.50 (d, *J* = 9.3 Hz, 2H), 3.13 – 2.82 (m, 2H), 2.78 – 2.57 (m, 3H), 1.97 – 1.71 (m, 2H), 1.56 – 0.75 (m, 5H). **$^{13}$C NMR** (101 MHz, $CDCl_3$) δ 165.27, 140.83, 132.60, 132.11, 129.60, 128.19, 127.59, 127.17, 95.72, 95.39, 77.38, 77.06, 76.75, 59.23, 49.54, 48.50, 46.73, 46.24, 45.68, 41.18, 29.70, 27.58, 22.23, 16.42, 14.86, 13.88, 12.33. **HRMS** (ESI) calcd for $[M+H]^+$ 468.1676, found 468.1678

N-(2-amino-2-oxoethyl)-1-(2-chloroacetyl)-4-(4-((1-(2-chlorophenyl)-3-hydroxypropyl)amino)-6-(methylamino)-1,3,5-triazin-2-yl)piperazine-2-carboxamide (**LU9**)

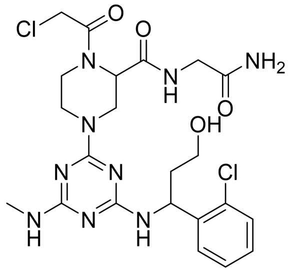

The preparation was described as above procedure to provide compound X by Prep-HPLC as solid ( mg, yield %). **$^1$H NMR** (400 MHz, DMSO) δ 8.27 (s, 1H), 7.61 – 7.01 (m, 7H), 6.57 (s, 1H), 5.48 (s, 1H), 5.14 – 3.99 (m, 7H), 3.90 – 3.42 (m, 6H), 2.81 – 2.57 (m, 3H), 1.94 – 1.69 (m, 2H), 1.39 – 1.21 (m, 2H). **$^{13}$C NMR** (101 MHz, MeOD) δ 172.73, 170.72, 169.88, 168.24, 166.32, 164.42, 141.71, 132.30, 132.06, 129.07, 127.81, 127.30, 126.94, 58.64, 54.91, 49.46, 48.53, 48.32, 48.10, 47.89, 47.68, 47.46, 47.25, 47.04, 42.68, 42.33, 42.11, 41.24, 37.76, 29.35, 26.39. **HRMS** (ESI) calcd for $[M+H]^+$ 554.1792, found 554.1795

N-(3-amino-3-oxopropyl)-1-(2-chloroacetyl)-4-(4-((1-(2-chlorophenyl)-3-hydroxypropyl)amino)-6-(methylamino)-1,3,5-triazin-2-yl)piperazine-2-carboxamide (LU10)

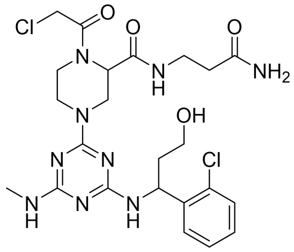


The preparation was described as above procedure to provide compound X by Prep-HPLC as solid ( mg, yield %). 1H NMR (400 MHz, DMSO) δ 8.02 (s, 1H), 7.69 – 7.15 (m, 6H), 6.82 (s, 1H), 5.48 (d, J = 7.2 Hz, 1H), 4.99 – 3.94 (m, 7H), 3.83 – 3.43 (m, 4H), 3.28 – 3.04 (m, 4H), 2.79 – 2.59 (m, 3H), 2.20 (dd, J = 15.7, 7.9 Hz, 2H), 1.95 – 1.67 (m, 2H). 13C NMR (101 MHz, MeOD) δ 174.95, 170.11, 170.06, 169.21, 167.99, 163.94, 132.32, 132.06, 129.11, 127.95, 127.34, 127.03, 58.53, 54.61, 49.55, 48.56, 48.35, 48.14, 47.93, 47.71, 47.50, 47.29, 47.07, 42.55, 42.32, 41.79, 41.38, 37.68, 35.78, 34.52, 26.46. **HRMS** (ESI) calcd for $[M+H]^+$ 568.1949, found 568.1949

1-(2-chloroacetyl)-4-(4-((1-(2-chlorophenyl)-3-hydroxypropyl)amino)-6-(methylamino)-1,3,5-triazin-2-yl)-N-((2-oxopyrrolidin-3-yl)methyl)piperazine-2-carboxamide (**LU11**)

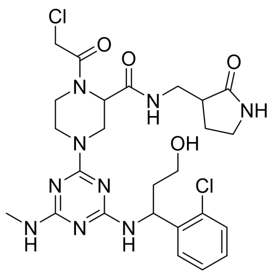


The preparation was described as above procedure to provide compound X by Prep-HPLC as solid ( mg, yield %). 1H NMR (400 MHz, DMSO) δ 7.98 (s, 1H), 7.74 – 7.09 (m, 6H), 6.51 (s, 1H), 5.49 (s, 1H), 5.14 – 3.94 (m, 7H), 3.86 – 3.42 (m, 4H), 3.17 (s, 7H), 2.76 – 2.59 (m, 3H), 2.33 (s, 1H), 2.08 – 1.43 (m, 4H). 13C NMR (101 MHz, MeOD) δ 178.92, 170.30, 167.99, 164.36, 141.86, 132.37, 132.09, 129.08, 127.84, 127.36, 126.90, 58.62, 54.83, 49.47, 48.54, 48.32, 48.11, 47.90, 47.69, 47.47, 47.26, 47.05, 42.46, 42.19, 41.87, 41.33, 40.05, 39.46, 37.72, 26.41, 25.02. **HRMS** (ESI) calcd for $[M+H]^+$ 594.2105, found 594.2106

1-(2-chloroacetyl)-4-(4-(((S)-1-(2-chlorophenyl)-3-hydroxypropyl)amino)-6-(methylamino)-1,3,5-triazin-2-yl)-N-((2-oxopyrrolidin-3-yl)methyl)piperazine-2-carboxamide (**LU12**)

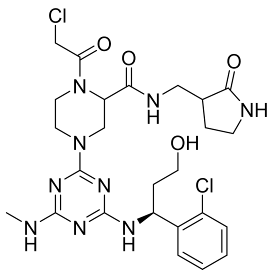

The preparation was described as above procedure to provide compound X by Prep-HPLC as solid ( mg, yield %). [1]**H NMR** (400 MHz, DMSO) δ 7.98 (s, 1H), 7.78 – 7.00 (m, 7H), 6.51 (s, 1H), 5.49 (s, 1H), 5.09 – 4.03 (m, 7H), 3.86 – 3.42 (m, 4H), 3.26 – 2.84 (m, 7H), 2.74 – 2.60 (m, 3H), 2.33 (s, 1H), 2.04 – 1.58 (m, 4H). [13]**C NMR** (101 MHz, MeOD) δ 178.88, 170.26, 167.99, 166.31, 165.52, 164.28, 141.88, 132.09, 129.08, 127.83, 127.36, 126.89, 58.61, 54.83, 49.48, 48.52, 48.31, 48.10, 47.89, 47.67, 47.46, 47.25, 47.03, 42.20, 41.87, 41.31, 40.05, 39.46, 37.76, 26.40, 25.02. **HRMS** (ESI) calcd for $[M+H]^+$ 594.2105, found 594.2103

**General procedure for synthesis of compound LU14**

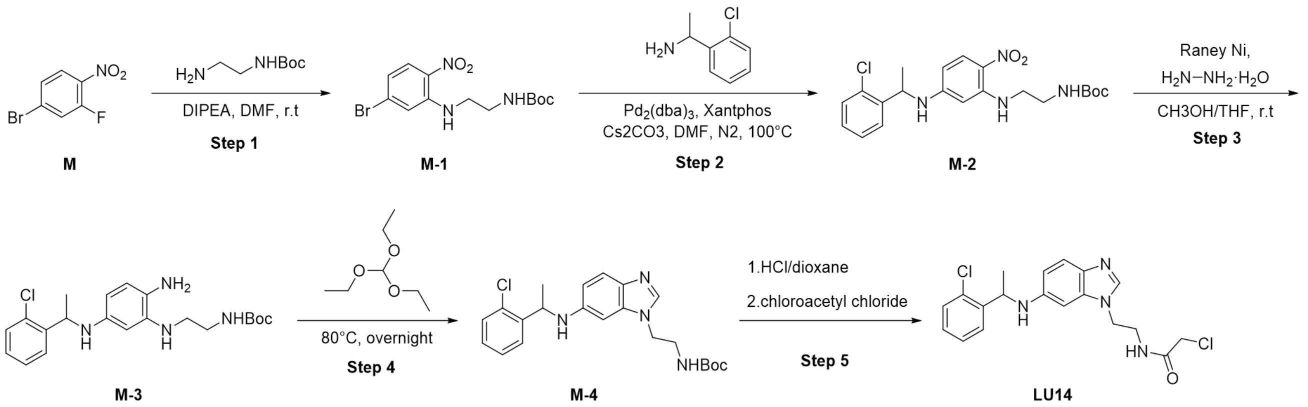


**Step 1:** To a solution of **M** (500mg, 2.27 mmol) and tert-butyl (2-aminoethyl)carbamate (364 mg, 2.27 mmol) in DMF (10 mL) was added DIPEA (881 mg, 6.82 mmol) at room temperature. The reaction mixture was stirred at 25℃ for 4h. LC-MS indicated that no starting materials remained. The purification by flash column chromatography ( Hexenes/EtOAc, 5:1) afforded the **M-1** (780 mg, yield 95%). [1]**H NMR** (400 MHz, DMSO) δ 8.28 (t, *J* = 5.8 Hz, 1H), 7.98 (d, *J* = 9.1 Hz, 1H), 7.33 (d, *J* = 2.1 Hz, 1H), 7.03 (t, *J* = 5.8 Hz, 1H), 6.83 (dd, *J* = 9.1, 2.0 Hz, 1H), 3.41 (q, *J* = 6.0 Hz, 2H), 3.18 (q, *J* = 5.9 Hz, 2H), 1.36 (s, 9H). [13]**C NMR** (101 MHz, DMSO) δ 156.45, 146.23, 131.43, 130.88, 128.54, 118.61, 117.05, 78.32, 42.98, 39.28, 28.63. **LRMS** (ESI) calcd for $[M-56+H]^+$ 304.1, found 304.1

**Step 2:** **M-1** (780 mg, 2.17 mmol) was dissolved in a flask fitted with a rubber septum charged with 1-(2-chlorophenyl)ethan-1-amine (344 mg, 2.21 mmol), $Pd_2(dba)_3$ (297 mg, 0.32 mmol), Xantphos (378 mg, 0.65 mmol), $Cs_2CO_3$ (2.1 g, 6.5 mmol), DMF (10 mL) and then purged with argon. The mixture was stirred at 100 °C overnight. The reaction mixture was then cooled to room temperature, diluted with ethyl acetate (20 mL), filtered through celite, and concentrated in vacuo. The purification by flash column chromatography (Hexanes/EtOAc, 3:1) afforded the **M-2** (395 mg, yield 42%) **$^1$H NMR** (400 MHz, DMSO) δ 8.38 (s, 1H), 7.80 (d, $J$ = 9.5 Hz, 1H), 7.75 (d, $J$ = 6.6 Hz, 1H), 7.44 (ddd, $J$ = 13.6, 7.8, 1.6 Hz, 2H), 7.36 – 7.25 (m, 2H), 6.97 (d, $J$ = 5.2 Hz, 1H), 6.07 (s, 1H), 5.47 (s, 1H), 4.94 (p, $J$ = 6.7 Hz, 1H), 3.23 – 2.97 (m, 4H), 1.47 (d, $J$ = 6.7 Hz, 3H), 1.37 (s, 9H). **$^{13}$C NMR** (101 MHz, DMSO) δ 156.28, 154.08, 148.13, 141.90, 132.01, 129.79, 129.30, 128.66, 128.46, 127.44, 122.89, 78.28, 49.40, 42.53, 38.99, 28.66, 22.32. **LRMS** (ESI) calcd for $[M-56+H]^+$ 379.1, found 379.0

**Step 3:** To a solution of **M-2** (395 mg, 0.91 mmol) in $CH_3OH$/THF (v/v=1/1, 5 mL/5 mL) at $N_2$ atmosphere was added 5%-10% Raney Ni (40 mg) and hydrazine monohydrate (1 mL) at room temperature for 1h. LC-MS indicated that no starting materials remained and then diluted with ethyl acetate (20 mL), filtered through celite, and concentrated in vacuo. The purification by flash column chromatography (Hexanes/EtOAc, 1:1) afforded the **M-3** (323 mg, yield 88%). **$^1$H NMR** (400 MHz, DMSO) δ 7.49 (dd, $J$ = 7.7, 1.8 Hz, 1H), 7.36 (dd, $J$ = 7.8, 1.4 Hz, 1H), 7.27 – 7.15 (m, 2H), 6.88 (t, $J$ = 5.7 Hz, 1H), 6.25 (d, $J$ = 8.0 Hz, 1H), 5.78 (d, $J$ = 2.4 Hz, 1H), 5.53 (dd, $J$ = 8.1, 2.3 Hz, 1H), 5.44 (d, $J$ = 7.6 Hz, 1H), 4.70 (p, $J$ = 6.7 Hz, 1H), 4.27 (t, $J$ = 5.6 Hz, 1H), 3.64 (s, 1H), 3.12 (q, $J$ = 6.2 Hz, 2H), 2.94 (tp, $J$ = 12.3, 5.8 Hz, 2H), 1.40 (s, 9H), 1.35 (d, $J$ = 6.7 Hz, 3H). **$^{13}$C NMR** (101 MHz, DMSO) δ 156.24, 144.22, 141.28, 138.11, 132.25, 129.42, 128.35, 127.85, 127.59, 125.75, 116.56, 101.53, 97.56, 78.16, 49.96, 43.86, 39.38, 28.73, 23.15. **LRMS** (ESI) calcd for $[M+H]^+$ 405.2, found 405.2

**Step 4:** The solution of **M-3** (323 mg, 0.80 mmol) in triethyl orthoformate (4 mL) was stirred at 80°C for 12h. LC-MS indicated that no starting materials remained. The crude product **M-4** was concentrated in vacuo without further purification. **LRMS** (ESI) calcd for $[M+H]^+$ 415.2, found 415.0

**Step 5:** Compound **M-4** (100 mg, 0.24 mmol) was subjected to N-Boc deprotection procedure with HCl (2mL, 4M in dioxane) and dioxane (2 mL). The reaction mixture was stirred for 2h. LC-MS indicated that no starting materials remained. Without further purification, the crude product was dissolved in 4 mL DCM and then added $NEt_3$ (98 mg, 0.97 mmol), 2-chloroacetyl chloride (30 mg, 0.27 mmol). The reaction mixture was stirred for 4h. The purification by Prep-HPLC afforded the **LU14** (72 mg, yield 76 %) as a white solid. **$^1$H NMR** (500 MHz, DMSO) δ 9.24 (s, 1H), 8.44 (t, $J$ = 5.9 Hz, 1H), 7.62 – 7.48 (m, 2H), 7.44 (dd, $J$ = 7.8, 1.5 Hz, 1H), 7.31 – 7.18 (m, 2H), 6.88 (dd, $J$ = 9.0, 2.1 Hz, 1H), 6.60 (d, $J$ = 2.1 Hz, 1H), 4.93 (q, $J$ = 6.6 Hz, 1H), 4.51 – 4.15 (m, 2H), 4.07 – 3.93 (m, 2H), 3.68 – 3.28 (m, 2H), 1.49 (d, $J$ = 6.7 Hz, 3H). **$^{13}$C NMR** (126 MHz, DMSO) δ 167.01, 147.25, 142.26, 139.58, 133.13, 132.57, 129.80, 129.05, 128.18, 127.47, 122.79, 115.80, 115.32, 92.12, 49.63, 46.02, 42.86, 38.18, 22.52. **LRMS** (ESI) calcd for $[M+H]^+$ 391.1, found 390.9

**General procedure for synthesis of compound LU15**

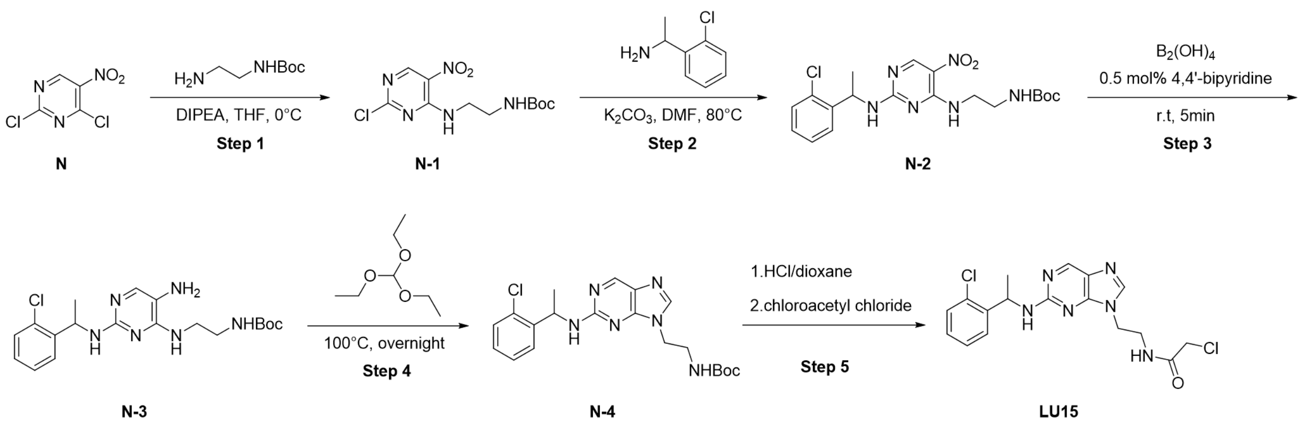


**Step 1:** To a solution of **N** (387mg, 2.0 mmol) and tert-butyl (2-aminoethyl)carbamate (320 mg, 2.0 mmol) in THF (6 mL) was added DIPEA (772 mg, 6.0 mmol). The reaction mixture was stirred at 0°C for 4h. LC-MS indicated that no starting materials remained. The purification by flash column chromatography ( Hexenes/EtOAc, 3:1) afforded the **N-1** (586 mg, yield 93%). **$^1$H NMR** (400 MHz, DMSO) δ 9.13 – 9.05 (m, 1H), 9.03 (s, 1H), 6.95 (t, $J$ = 5.9 Hz, 1H), 3.60 (q, $J$ = 5.9 Hz, 2H), 3.21 (q, $J$ = 5.8 Hz, 2H), 1.35 (s, 9H). **$^{13}$C NMR** (101 MHz, DMSO) δ 162.76, 157.64, 156.32, 155.83, 127.88, 78.24, 41.93, 39.21, 28.62. **LRMS** (ESI) calcd for $[M-H]^+$ 316.1, found 315.9

**Step 2:** Compound **N-1** (586 mg, 1.85 mmol), 1-(2-chlorophenyl)ethan-1-amine (288 mg, 1.85 mmol), $K_2CO_3$ (766 mg, 5.54 mmol) was dissolved in DMF (8 mL). The mixture was stirred at 80 °C for 4h. LC-MS indicated that no starting materials remained. The purification by flash column chromatography (Hexanes/EtOAc, 3:1) afforded the **N-2** (620 mg, yield 77%) **$^1$H NMR** (400 MHz, DMSO) δ 8.86 – 8.62 (m, 1H), 7.54 – 7.20 (m, 4H), 5.71 – 5.35 (m, 1H), 3.47 – 3.37 (m, 1H), 3.26 – 3.17 (m, 1H), 3.14 – 2.78 (m, 2H), 1.55 (d, *J* = 7.0 Hz, 1H), 1.42 (d, *J* = 6.9 Hz, 2H), 1.36 (d, *J* = 9.7 Hz, 9H), 1.29 – 1.16 (m, 2H). **LRMS** (ESI) calcd for $[M+H]^+$ 437.2, found 437.0

**Step 3:** To a solution of **N-2** (216 mg, 0.49 mmol) in DMF (5 mL) was added $B_2(OH)_4$ (133 mg, 1.48 mmol) and 4,4'-bipyridine (39 mg, 0.25 mmol) at room temperature for 5min. LC-MS indicated that no starting materials remained. The purification by flash column chromatography (DCM/MeOH, 20:1) afforded the **N-3** (183 mg, yield 91%). **$^1$H NMR** (400 MHz, DMSO) δ 7.48 (dd, *J* = 7.7, 1.8 Hz, 1H), 7.35 (dd, *J* = 7.8, 1.4 Hz, 1H), 7.25 (td, *J* = 7.5, 1.4 Hz, 1H), 7.22 – 7.14 (m, 2H), 6.85 (s, 1H), 6.73 (t, *J* = 5.5 Hz, 1H), 6.56 (s, 1H), 5.27 (p, *J* = 7.0 Hz, 1H), 3.28 (q, *J* = 5.9 Hz, 2H), 3.13 – 2.82 (m, 2H), 1.38 (s, 9H), 1.34 (d, *J* = 7.0 Hz, 3H). **$^{13}$C NMR** (101 MHz, DMSO) δ 156.11, 155.30, 144.64, 131.95, 129.41, 128.20, 127.71, 127.20, 119.21, 78.10, 48.14, 28.72, 22.08. **LRMS** (ESI) calcd for $[M+H]^+$ 407.2, found 407.0

**Step 4:** The solution of **N-3** (183 mg, 0.45 mmol) in triethyl orthoformate (4 mL) was stirred at 100°C for 12h. LC-MS indicated that no starting materials remained. The purification by flash column chromatography (Hexanes/EtOAc, 1:1) afforded the **N-4** (167 mg, yield 89%). **$^1$H NMR** (400 MHz, MeOD) δ 8.54 (d, *J* = 2.4 Hz, 1H), 7.89 (s, 1H), 7.52 (dd, *J* = 7.6, 1.9 Hz, 1H), 7.38 (dd, *J* = 7.7, 1.5 Hz, 1H), 7.24 – 7.15 (m, 2H), 5.54 (q, *J* = 7.0 Hz, 1H), 4.17 (q, *J* = 7.0 Hz, 2H), 3.45 (dt, *J* = 14.3, 5.4 Hz, 1H), 3.30 – 3.20 (m, 1H), 1.54 (d, *J* = 7.0 Hz, 3H), 1.35 (s, 9H). **$^{13}$C NMR** (101 MHz, MeOD) δ 158.64, 156.77, 153.00, 148.22, 143.12, 132.52, 128.96, 127.57, 126.94, 126.39, 78.86, 48.37, 42.88, 39.15, 27.24, 20.38. **LRMS** (ESI) calcd for $[M+H]^+$ 417.2, found 417.0

**Step 5:** Compound **N-4** (103 mg, 0.33 mmol) was subjected to N-Boc deprotection procedure with HCl (2mL, 4M in dioxane) and dioxane (2 mL). The reaction mixture was stirred for 2h. LC-MS indicated that no starting materials remained. Without further purification, the crude product was dissolved in 4 mL DCM and then added $NEt_3$ (132 mg, 1.32 mmol), 2-chloroacetyl chloride (41 mg, 0.33 mmol). The reaction mixture was stirred for 4h. The purification by Prep-HPLC afforded the **LU15** (69 mg, yield 71 %) as a white solid. **$^1$H NMR** (500 MHz, DMSO) δ 8.85 (s, 2H), 8.43 (s, 1H), 8.31 (s, 1H), 7.56 (dd, *J* = 7.7, 1.7 Hz, 1H), 7.40 (dd, *J* = 7.9, 1.3 Hz, 1H), 7.29 (td, *J* = 7.5, 1.3 Hz, 1H), 7.22 (td, *J* = 7.6, 1.7 Hz, 1H), 5.49 (p, *J* = 6.5 Hz, 1H), 4.17 (td, *J* = 12.4, 7.9 Hz, 2H), 3.98 (s, 2H), 3.66 – 3.25 (m, 2H), 1.48 (d, *J* = 7.0 Hz, 3H). **$^{13}$C NMR** (126 MHz, DMSO) δ 166.85, 132.34, 129.58, 128.79, 127.97, 127.41, 124.97, 119.62, 117.31, 114.99, 112.68, 48.45, 43.42, 42.89, 38.44, 21.66. **LRMS** (ESI) calcd for $[M+H]^+$ 393.1, found 392.9
